# Genome-wide sequence-based genotyping supports a nonhybrid origin of *Castanea alabamensis*

**DOI:** 10.1101/680371

**Authors:** M. Taylor Perkins, Tetyana Zhebentyayeva, Paul H. Sisco, J. Hill Craddock

**Affiliations:** Department of Biology, Geology, and Environmental Science, The University of Tennessee at Chattanooga, Chattanooga, TN 37403, USA; Department of Ecosystem Science and Management, The Pennsylvania State University, University Park, PA 16802, USA; The American Chestnut Foundation, Carolinas Chapter, Asheville, NC 28804, USA

**Keywords:** American chestnut, *Castanea dentata*, *Castanea pumila*, chinquapin, genotyping-by-sequencing, hybridization, Fagaceae

## Abstract

The genus *Castanea* in North America contains multiple tree and shrub taxa of conservation concern. The two species within the group, American chestnut (*Castanea dentata*) and chinquapin (*C. pumila* sensu lato), display remarkable morphological diversity across their distributions in the eastern United States and southern Ontario. Previous investigators have hypothesized that hybridization between *C. dentata* and *C. pumila* has played an important role in generating morphological variation in wild populations. A putative hybrid taxon, *Castanea alabamensis*, was identified in northern Alabama in the early 20th century; however, the question of its hybridity has been unresolved. We tested the hypothesized hybrid origin of *C. alabamensis* using genome-wide sequence-based genotyping of *C. alabamensis*, all currently recognized North American *Castanea* taxa, and two Asian *Castanea* species at >100,000 single-nucleotide polymorphism (SNP) loci. With these data, we generated a high-resolution phylogeny, tested for admixture among taxa, and analyzed population genetic structure of the study taxa. Bayesian clustering and principal components analysis provided no evidence of admixture between *C. dentata* and *C. pumila* in *C. alabamensis* genomes. Phylogenetic analysis of genome-wide SNP data indicated that *C. alabamensis* forms a distinct group within *C. pumila* sensu lato. Our results are consistent with the model of a nonhybrid origin for *C. alabamensis*. Our finding of *C. alabamensis* as a genetically and morphologically distinct group within the North American chinquapin complex provides further impetus for the study and conservation of the North American *Castanea* species.

---

Hybridization is an important and widespread phenomenon in plants (Ellstrand et al., 1996; Mallet, 2005; Soltis and Soltis, 2009; Whitney et al., 2010). Evolutionary outcomes of hybridization can include reinforcement or breakdown of reproductive barriers, increased intraspecific diversity, transfer of genetic adaptations between species, and the origin of new ecotypes or species (Rieseberg, 1997). Many authors have asserted that the descendants of natural interspecific hybridization warrant conservation because they represent a natural part of the evolutionary legacy of taxa (Whitham et al., 1991; Allendorf et al., 2001; Allendorf et al., 2013; Stronen and Paquet, 2013; Jackiw et al., 2015). Thus, the identification and characterization of natural hybrids, particularly in groups of conservation concern, is a priority for understanding and protecting plant biodiversity. The genus *Castanea* Mill. (Fagaceae) in North America exemplifies this concept.

There exists remarkable morphological diversity within the North American *Castanea*, a group that is thought to be comprised of just two species, American chestnut (*Castanea dentata* (Marsh.) Borkh.) and North American chinquapin (*C. pumila* (L.) Mill.) (Johnson, 1988). Within *C. pumila* alone, plant habit at maturity varies from rhizomatous subshrub in some populations of the Gulf Coastal Plain to canopy tree in other populations (Johnson, 1988; Nixon, 1997). Such variation in plant habit, as well as variation in flower, fruit, leaf, and twig morphology has led several authors to speculate that hybridization between *C. dentata* and *C. pumila* has contributed significantly to the variation observed in wild North American *Castanea* populations (Dode, 1908; Camus, 1929; Elias, 1971; Little, 1979; Nixon, 1997; Kubisiak and Roberds, 2006; Binkley, 2008; Shaw et al., 2012; Li and Dane, 2013). Reports of naturally occurring hybrids between *C. dentata* and *C. pumila* var. *pumila* have resulted in the description of two hybrid taxa. Dode (1908) published the first description of putative hybrid progeny of *C. dentata* and *C. pumila*, referring to these plants from North Carolina, Virginia, and Maryland as *C.* × *neglecta* Dode. The second hybrid taxon resulted from Camus’ (1929) revised treatment of the nonhybrid species *C. alabamensis* Ashe, a tree endemic to the Appalachian Mountains of northern Alabama. The putative hybrid taxa, *C. alabamensis* and *C.* × *neglecta*, both possess a combination of traits typically used to differentiate *C. dentata* from *C. pumila*. In *C. alabamensis*, specifically, plants produce one pistillate flower per cupule and one nut per bur, like *C. pumila*, yet mature plants grow as medium to large trees, have glabrous twigs, and have apparently glabrous abaxial leaf surfaces when viewed without magnification, like *C. dentata* with which they are sympatric (Ashe, 1925).

Multiple genetic studies over the past decade have investigated the putative hybridity of several North American *Castanea* populations (Binkley, 2008; Dane, 2009; Shaw et al., 2012; Li and Dane, 2013). Binkley (2008) and Shaw et al. (2012) used a small number of noncoding cpDNA regions to investigate populations of suspected hybrids in northwestern Georgia, which they described as having an “intermediate morphology” that corresponded to the hybrid taxon *C.* × *neglecta*. Their *C.* × *neglecta* accessions had a unique cpDNA haplotype that was not found in a diverse panel of the putative parental species, *C. dentata* and *C. pumila*, contrary what they predicted would have been observed in hybrid individuals. Authors of both studies concluded that further investigation would be needed to rigorously test the hybridity of their study populations in northwestern Georgia (Binkley, 2008; Shaw et al., 2012). The most recent study of reputed naturally-occurring hybrids between *C. dentata* and *C. pumila* used six noncoding cpDNA regions and 680 bp from two nuclear DNA regions to investigate the extent of hybridization between these two species (Li and Dane, 2013). An important finding of Li and Dane (2013) related to their discovery of two different morphological groups at a site in northern Alabama, which they designated as distinct morphological types of *C. dentata*: Type I *C. dentata* (possessing morphological features typical of *C. dentata* throughout its range) and Type II *C. dentata* (possessing several leaf trichome features that the authors described as indicative of morphological intermediacy between *C. dentata* and *C. pumila*). In phylogenetic analyses of cpDNA and nuclear DNA sequences, Type I *C. dentata* grouped with other *C. dentata* plants sampled throughout the species’ range, while Type II *C. dentata* grouped with *C. pumila* in analyses of both chloroplast and nuclear DNA datasets. Li and Dane (2013) concluded that the genetic and morphological patterns observed in Type II *C. dentata* might be the result of past hybridization between *C. dentata* and *C. pumila*.

A limitation of previous genetic studies of putative hybrids between *C. dentata* and *C. pumila* is that nearly all of the analyses relied principally on datasets comprised of information from a small number of maternally-inherited cpDNA loci. In the few cases where loci from the nuclear genome were analyzed, these data were not tested for signatures of hybridization. Although, Li and Dane (2013) used both cpDNA and nuclear DNA, results of their analysis of 680 bp from two nuclear loci showed no genetic evidence of *C. dentata* ancestry in their Type II *C. dentata* individuals. Their dataset also lacked the information necessary to more finely discern relationships between different putatively hybridized Type II *C. dentata* individuals and the *C. pumila* individuals found on the same branch of the phylogenetic tree. Moreover, analytical methods used to make inferences about past hybridization in this group have been limited to phylogenetic trees and cpDNA haplotype networks, using models that assume a strictly bifurcating pattern of lineage evolution (Binkley, 2008; Dane, 2009; Shaw et al., 2012; Li and Dane, 2013). Therefore, despite much speculation, convincing evidence regarding the occurrence or extent of natural hybridization between the North American *Castanea* taxa has not been produced.

The North American *Castanea* are currently thought to consist of either two or three species. The most recent formal taxonomic revision of the group relied on morphology and concluded that North American *Castanea* is comprised of two species, American chestnut (*C. dentata*) and North American chinquapin (*Castanea pumila*)—the latter species containing the botanical varieties Allegheny chinquapin (*C. pumila* (L.) Mill. var. *pumila*) and Ozark chinquapin (*C. pumila* var. *ozarkensis* (Ashe) Tucker) (Johnson, 1988). Other taxonomists contend that the ecological and morphological differences between Allegheny chinquapin and Ozark chinquapin are substantial enough to justify recognition at the level of species, and they recognize three species of North American *Castanea*: *C. dentata*, *C. pumila*, and *C. ozarkensis* (Nixon, 1997; Weakley, 2015). American chestnut (*C. dentata*) produces three pistils per cupule, three nuts per bur, and apparently glabrous abaxial leaf surfaces that rarely contain stellate trichomes, and healthy individuals grow as trees at maturity (Nixon, 1997). Ozark chinquapin and Allegheny chinquapin typically produce one pistil per cupule, one nut per bur, and have sun leaves with abaxial surfaces that are covered in stellate trichomes (Johnson, 1988; Nixon, 1997). However, Ozark chinquapin grows as a tree at maturity and Allegheny chinquapin varies from rhizomatous shrub, to non-rhizomatous shrub, to small tree at maturity (Johnson, 1988). Dode (1908) describes the hybrid taxon *C.*× *neglecta* as having slightly pubescent twigs and leaves, reminiscent of the putative parent *C. pumila*, and larger burs and arborescent habit, like the putative parent *C. dentata*. Ashe (1925) originally described *C. alabamensis* as a distinct chinquapin species, but it was treated as a hybrid taxon throughout most of the 20^th^ century (Camus, 1929; Elias, 1971; Little, 1979). In the most recent taxonomic revision of the group, Johnson (1988) found no living plants that corresponded to the description *C.* × *neglecta*. Johnson (1988) treated *C. alabamensis* as a synonym of *C. pumila* var. *ozarkensis* on the basis of morphological similarities between the two taxa and, after field work near Ashe’s *C. alabamensis* type locality in northern Alabama, he concluded that *C. pumila* var. *ozarkensis* had been extirpated from east of the Mississippi River by chestnut blight.

The advent of affordable genome-wide sequence-based genotyping methods has recently allowed new insights into questions of hybridization and admixture in plants (Escudero et al., 2014; Eaton et al., 2015; Baute et al., 2016; Owens et al., 2016; Leroy et al., 2017; McVay et al., 2017; Zhao et al., 2018; Turner et al., 2018). The availability of large numbers of markers distributed across the genome has allowed researchers to address questions that were previously intractable with less comprehensive datasets. Notably, these methods have allowed researchers to understand hybridization, introgression, and species boundaries on a fine scale in *Quercus*, a genus closely related to *Castanea* that is well known for a history of interspecific hybridization (Eaton et al., 2015; Leroy et al., 2017; McVay et al., 2017). Thus, genome-wide sequence-based genotyping represents a potentially useful approach for understanding hybridization and admixture in the North American *Castanea* species.

Here, we investigate the origins of purported hybrid populations of *Castanea* in the eastern United States. Our main objective was to test the hypothesized hybrid origin of samples corresponding to the morphological description of *C. alabamensis*. A secondary objective was to test for admixture in sympatric populations of *C. dentata* and *C. pumila*. We used sequence-based genotyping to gather genome-wide single nucleotide polymorphism (SNP) data from samples representing *C. alabamensis*, all currently recognized North American *Castanea* taxa (*C. dentata*, *C. pumila* var. *pumila*, *C. pumila* var. *ozarkensis*) and two East Asian outgroup species (*C. crenata* and *C. mollissima*). Additionally, we examined the morphology of 865 herbarium accessions representing the type specimens of *C. alabamensis* and all extant *Castanea* species to place our samples in taxonomic context with recent studies of hybridization in the North American *Castanea* (Binkley, 2008; Shaw et al., 2012; Li and Dane, 2013) and earlier hypotheses based on morphology (Ashe, 1925; Camus, 1929; Elias, 1971; Little, 1979; Johnson, 1988).

## MATERIALS AND METHODS

### Plant materials, DNA extraction, and morphological comparisons

Young leaf tissues for DNA extraction were collected from 106 naturally-occurring North American *Castanea* and cultivated East Asian *Castanea* plants. We sampled plants corresponding to Ashe’s (1925) description of *C. alabamensis*, the putative parent taxa, *C. dentata* and *C. pumila* var. *pumila*, the remaining North American taxon, *C. pumila* var. *ozarkensis*, and two East Asian outgroup species, *C. mollissima* Blume (Chinese chestnut) and *C. crenata* Siebold & Zucc. (Japanese chestnut) (Table 1; Appendix S1). Samples representing two of the *C.* × *neglecta* populations studied by Binkley (2008) and Shaw et al. (2012) were included in sequencing (Table 1; Appendix S1). Total genomic DNA was extracted from 100 mg of fresh or frozen leaf tissue using the Qiagen DNeasy Plant Mini Kit (Qiagen, Valencia, California, USA) or a modified version of the CTAB protocol of Kubisiak et al. (2013). DNA quality and integrity were examined using a NanoDrop ND-8000 spectrophotometer (Thermo Fisher Scientific Inc., Waltham, Massachusetts, USA) and electrophoresis on 1% agarose gels. Genomic DNA was quantified using the Qubit 2.0 fluorometric quantification assay (Life Technologies, Carlsbad, California, USA).

**Table 1.** Morphological and locality information for Castanea populations genotyped in this study. Data for numbers of pistils per cupule and nuts per bur reflect observations from spring and fall, respectively.

| Taxon/County, State | Site name | No. of plants | GBS IDs | Morphological features |  |  |  |  |  |
| --- | --- | --- | --- | --- | --- | --- | --- | --- | --- |
|  |  |  |  | Abaxial with abundant stellate trichomes | Simple trichomes on laminar surfaces | Glaucous abaxial surface | Ciliate leaf margin | Pubescent twigs | Pistils per cupule/ nuts per bur |
| ***C. pumila* var. *pumila*** |  |  |  | **Yes** | **Yes** | **No** | **Yes** | **Yes** | **1 (-3)** |
| Oconee, SC | BF | 4 | 29, 33, 37, 42 | Yes | Yes | No | Yes | Yes | NA |
| Oconee, SC | CR | 5 | 15, 19, 21, 25, 30 | Yes | Yes | No | Yes | Yes | NA |
| Cherokee, NC | SE | 8 | 53, 54, 55, 57, 58, 60, 61, 62 | Yes | Yes | No | Yes | Yes | 1 |
| Charleston, SC | AW | 9 | 1, 5, 9, 13, 17, 23, 28, 32, 36 | Yes | Yes | No | Yes | Yes | NA |
| Sumter, SC | WB | 2 | 41, 44 | Yes | Yes | No | Yes | Yes | NA |
| Suwannee, FL | SS | 3 | 80, 81, 82 | Yes | Yes | No | No | No | 1,2,3 |
| Transylvania, NC | GO | 2 | 20, 24 | Yes | Yes | No | Yes | Yes | NA |
| Baker, GA | JR | 1 | 26 | Yes | Yes | No | Yes | Yes | 1 |
| Pickens, GA | TM | 6 | 35, 40, 43, 46, 47, 48 | Yes | Yes | No | Yes | Yes | 1 |
| ***C. pumila* var. *ozarkensis*** |  |  |  | **Yes** | **Yes** | **No** | **Yes** | **No** | **1** |
| Lawrence, MO | MO | 1 | 63 | Yes | Yes | No | Yes | No | 1 |
| Sharp, AR | SH | 1 | 64 | Yes | Yes | No | Yes | No | 1 |
| ***C. alabamensis*** |  |  |  | **No** | **Yes** | **Yes** | **Yes** | **No** | **1** |
| Cleburne, AL | HU | 3 | 67, 86, 89 | No | Yes | Yes | Yes | No | 1 |
| Clay, AL | AG | 5 | 73, 75, 76, 85, 87 | No | Yes | Yes | Yes | No | 1 |
| Calhoun, AL | CH | 6 | 90, 92, 93, 94, 95, 96 | No | Yes | Yes | Yes | No | 1 |
| Floyd, GA | JM<sup>†</sup> | 1 | 22 | No | Yes | Yes | Yes | No | 1 |
| Floyd, GA | PO<sup>†</sup> | 1 | 38 | No | Yes | Yes | Yes | No | 1 |
| ***C. dentata*** |  |  |  | **No** | **No** | **No** | **No** | **No** | **3** |
| Cleburne, AL | HU | 3 | 68, 69, 70 | No | No | No | No | No | NA |
| Clay, AL | AG | 2 | 74, 77 | No | No | No | No | No | NA |
| Calhoun, AL | CH | 1 | 91 | No | No | No | No | No | NA |
| Cleburne, AL | FR | 3 | 71, 72, 84 | No | No | No | No | No | NA |
| Morgan, AL | LA | 3 | 78, 79, 88 | No | No | No | No | No | NA |
| Pickens, GA | TM | 2 | 27, 31 | No | No | No | No | No | NA |
| Lincoln, TN | LI | 1 | 83 | No | No | No | No | No | 3 |
| Oconee, SC | BF | 2 | 3, 45 | No | No | No | No | No | NA |
| Oconee, SC | CR | 2 | 7, 11 | No | No | No | No | No | NA |
| Pickens, SC | CE | 4 | 2, 6, 10, 14 | No | No | No | No | No | NA |
| Transylvania, NC | GO | 1 | 18 | No | No | No | No | No | NA |
| Cherokee, NC | JB | 3 | 50, 51, 52 | No | No | No | No | No | NA |
| Cherokee, NC | SE | 1 | 59 | No | No | No | No | No | NA |
| Buncombe, NC | DO | 1 | 102 | No | No | No | No | No | 3 |
| Avery, NC | CB | 1 | 103 | No | No | No | No | No | 3 |
| Washington, VA | WN | 1 | 97 | No | No | No | No | No | 3 |
| Adair, KY | KY | 1 | 16 | NA | NA | NA | NA | NA | NA |
| Westmoreland, PA | WE | 1 | 12 | No | No | No | No | No | NA |
| Warren, PA | WR | 1 | 8 | No | No | No | No | No | NA |
| Huntingdon, PA | RO | 1 | 4 | No | No | No | No | No | NA |
| Knox, ME | KN | 1 | 34 | No | No | No | No | No | NA |
| Piscataquis, ME | AT | 1 | 39 | No | No | No | No | No | NA |
| Piscataquis, ME | SB | 1 | 66 | No | No | No | No | No | NA |
| Waldo, ME | WA | 1 | 65 | No | No | No | No | No | NA |
| ***C. mollissima*** |  |  |  | **NA** | **NA** | **NA** | **NA** | **NA** | **3** |
| ‘Nanking’ | NK | 1 | 100 | NA | NA | NA | NA | NA | 3 |
| ‘Mahogany’ | MY | 1 | 98 | NA | NA | NA | NA | NA | 3 |
| ***C. crenata*** |  |  |  | **NA** | **NA** | **NA** | **NA** | **NA** | **3** |
| ‘Fort Defiance’ | FD | 1 | 104 | NA | NA | NA | NA | NA | 3 |
| ‘Morrow Mountain’ | MM | 1 | 107 | NA | NA | NA | NA | NA | 3 |
| ‘Rita 1’ | RI | 1 | 108 | NA | NA | NA | NA | NA | NA |
| ‘Rita 2’ | RA | 1 | 110 | NA | NA | NA | NA | NA | NA |

Note: NA = character was not observed in the individual or species; † = samples 22 and 38, from the JM and PO sites, were previously identified as *C. × neglecta* and sequenced in the studies of Binkley (2008) and Shaw et al. (2012).

Corresponding herbarium vouchers were obtained for wild-collected North American *Castanea* samples and assessed for a suite of morphological features that differentiate *C. dentata*, *C. pumila* var. *pumila*, *C. pumila* var. *ozarkensis*, and *C. alabamensis* (Table 1). Herbarium vouchers were also collected from the northern Alabama population containing the suspected hybrid “Type II *C. dentata*” studied by Li and Dane (2013) (Appendix S2). Finally, morphological comparisons were made between the genotyped plants and the entire *Castanea* collection at the University of North Carolina Chapel Hill Herbarium (NCU), which contains a total of 865 specimens representing all extant *Castanea* species and the type specimens of *C. alabamensis* (Appendix S3). Morphological features assessed included four leaf characters (sun leaf abaxial surface covered with stellate trichomes or not; simple trichomes present or absent on abaxial lamina; abaxial surface glaucous or not; leaf margin ciliate or eciliate), one twig character (twigs pubescent or not), and one flower/fruit character (pistil per cupule ratio or nut per bur ratio).

### GBS library construction, data processing, and SNP discovery

Genotyping-by-sequencing (GBS) libraries were prepared using a modified version of the methods of Elshire et al. (2011), as described by Zhebentyayeva et al. (2019). We followed all aspects of the GBS library preparation protocol of Zhebentyayeva et al. (2019), with one exception—we double-digested DNA samples with *Pst*I and *Mse*I. Briefly, we performed double digestion of DNA samples, ligated fragments to Illumina sequencing adapters and custom barcoded adapters (see Appendix S4 for barcode sequences), pooled samples, then purified the pooled samples using a QIAquick PCR purification kit (Qiagen, Valencia, California, USA). Pooled samples were amplified using 18 cycles of PCR and size-selection was performed using 0.4× and 0.8× volumes of Mag-Bind Total Pure NGS magnetic beads (Omega Bio-Tek, Georgia, USA) to remove fragments larger than 1.5 kb and smaller than 121 bp, respectively. The quality of GBS libraries was validated using a 2100 Bioanalyzer (Agilent Genomics, Santa Clara, California, USA). 96-plex GBS libraries were paired-end sequenced (2 × 125 bp) on an Illumina HiSeq 2500 (Illumina Inc., San Diego, California, USA) at the Hollings Cancer Center at the Medical University of South Carolina.

Data processing of Illumina reads was performed using Clemson University’s Palmetto Cluster high-performance computing resource as described by Zhebentyayeva et al. (2019). Default settings of the Stacks 1.45 program ‘process_radtags’ (Catchen et al., 2011; Catchen et al., 2013; Rochette and Catchen, 2017) were used to demultiplex raw Illumina reads according to barcodes, discard reads with uncalled bases, discard reads with low quality scores, and filter reads for the presence of *Pst*I and *Mse*I restriction sites. Demultiplexed reads were uploaded to the National Center for Biotechnology Information (NCBI) Sequence Read Archive (SRA) under BioProject ID: PRJNA541592. Individual plants with less than 100,000 total reads were removed from further processing. Samples from the *C. mollissima* and *C. crenata* outgroup cultivars were replicated twice in GBS libraries, and technical replicates with the highest number of retained reads were used for further processing. Demultiplexed reads were aligned to the *C. mollissima* reference genome v.1.1 (https://www.hardwoodgenomics.org) using the GSNAP software package (Wu and Nacu, 2010). Single nucleotide polymorphisms were called against the *C. mollissima* reference genome sequence and a catalog of tags and SNPs was generated using the ‘ref_map.pl’ command in Stacks, with default settings. SNP genotypes were generated in the ‘populations’ program in Stacks and filtered to remove indels and multi-nucleotide variants, leaving only bi-allelic SNPs and invariant sites. To avoid issues resulting from analysis of tightly linked markers, we used the *whitelist* feature in the ‘populations’ program to retain only one SNP per locus and create a list of 500,000 randomly selected SNPs for further processing. Individual genotypes were filtered to retain only SNPs supported by five or more reads and SNPs with a minimum allele frequency of 0.01. The SNPs were further filtered to produce three different datasets for analysis, two of which were exported in Variant Call Format (VCF) and the third as a .txt file: (1) a dataset comprised of 103,616 SNPs and 103 individuals representing North American and east Asian *Castanea* species was produced by filtering for SNPs present in >80% of individuals (dataset ‘NAC+EAC’); a dataset comprised of 190,656 SNPs and 96 individuals representing the North American *Castanea* taxa was produced by filtering for SNPs present in >75% of individuals (dataset ‘NAC’); and a smaller dataset comprised of 583 SNPs and 103 individuals representing North American and east Asian *Castanea* species for STRUCTURE analysis was produced by filtering for SNPs present in >95% of individuals (dataset ‘NAC+EAC583’). All datasets exported by Stacks were deposited on GitHub: https://github.com/MTPerkins/Nonhybrid_origin_of_Castanea_alabamensis (see Appendix S5 for correspondence of dataset names to files on GitHub).

### Phylogeny and genetic differentiation

We inferred a maximum likelihood phylogenetic tree for the North American and East Asian *Castanea* taxa (dataset ‘NAC+EAC’) using the RAxML 8.2.11 (Stamatakis 2014) plugin for Geneious 11.1.5 (http://www.geneious.com/). We used the GTRCAT model of nucleotide evolution with 100 rapid bootstrap replicates and a subsequent search for the best scoring maximum likelihood tree.

To compare levels of genetic differentiation between the groups identified in the phylogenetic tree, we estimated Weir and Cockerham’s *FST* (1984) between pairs of taxa in the ‘NAC+EAC’ dataset using the ‘SNPRelate’ package (Zheng et al. 2012) in R (R Core Team 2018), with default settings.

### STRUCTURE analyses

To determine the number of genetically differentiated clusters (*K*) and to detect admixture among all samples (i.e., North American and East Asian species), we performed Bayesian clustering analysis on the ‘NAC+EAC583’ dataset with STRUCTURE 2.3.4 software (Pritchard et al. 2000; Falush et al. 2003). We ran STRUCTURE using 35,000 burn-in repetitions, 35,000 Markov Chain Monte Carlo repetitions after burn-in, the correlated allele frequencies setting, and the admixture model for 10 iterations at *K* = 1-9 (i.e., the number of putative taxa, plus three). After finding that *C. alabamensis* grouped with *C. pumila* sensu lato in phylogenetic analysis and the first STRUCTURE analysis, we determined the number of genetic clusters and levels of admixture in this group by removing *C. dentata*, *C. crenata*, and *C. mollissima* from the ‘NAC+EAC583’ dataset and running STRUCTURE with the same settings as above, apart from using *K* = 1-16 (i.e., the number of chinquapin sample sites, plus one). The number of clusters within our *Castanea* samples from North America and eastern Asia and within the chinquapins were determined by the Δ*K* method (Evanno et al. 2005) implemented within the STRUCTURE HARVESTER program (Earl and vonHoldt 2012). Clustering analysis output was summarized and visualized using the Cluster Markov Packager Across K algorithm (CLUMPAK), with default settings (Kopelman et al. 2015). Because the STRUCTURE analysis of the full dataset identified one putative *C. dentata* sample and one putative *C. pumila* var. *ozarkensis* sample as admixed with East Asian *Castanea* spp., we did not include these two samples in further analysis.

### Principal components analysis

To understand the partitioning of genetic variation within the North American *Castanea* species and determine the relative placement of *C. alabamensis* samples, we performed principal components analysis (PCA) using the ‘NAC’ dataset. Principal components analysis was performed using the ‘SNPRelate’ package in R, with settings “remove.monosnp=TRUE”, “maf=NaN”, and “missing.rate=NaN”. Because missing genotypes as a result of variation in DNA quality can introduce apparent, but erroneous, population structure in PCA (Patterson et al., 2006), the dataset was further filtered for PCA by identifying individuals with <30% missing SNP loci using TASSEL v5.0 (Bradbury et al. 2007) and specifying for analysis only individuals below this threshold.

## RESULTS

### Morphological comparisons

Our collections from northern Alabama *Castanea* populations yielded samples of *C. dentata* and several samples that corresponded to the taxonomic description of *C. alabamensis* (Ashe, 1925). Plants identified as *Castanea alabamensis* differed from all other North American *Castanea* taxa in certain key aspects (Table 1; Fig. 1).

**Fig. 1.**
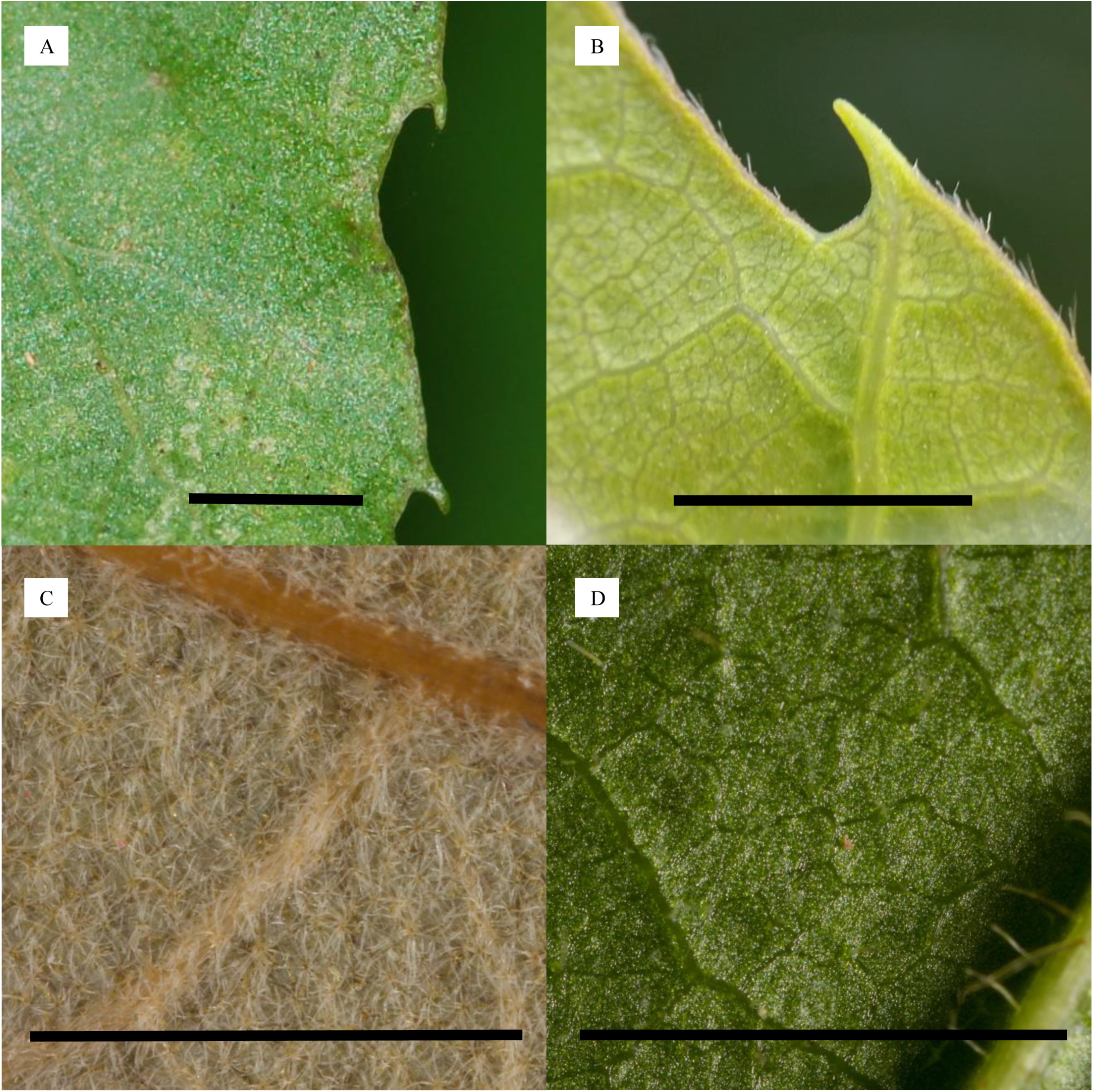
Morphological features distinguishing *C. alabamensis* from *C. dentata* and *C. pumila* var. *ozarkensis*. (A) Eciliate leaf margin of *C. dentata*. (B) Ciliate leaf margin of *C. alabamensis*. (C) Abaxial leaf surface of *C. pumila* var. *ozarkensis* densely covered with stellate trichomes. (D) Abaxial leaf surface of *C. alabamensis* lacking stellate trichomes, but with occasional simple trichomes on laminar surface. Simple trichomes projecting from lateral vein are visible in the lower righthand corner of the image. Scale bars = 5 mm.

While the prevalence of chestnut blight infection prevented us from gathering data on floral/fruit morphology in many wild populations (i.e., plants were typically not reproductively mature), the nut per bur and pistil per bur ratios observed were consistent with the most recent taxonomic treatments of the group (Johnson, 1988; Nixon, 1997; Weakley, 2015). Chestnut blight infection also prevented us from assessing mature plant habit in most North American *Castanea* populations; however, we did observe a few reproductively mature *C. alabamensis* individuals where we could determine that plant habit of mature, healthy trees is typically that of a single-stemmed tree, rather than a shrub, consistent with Ashe’s description of the species (1925).

Morphological study of the entire *Castanea* collection at NCU, the voucher specimens from Binkley (2008) and Shaw et al. (2012), and recent collections from the Ruffner Mtn., AL, site studied by Li and Dane (2013) revealed that the *C.* × *neglecta* populations studied by Binkley (2008) and Shaw et al. (2012) and the Type II *C. dentata* studied by Li and Dane (2013) are morphologically identical to *C. alabamensis* (Ashe, 1925) (Appendix S2). The *C.* × *neglecta* samples of Binkley (2008) and Shaw et al. (2012) and the Type II *C. dentata* samples of Li and Dane (2013) are identical to Ashe’s (1925) original collections of *C. alabamensis* in all aspects of leaf, twig, and flower/fruit morphology that we assessed.

### Illumina sequencing and SNP discovery

A total of 489.9 million Illumina reads were obtained for 106 plants (including both technical replicates of the East Asian cultivars) (Appendix S4). Three plants, Haun *C. dentata*, SE56 *C. pumila* var. *pumila*, and TM49 *C. pumila* var. *pumila*, had <100,000 retained reads and were excluded from further processing. Only the higher quality technical replicate of each *C. mollissima* and *C. crenata* cultivar was retained. After removal of the three failed samples and four lower quality technical replicates of Asian *Castanea* samples, the average clean reads per individual was 4.384 million. In the 103 samples that were retained for genotyping, average coverage depth was 44×. Three datasets were produced for analysis by Stacks 1.45: dataset ‘NAC+EAC’ contained 103 individuals of North American and eastern Asian *Castanea* species and 103,616 SNPs; dataset ‘NAC’ contained 96 individuals of North American *Castanea* species and 190,656 SNPs; and dataset ‘NAC+EAC583’ contained 103 individuals of North American and eastern Asian *Castanea* species and 583 SNPs.

### Phylogeny and genetic differentiation

The maximum likelihood phylogeny inferred from the ‘NAC+EAC’ dataset indicated the existence of six distinct groups corresponding to the morphologically-defined taxa present in our dataset: *C. mollissima*, *C. crenata*, *C. dentata*, *C. pumila* var. *ozarkensis*, *C. pumila* var. *pumila*, and *C. alabamensis* (Fig. 2). The North American *Castanea* species formed a monophyletic group. Two main clades were present within the North American *Castanea* group: (1) *C. dentata* and (2) the North American chinquapins, inclusive of *C. alabamensis*, *C. pumila* var. *pumila*, and *C. pumila* var. *ozarkensis*. Bootstrap values for the nodes separating named taxa were 100% in all cases, except the node separating *C. alabamensis* and *C. pumila* var. *pumila*, which was 91% (Fig. 2; Appendix S6).

**Fig. 2.**
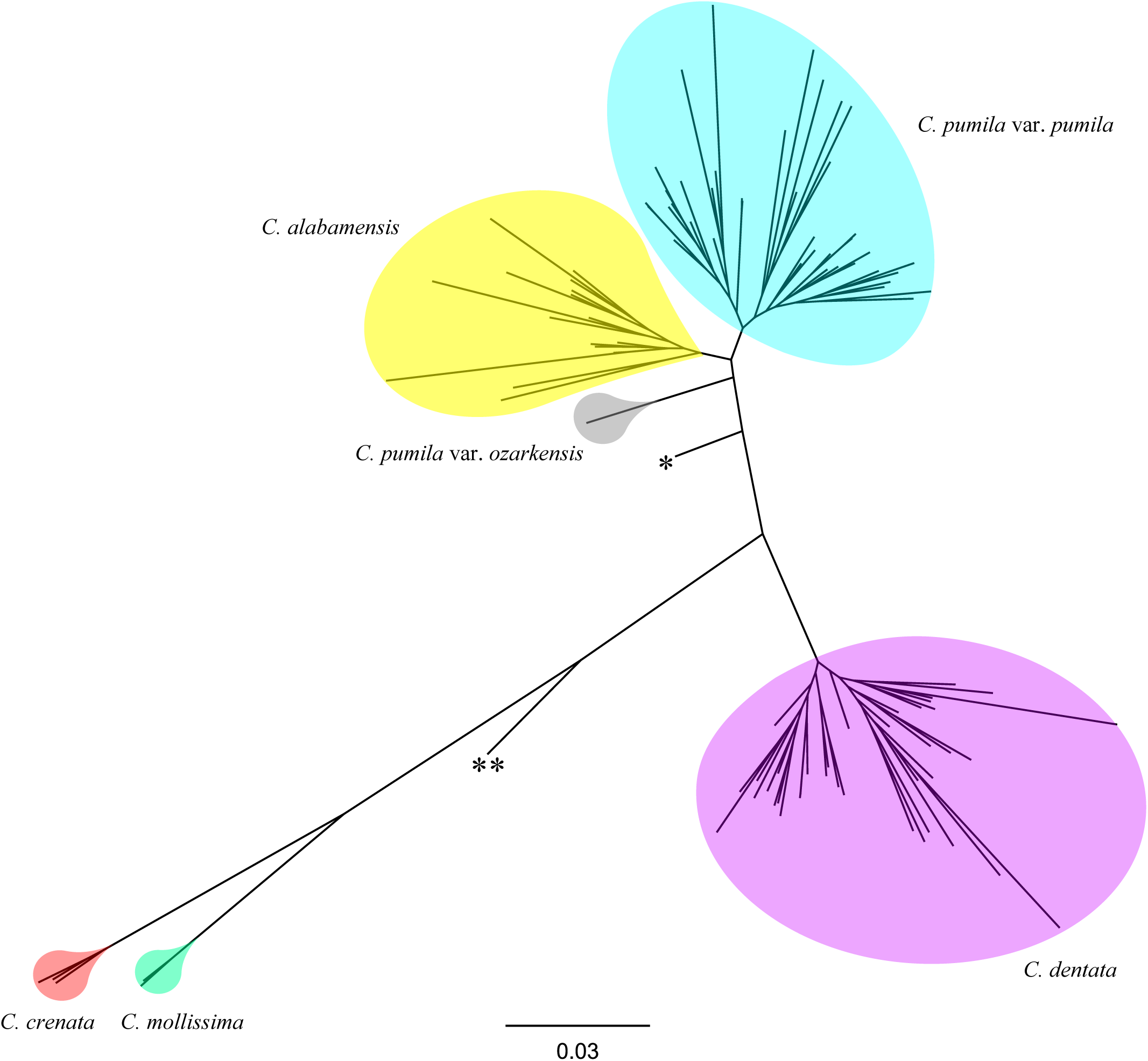
Maximum likelihood phylogenetic tree inferred for North American and eastern Asian *Castanea* samples using genome-wide data from 103,616 SNP loci. Asterisks indicate interspecific hybrids identified by STRUCTURE analysis. One asterisk (*) corresponds to an interspecific hybrid derived from *C. pumila* var. *ozarkensis*, *C. dentata*, and an unidentified East Asian *Castanea* sp.; two asterisks (**) correspond to a first-backcross descendant of *C. dentata* and *C. mollissima*. Bootstrap support for the nodes separating the highlighted taxa was 100% in all cases except the node separating *C. alabamensis* and *C. pumila* var. *pumila*, where bootstrap support was 91%. Scale bar is proportional to 0.03 substitutions/site.

The *C. dentata* clade was comprised of two distinct groups: (1) a group containing only individuals from the northern portion of the species’ range (samples from Maine, Pennsylvania, and Virginia) and individuals from the Blue Ridge of the southern portion of the species’ range (samples from North Carolina, South Carolina, and Georgia) and (2) a group containing only individuals from outside the Blue Ridge in the southern portion of the species’ range (samples from Piedmont of South Carolina, Interior Plateau of Tennessee, and Ridge and Valley and Southwestern Appalachians of Alabama). Bootstrap support for the two *C. dentata* groups was 100% (Appendix S6).

Within the *C. pumila* sensu lato clade, *C. alabamensis* and the different botanical varieties of chinquapin, *C. pumila* var. *pumila* and *C. pumila* var. *ozarkensis*, formed three distinct groups. The *C. pumila* var. *pumila* group was comprised of two subgroups: (1) a group containing only individuals from the southern Blue Ridge (samples from North Carolina, South Carolina, and Georgia) and (2) a group containing one individual from the southern Blue Ridge (a sample from South Carolina) and all individuals from the Coastal Plain (samples from South Carolina, Florida, and Georgia). Bootstrap support for the split separating *C. pumila* var. *pumila* samples into two groups was 100% (Appendix S6).

The phylogenetic placement of two samples did not match their putative species assignments. The first sample (dent hyb16_KY), thought to represent a *C. dentata* collection, was placed at an intermediate position between *C. dentata* and the East Asian *Castanea* species on the phylogenetic tree; review of our records showed that this sample was a first-backcross hybrid of *C. dentata* and *C. mollissima* ancestry. The second sample (ozar hyb63_MO), thought to represent *C. pumila* var. *ozarkensis*, was placed at an intermediate position between *C. pumila* sensu lato and *C. dentata* on the phylogenetic tree; STRUCTURE analysis results (detailed below) showed that this individual derived ancestry from *C. pumila* var. *ozarkensis*, *C. dentata*, and an undetermined East Asian *Castanea* species.

Mean genetic differentiation, as estimated by Weir and Cockerham’s (1984) *FST*, ranged from 0.71 (*C. mollissima* – *C. pumila* var. *ozarkensis*) to 0.11 (*C. pumila* var. *pumila* – *C. alabamensis*) (Appendix S7). In contrast to what would be expected if *C. alabamensis* were a hybrid taxon derived from *C. dentata* and *C. pumila* var. *pumila*, genetic differentiation was greater between *C. alabamensis* and *C. dentata* (*FST* = 0.40) than between *C. pumila* var. *pumila* and *C. dentata* (*FST* = 0.38).

### STRUCTURE analyses

STRUCTURE analysis indicated the presence of three genetically differentiated clusters (*K* = 3) in dataset ‘NAC+EAC583’ (Fig. 3A) (see Appendix S7 for Δ*K* values) These three clusters corresponded to (1) *C. dentata*, (2) *C. pumila* var. *pumila*, *C. pumila* var. *ozarkensis*, and *C. alabamensis* combined, and (3) the eastern Asian *Castanea* samples. STRUCTURE did not identify a *C. dentata* contribution to *C. alabamensis* genomes (Fig. 3A). Although one *C. alabamensis* sample, AG87_AL, appeared to have a small fraction of *C. dentata* ancestry (*C. dentata* ancestry proportion estimate = 0.05), this sample had the highest percentage of missing SNPs (82% missing SNPs) of any samples in the ‘NAC+EAC583’ dataset and the lowest number of retained reads of any plants included in analyses; this result is likely an artifact of low sample quality.

**Fig. 3.**
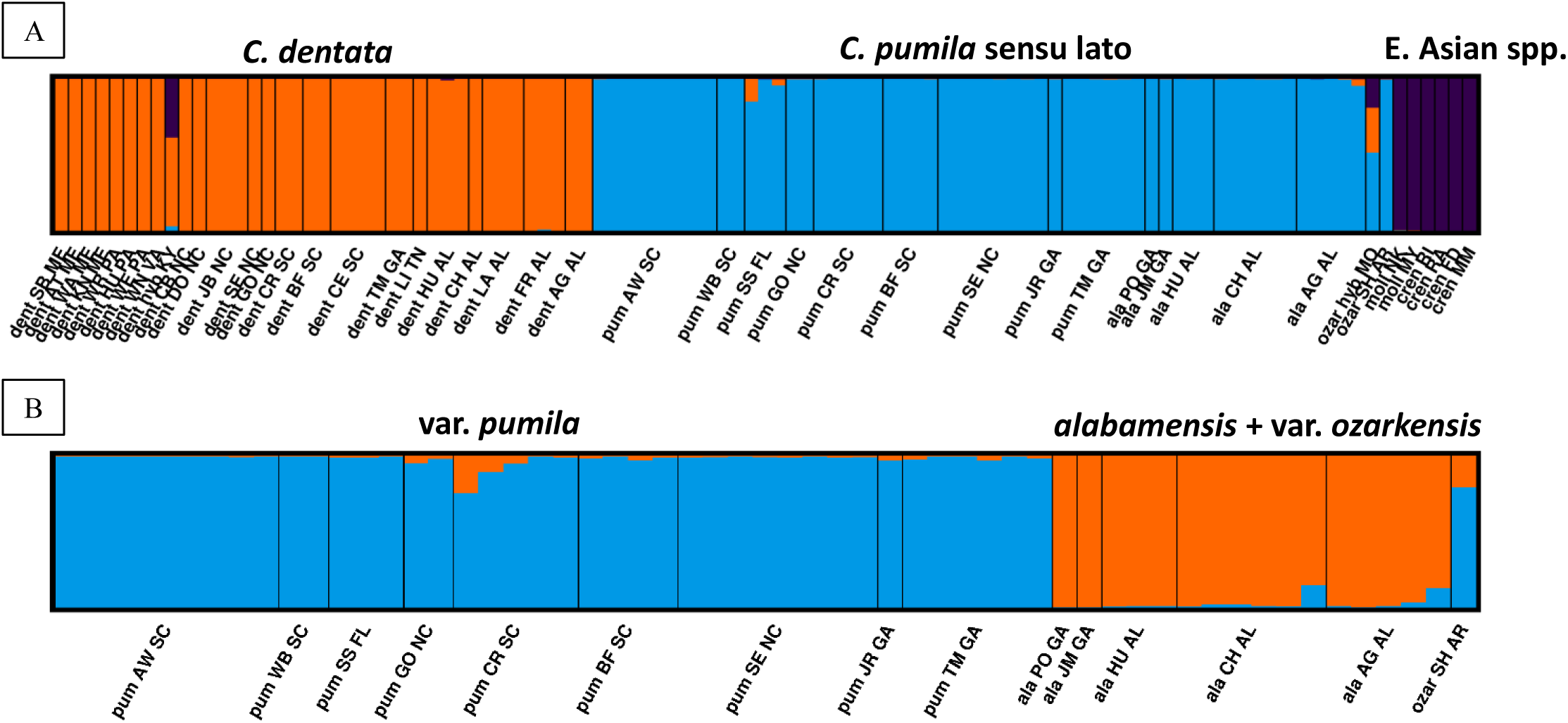
Bar plots of ancestry proportions from STRUCTURE analyses of (A) North American and eastern Asian *Castanea* samples and (B) North American chinquapin samples only. Each column represents the ancestry proportion estimate for an individual plant, with each individual’s estimated membership fraction illustrated by colored segments that correspond to *K* inferred clusters in the dataset. Black vertical lines group individuals according to their sample sites. Sample site information along the lower edge of the plots indicate species, site code, and state (e.g., pum AW SC). (A) Results of analysis of 583 SNP loci in 103 plants representing *C. alabamensis*, *C. pumila* var. *pumila*, *C. pumila* var. *ozarkensis*, *C. dentata*, *C. mollissima*, and *C. crenata*. The number of distinct genetic clusters identified (*K* = 3) corresponds to *C. dentata*, *C. pumila* sensu lato, and the sampled eastern Asian *Castanea* spp., *C. mollissima* and *C. crenata*. (B) Results of analysis of 583 SNP loci in 57 individuals representing *C. alabamensis*, *C. pumila* var. *pumila*, and *C. pumila* var. *ozarkensis*. The number of distinct genetic clusters identified (*K* = 2) corresponds to *C. pumila* var. *pumila* and *C. alabamensis*/*C. pumila* var. *ozarkensis*.

Analysis of *C. dentata* samples also failed to detect a genetic contribution from *C. pumila* sensu lato in sympatric populations. However, two *C. pumila* var. *pumila* individuals, both from the allopatric site SS_FL, had evidence of *C. dentata* ancestry. Of these two admixed samples, SS80_FL, was estimated to derive as much as 0.15 of its genome from *C. dentata*.

STRUCTURE analysis of the North American chinquapins *Castanea pumila* sensu lato (including *C. alabamensis*) indicated the presence of two genetically differentiated clusters (*K* = 2)—the first corresponding to *C. pumila* var. *pumila* and the second corresponding to *C. alabamensis* and *C. pumila* var. *ozarkensis* combined (Fig. 3B; see Appendix S7 for Δ*K* values). Low to moderate levels of admixture between the different botanical varieties of chinquapin were found in the majority of populations analyzed. The single *C. pumila* var. *ozarkensis* individual (ozar_SH_AR) remaining after removal of the interspecific hybrid from Missouri showed evidence of substantial admixture with *C. pumila* var. *pumila*. Evidence of admixture was also present in *C. alabamensis* populations, with higher levels of *C. pumila* var. *pumila* ancestry in the more southerly *alabamensis* populations, CH_AL and AG_AL (Fig. 3B).

### Principal component analysis

To provide an additional test of the hybridization hypothesis for *C. alabamensis* and to better understand partitioning of genetic variation within the North American clade, we performed PCA using the 80 North American *Castanea* samples in dataset ‘NAC’ with <30% missing SNP loci. Principal component analysis of the North American *Castanea* identified 32 significant principal components (PCs) that explained 56.52% of the total variation (Appendix S7). The first two principal components provided clear separation of the three currently recognized North American taxa and *C. alabamensis* from one another (Fig. 4). The first component explained 10.96% of the total variation and separated *C. dentata* samples from samples of *C. alabamensis*, *C. pumila* var. *pumila*, and *C. pumila* var. *ozarkensis* (Fig. 4; Appendix S7). The second component explained 2.75% of the total variation and separated *C. pumila* var. *pumila*, *C. pumila* var. *ozarkensis*, and *C. alabamensis* from one another. *C. pumila* var. *pumila* samples from the Coastal Plain of Florida and South Carolina were placed along the leftmost portion of the axis created by PC2 in Fig. 4 (i.e., lower values for PC2). However, samples from other parts of the geographical distribution of *C. pumila* var. *pumila* were not completely excluded from the leftmost portion of PC2, as one sample from the Blue Ridge province of South Carolina, CR6_SC, clustered with Coastal Plain samples. *C. pumila* var. *pumila* samples from sites farther inland—specifically, the Blue Ridge of South Carolina, North Carolina, and Georgia—clustered near the midrange of PC2. In contrast to what would be expected for a hybrid taxon, PCA did not place any *C. alabamensis* samples intermediate to the *C. pumila* sensu lato and *C. dentata* clusters.

**Fig. 4.**
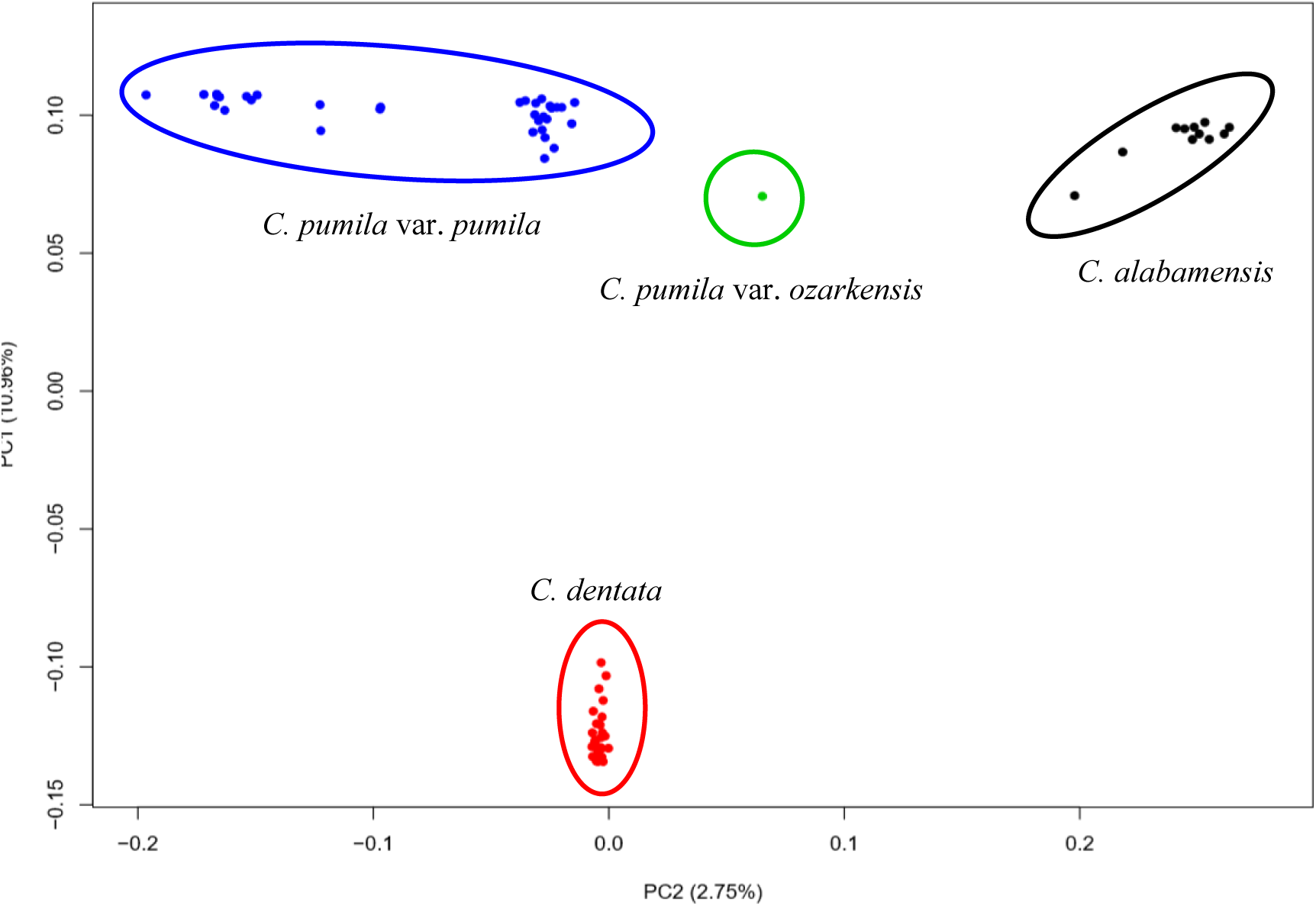
Graph of the first two axes from a principal components analysis of 80 individuals representing the putative hybrid, *C. alabamensis*, and all other North American *Castanea* taxa. Dot colors correspond to the following taxa: blue = *C. pumila* var. *pumila*, green = *C. pumila* var. *ozarkensis*, black = *C. alabamensis*, and red = *C. dentata*.

## DISCUSSION

### C. alabamensis is a distinct variety of North American Castanea that includes plants previously identified as hybrids between Castanea species

While agreeing with previous studies that North American *Castanea* is composed of two species, *C. dentata* and *C. pumila* sensu lato, we have presented compelling evidence that plants identified by Ashe (1925) as *C. alabamensis* comprise a distinct variety within *C. pumila* sensu lato along with *C. pumila* var. *pumila* and *C. pumila* var. *ozarkensis*. Our study represents the first use of genome-wide data to understand evolution and hybridization in the North American *Castanea* species. With different population sampling methods, our analytical approach can be applied to other questions of importance for conservation and restoration of the American chestnut and chinquapins. Our findings provide much needed clarification of the evolutionary relationships, species boundaries, and history of admixture of the North American *Castanea* taxa and will allow workers in the field of chestnut and chinquapin conservation to more efficiently conserve and restore the biodiversity of these imperiled taxa.

In contrast to multiple recent studies of closely-related sympatric species in the Fagaceae (Cavender-Bares and Pahlich, 2009; Leroy et al. 2017; Kim et al. 2018), we find that *C. dentata* and *C. pumila* are both genetically and morphologically discrete where they co-occur, with no evidence of local hybridization and introgression. Jaynes (1964) reported that fertile hybrids can be made among all *Castanea* species. The reproductive barrier between the species is most likely flowering time; *C. pumila* blooms 1-2 weeks earlier than *C. dentata* (P.H. Sisco, J.H. Craddock, unpublished data).

### Admixture among taxa

Although we did not document admixture with *C. dentata* in our *C. alabamensis* samples, many other cases of intra- and interspecific admixture were identified. Perhaps the most interesting case of interspecific admixture that we documented involved samples from a morphologically intriguing population of *C. pumila* var. *pumila* in Florida. Despite the stark morphological differences between these *C. pumila* var. *pumila* individuals and *C. dentata*, two of three individuals had evidence of introgression from *C. dentata* and one of the plants had an ancestry proportion estimate of 0.15 from *C. dentata* in STRUCTURE. These plants, which were perhaps the most morphologically unique of the study, are the rhizomatous subshrub form (only ∼0.5-2 m tall at reproductive maturity) of chinquapin that have been treated as *C. alnifolia* (common name: trailing chinquapins) in older taxonomic works (Nuttall, 1818; Sargent, 1919; Ashe, 1922). These Florida *C. pumila* populations are currently separated from the southernmost known *C. dentata* populations by hundreds of kilometers. The finding of *C. dentata* ancestry in these *C. pumila* plants is consistent with the hypothesis that the range of *C. dentata* once extended much farther south than is currently observed (Davis, 1983). While other population genetics studies have inferred a post-Pleistocene expansion of *C. dentata* populations from south to north along the Appalachians (e.g., Gailing and Nelson, 2017), our finding of *C. dentata* ancestry in *C. pumila* populations from Florida is the most direct genetic evidence produced in support of a Pleistocene refugium for *C. dentata* in the southeastern Coastal Plain.

Finally, many cases of intraspecific admixture within the chinquapin clade were also documented in our data. Within *C. pumila* sensu lato, admixture between the two genetic groups identified by STRUCTURE—(1) *C. pumila* var. *pumila* and (2) *C. alabamensis* and *C. pumila* var. *ozarkensis*—was observed in most populations, indicating a recent history of shared ancestry between the different botanical varieties of chinquapin. It should be noted, however, that none of the chinquapin populations analyzed in this study were collected from areas of sympatry for distinct chinquapin varieties (i.e., where *C. pumila* var. *pumila* and *C. pumila* var. *ozarkensis* occur at the same site or within effective pollination distance). Thus, levels of admixture in sympatric populations containing multiple chinquapin varieties cannot be assessed from our data. Given the low to moderate levels of admixture between chinquapin varieties documented in allopatric populations in the present study, we expect sympatric populations to display even greater signatures of shared ancestry.

## CONCLUSIONS

We have used genome-wide SNP data and morphology to show that *C. alabamensis* is a distinct variety of North American chinquapin (*C. pumila* sensu lato) that includes plants previously identified morphologically as hybrids between *Castanea* species. A combination of genome-wide genotyping and morphological analysis was required to better understand the origin of the putative hybrids and the nature of species boundaries in North American *Castanea*. Presumed naturally-occurring admixtures between different *Castanea* species and varieties were found, most notably between *C. pumila* var. *pumila* in northern Florida and *C. dentata*. Our results demonstrate the capability of genomic approaches to resolve previously intractable questions of *Castanea* evolution and highlight the need for further exploration of *Castanea* diversity.

## ACKNOWLEDGMENTS

We acknowledge funding support from the Foundation for the Carolinas (by philanthropists Shelli and Brad Stanback), the University of Tennessee at Chattanooga Department of Biology, Geology, & Environmental Science, the Southern Appalachian Botanical Society, the University of Tennessee at Chattanooga Graduate School, and the American Chestnut Foundation. We thank Lisa W. Alexander for helpful discussions regarding chestnut hybridization. We acknowledge field sampling guidance and private property access provided by David Morris, Ed Schwartzman, Robert Simons, Jack Agricola, Bruce and Francine Hutchinson, Gary Carver, Larry Brasher, Glenda Frames, Marty Schulman, Will Calhoun, Jim Lacefield, Donald Hagan, and Timothy Spira. We thank Mary Klinghard, Cameron Perkins, Jessica Tzeng, Brooke Hadden, and Conrad Blunck for assistance with field sampling. We thank Dr. Ethan Hereth for help with the NCBI SRA data upload. Trent Deason generously shared early access to his voucher specimens from the Ruffner Mtn. sample site. Caleb Powell provided valuable field sampling assistance and herbarium database curation. Dan Thornton and Stylianos Chatzimanolis provided helpful guidance with leaf macrophotography. Information regarding naturally-occurring American chestnut trees in the breeding program of the American Chestnut Foundation was provided by Kendra Collins and Eric Evans. Xiaoxia Xia, Rooksana Noorai, Chris Saski, and the Clemson University Genomics and Computational Biology Laboratory generously provided bioinformatic and analytical assistance. We thank the University of North Carolina Chapel Hill Herbarium for the generous loan of museum specimens used in this study.

## DATA ACCESSIBILITY

Sequence data are available at the NCBI SRA (BioProject ID: PRJNA541592). All datasets used in analyses are available on GitHub: https://github.com/MTPerkins/Nonhybrid_origin_of_Castanea_alabamensis.

## APPENDICES

**Appendix S1.** Sample voucher and NCBI SRA information for individual *Castanea* plants genotyped. GBS IDs correspond to sample names in NCBI SRA BioProject ID “PRJNA541592”.

| County, State or<br>cultivar/selection<br>name | Sample<br>site<br>code | GBS<br>ID | Taxon | Herbarium<br>voucher | DNA tube<br>label |
| --- | --- | --- | --- | --- | --- |
| Charleston, SC | AW | 1 | *C. pumila* var. *pumila* | Perkins 15-1 | AW1 |
| Charleston, SC | AW | 5 | *C. pumila* var. *pumila* | Perkins 15-2 | AW2 |
| Charleston, SC | AW | 9 | *C. pumila* var. *pumila* | Perkins 15-3 | AW3 |
| Charleston, SC | AW | 13 | *C. pumila* var. *pumila* | Perkins 15-4 | AW4 |
| Charleston, SC | AW | 17 | *C. pumila* var. *pumila* | Perkins 15-5 | AW5 |
| Charleston, SC | AW | 23 | *C. pumila* var. *pumila* | Perkins 15-7 | AW7 |
| Charleston, SC | AW | 28 | *C. pumila* var. *pumila* | Perkins 15-8 | AW8 |
| Charleston, SC | AW | 32 | *C. pumila* var. *pumila* | Perkins 15-9 | AW9 |
| Charleston, SC | AW | 36 | *C. pumila* var. *pumila* | Perkins 15-10 | AW10 |
| Sumter, SC | WB | 41 | *C. pumila* var. *pumila* | Perkins 15-11 | WB1 |
| Sumter, SC | WB | 44 | *C. pumila* var. *pumila* | Perkins 15-12 | WB2 |
| Transylvania, NC | GO | 18 | *C. dentata* | Perkins 15-13 | G1 |
| Transylvania, NC | GO | 20 | *C. pumila* var. *pumila* | Perkins 15-14 | G2 |
| Transylvania, NC | GO | 24 | *C. pumila* var. *pumila* | Perkins 15-15 | G3 |
| Oconee, SC | CR | 7 | *C. dentata* | Perkins 15-16 | CR1 |
| Oconee, SC | CR | 11 | *C. dentata* | Perkins 15-17 | CR2 |
| Oconee, SC | CR | 15 | *C. pumila* var. *pumila* | Perkins 15-18 | CR3 |
| Oconee, SC | CR | 19 | *C. pumila* var. *pumila* | Perkins 15-19 | CR4 |
| Oconee, SC | CR | 21 | *C. pumila* var. *pumila* | Perkins 15-20 | CR5 |
| Oconee, SC | CR | 25 | *C. pumila* var. *pumila* | Perkins 15-21 | CR6 |
| Oconee, SC | CR | 30 | *C. pumila* var. *pumila* | Perkins 15-22 | CR7 |
| Oconee, SC | BF | 29 | *C. pumila* var. *pumila* | Perkins 15-23 | BF1 |
| Oconee, SC | BF | 33 | *C. pumila* var. *pumila* | Perkins 15-24 | BF2 |
| Oconee, SC | BF | 37 | *C. pumila* var. *pumila* | Perkins 15-25 | BF3 |
| Oconee, SC | BF | 42 | *C. pumila* var. *pumila* | Perkins 15-26 | BF4 |
| Oconee, SC | BF | 45 | *C. dentata* | Perkins 15-27 | BF5 |
| Oconee, SC | BF | 3 | *C. dentata* | Perkins 15-28 | BF6 |
| Pickens, SC | CE | 2 | *C. dentata* | Perkins 15-29 | CEF1 |
| Pickens, SC | CE | 6 | *C. dentata* | Perkins 15-30 | CEF2 |
| Pickens, SC | CE | 10 | *C. dentata* | Perkins 15-31 | CEF3 |
| Pickens, SC | CE | 14 | *C. dentata* | Perkins 15-32 | CEF4 |
| Suwannee, FL | SS | 80 | *C. pumila* var. *pumila* | Perkins 17-1 | SS2 |
| Suwannee, FL | SS | 81 | *C. pumila* var. *pumila* | Perkins 17-2 | SS3 |
| Suwannee, FL | SS | 82 | *C. pumila* var. *pumila* | Perkins 17-3 | SS4 |
| Lincoln, TN | LI | 83 | *C. dentata* | Perkins 17-4 | TNLIN1 |
| Calhoun, AL | CH | 90 | *C. alabamensis* | Perkins 17-5 | CH21 |
| Calhoun, AL | CH | 91 | *C. dentata* | Perkins 17-6 | CH22 |
| Calhoun, AL | CH | 92 | *C. alabamensis* | Perkins 17-7 | CH23 |
| Calhoun, AL | CH | 93 | *C. alabamensis* | Perkins 17-8 | CH24 |
| Calhoun, AL | CH | 94 | *C. alabamensis* | Perkins 17-9 | CH25 |
| Calhoun, AL | CH | 95 | *C. alabamensis* | Perkins 17-10 | CH26 |
| Calhoun, AL | CH | 96 | *C. alabamensis* | Perkins 17-11 | CH27 |
| Cleburne, AL | HU | 67 | *C. alabamensis* | Perkins 17-12 | H1 |
| Cleburne, AL | HU | 68 | *C. dentata* | Perkins 17-13 | H2 |
| Cleburne, AL | HU | 69 | *C. dentata* | Perkins 17-14 | H3 |
| Cleburne, AL | HU | 70 | *C. dentata* | Perkins 17-15 | H4 |
| Cleburne, AL | HU | 86 | *C. alabamensis* | Perkins 17-16 | H9 |
| Cleburne, AL | HU | 89 | *C. alabamensis* | Perkins 17-17 | H6 |
| Cleburne, AL | FR | 71 | *C. dentata* | Perkins 17-18 | F1 |
| Cleburne, AL | FR | 72 | *C. dentata* | Perkins 17-19 | F5 |
| Cleburne, AL | FR | 84 | *C. dentata* | Perkins 17-20 | F3 |
| Clay, AL | AG | 73 | *C. alabamensis* | Perkins 17-21 | AG1 |
| Clay, AL | AG | 74 | *C. dentata* | Perkins 17-22 | AG2 |
| Clay, AL | AG | 75 | *C. alabamensis* | Perkins 17-23 | AG3 |
| Clay, AL | AG | 76 | *C. alabamensis* | Perkins 17-24 | AG6 |
| Clay, AL | AG | 77 | *C. dentata* | Perkins 17-25 | AG7 |
| Clay, AL | AG | 85 | *C. alabamensis* | Perkins 17-26 | AG4 |
| Clay, AL | AG | 87 | *C. alabamensis* | Perkins 17-27 | AG5 |
| Morgan, AL | LA | 78 | *C. dentata* | Perkins 17-28 | L14 |
| Morgan, AL | LA | 79 | *C. dentata* | Perkins 17-29 | L19 |
| Morgan, AL | LA | 88 | *C. dentata* | Perkins 17-30 | L10 |
| Floyd, GA | JM | 22 | *C. alabamensis* | Binkley 412 | 412 |
| Floyd, GA | PO | 38 | *C. alabamensis* | Binkley 420 | 420 |
| Cherokee, NC | JB | 50 | *C. dentata* | Binkley 507 | 507 |
| Cherokee, NC | JB | 51 | *C. dentata* | Binkley 513 | 513 |
| Cherokee, NC | JB | 52 | *C. dentata* | Binkley 514 | 514 |
| Cherokee, NC | SE | 53 | *C. pumila* var. *pumila* | Binkley 124 | 124 |
| Cherokee, NC | SE | 54 | *C. pumila* var. *pumila* | Binkley 525 | 525 |
| Cherokee, NC | SE | 55 | *C. pumila* var. *pumila* | Binkley 126 | 126 |
| Cherokee, NC | SE | 56 | *C. pumila* var. *pumila* | Binkley 527 | 527 |
| Cherokee, NC | SE | 57 | *C. pumila* var. *pumila* | Binkley 528 | 528 |
| Cherokee, NC | SE | 58 | *C. pumila* var. *pumila* | Binkley 529 | 529 |
| Cherokee, NC | SE | 59 | *C. dentata* | Binkley 531 | 531 |
| Cherokee, NC | SE | 60 | *C. pumila* var. *pumila* | Binkley 532 | 532 |
| Cherokee, NC | SE | 61 | *C. pumila* var. *pumila* | Binkley 535 | 535 |
| Cherokee, NC | SE | 62 | *C. pumila* var. *pumila* | Binkley 149 | 149 |
| Baker, GA | JR | 26 | *C. pumila* var. *pumila* | Binkley 735 | 735 |
| Pickens, GA | TM | 27 | *C. dentata* | Binkley 645 | 645 |
| Pickens, GA | TM | 31 | *C. dentata* | Binkley 646 | 646 |
| Pickens, GA | TM | 35 | *C. pumila* var. *pumila* | Binkley 647 | 647 |
| Pickens, GA | TM | 40 | *C. pumila* var. *pumila* | Binkley 648 | 648 |
| Pickens, GA | TM | 43 | *C. pumila* var. *pumila* | Binkley 649 | 649 |
| Pickens, GA | TM | 46 | *C. pumila* var. *pumila* | Binkley 650 | 650 |
| Pickens, GA | TM | 47 | *C. pumila* var. *pumila* | Binkley 651 | 651 |
| Pickens, GA | TM | 48 | *C. pumila* var. *pumila* | Binkley 652 | 652 |
| Pickens, GA | TM | 49 | *C. pumila* var. *pumila* | Binkley 653 | 653 |
| Knox, ME | KN | 34 | *C. dentata* | Binkley 671 | 671 |
| Piscataquis, ME | AT | 39 | *C. dentata* | Binkley 672 | 672 |
| Waldo, ME | WA | 65 | *C. dentata* | Binkley 673 | 673 |
| Piscataquis, ME | SB | 66 | *C. dentata* | Binkley 674 | 674 |
| Huntingdon, PA | RO | 4 | *C. dentata* | Binkley 669 | 669 |
| Warren, PA | WR | 8 | *C. dentata* | Binkley 710 | 710 |
| Westmoreland, PA | WE | 12 | *C. dentata* | Binkley 675 | 675 |
| Sharp, AR | SH | 64 | *C. pumila* var. *ozarkensis* | Binkley 705 | 705 |
| Lawrence, MO | MO | 63 | *ozarkensis* complex hybrid | Binkley 694 | 694 |
| Adair, KY | KY | 16 | *dentata-mollissima* hybrid | NA | KYADA1 |
| Buncombe, NC | DO | 102 | *C. dentata* | Sisco & Perkins 15-1 | NCDOT |
| Avery, NC | CB | 103 | *C. dentata* | NA | Cranberry |
| Washington, VA | WN | 97 | *C. dentata* | NA | AD98 |
| cultivar 'Nanking' | NK | 100 | *C. mollissima* | NA | Nanking |
| cultivar 'Mahogany' | MY | 98 | *C. mollissima* | NA | Mahogany |
| selection 'Fort Defiance' | FD | 104 | *C. crenata* | NA | Ft. Defiance |
| selection 'Morrow Mountain' | MM | 107 | *C. crenata* | NA | Morrow Mtn. |
| selection 'Rita 1' | RI | 108 | *C. crenata* | NA | Rita 1 |
| selection 'Rita 2' | RA | 110 | *C. crenata* | NA | Rita 2 |

**Appendix S2.** Annotations of plants studied by Binkley (2008), Shaw et al. (2012), and Li and Dane (2013). Two plants from the study of Binkley (2008) and Shaw et al. (2012) were genotyped here. Herbarium vouchers from all plants listed below were assessed for morphological traits listed in Table 1 and annotated.

| County, State | Sample site | Sample ID in previous study | Taxonomic determination of previous study | Reference | Taxonomic determination of present study | Herbarium voucher | Sample ID in present study |
| --- | --- | --- | --- | --- | --- | --- | --- |
| Floyd, GA | the Pocket | 420 | *C. x neglecta* | Binkley (2008); Shaw et al. (2012) | *C. alabamensis* | Binkley 420 | PO38_AL |
| Floyd, GA | Johns Mountain | 412 | *C. x neglecta* | Binkley (2008); Shaw et al. (2012) | *C. alabamensis* | Binkley 412 | JM22_AL |
| Jefferson, AL | Ruffner Mountain | AL-M65 | Type II *C. dentata* | Li and Dane (2013) | *C. alabamensis* | Perkins 15-101 | Not genotyped |
| Jefferson, AL | Ruffner Mountain | AL-M35 | Type II *C. dentata* | Li and Dane (2013) | *C. alabamensis* | Perkins 15-102 | Not genotyped |
| Jefferson, AL | Ruffner Mountain | AL-M38 | Type I *C. dentata* | Li and Dane (2013) | *C. dentata* | Deason MS38 | Not genotyped |
| Jefferson, AL | Ruffner Mountain | AL-M68 | Type II *C. dentata* | Li and Dane (2013) | *C. alabamensis* | Deason MS68 | Not genotyped |
| Jefferson, AL | Ruffner Mountain | AL-4CN | Type I *C. dentata* | Li and Dane (2013) | *C. dentata* | Deason 4CN | Not genotyped |
| Jefferson, AL | Ruffner Mountain | AL-7CN | Type I *C. dentata* | Li and Dane (2013) | *C. dentata* | Deason 7CN | Not genotyped |

**Appendix S3.**
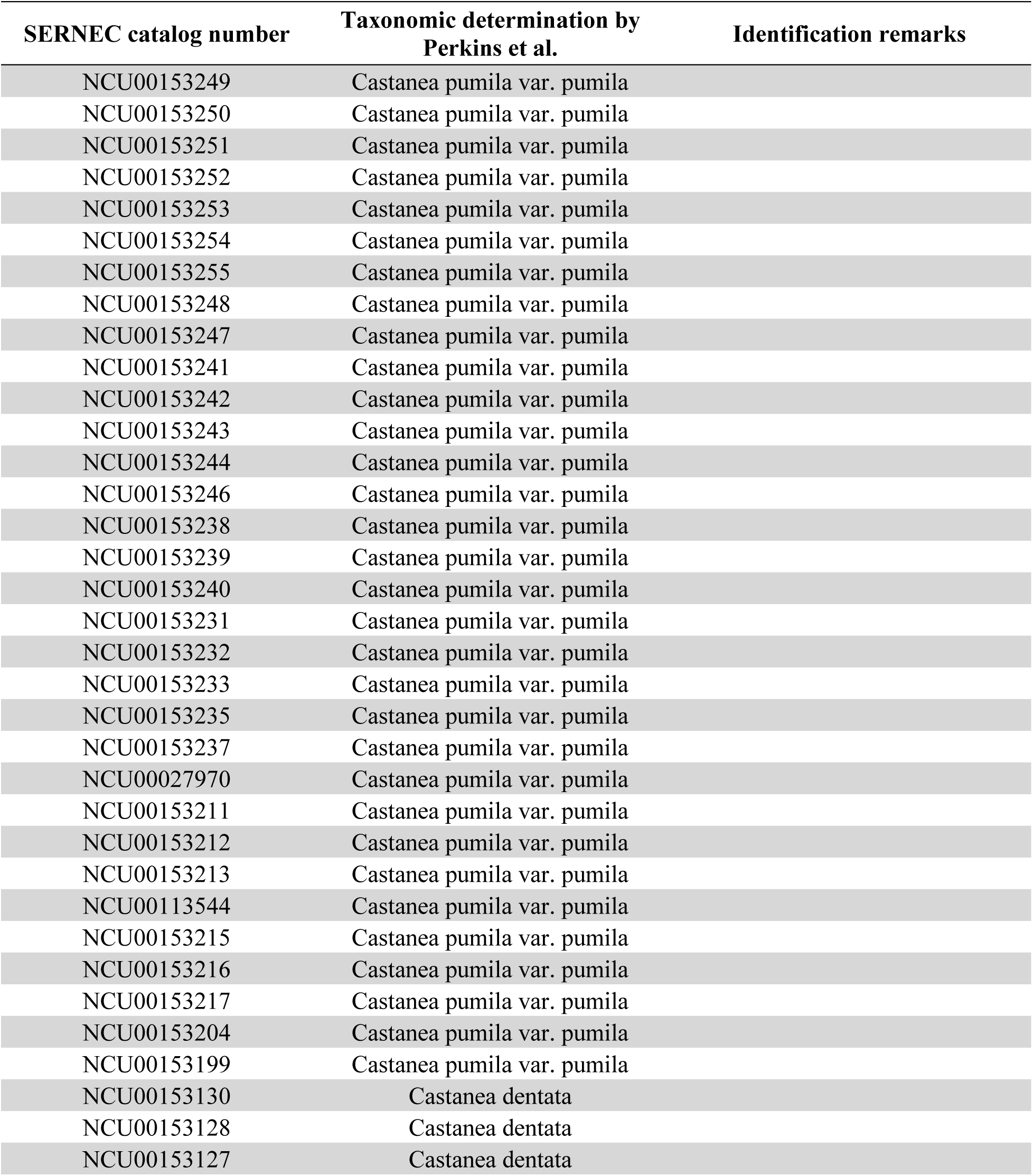
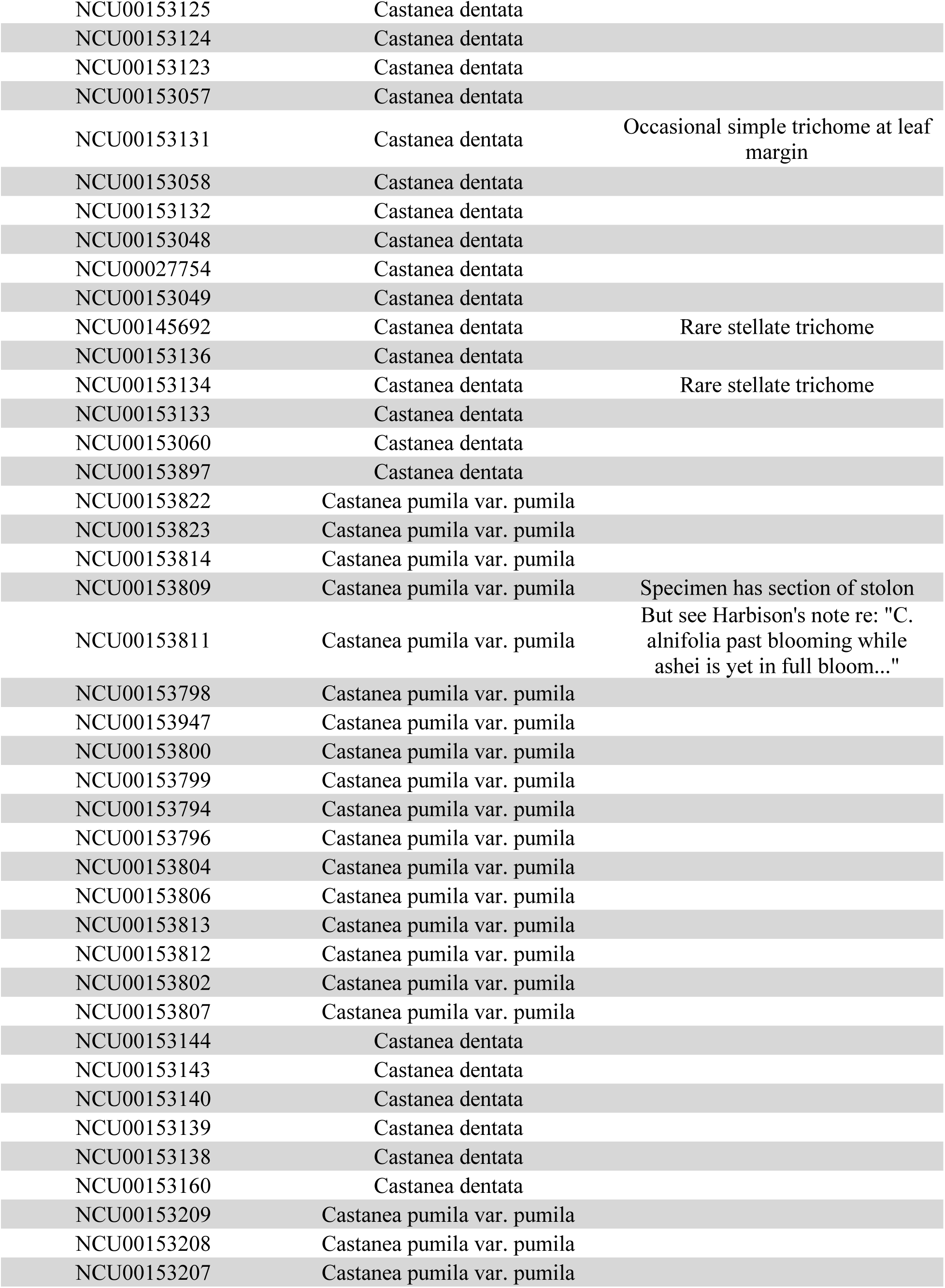
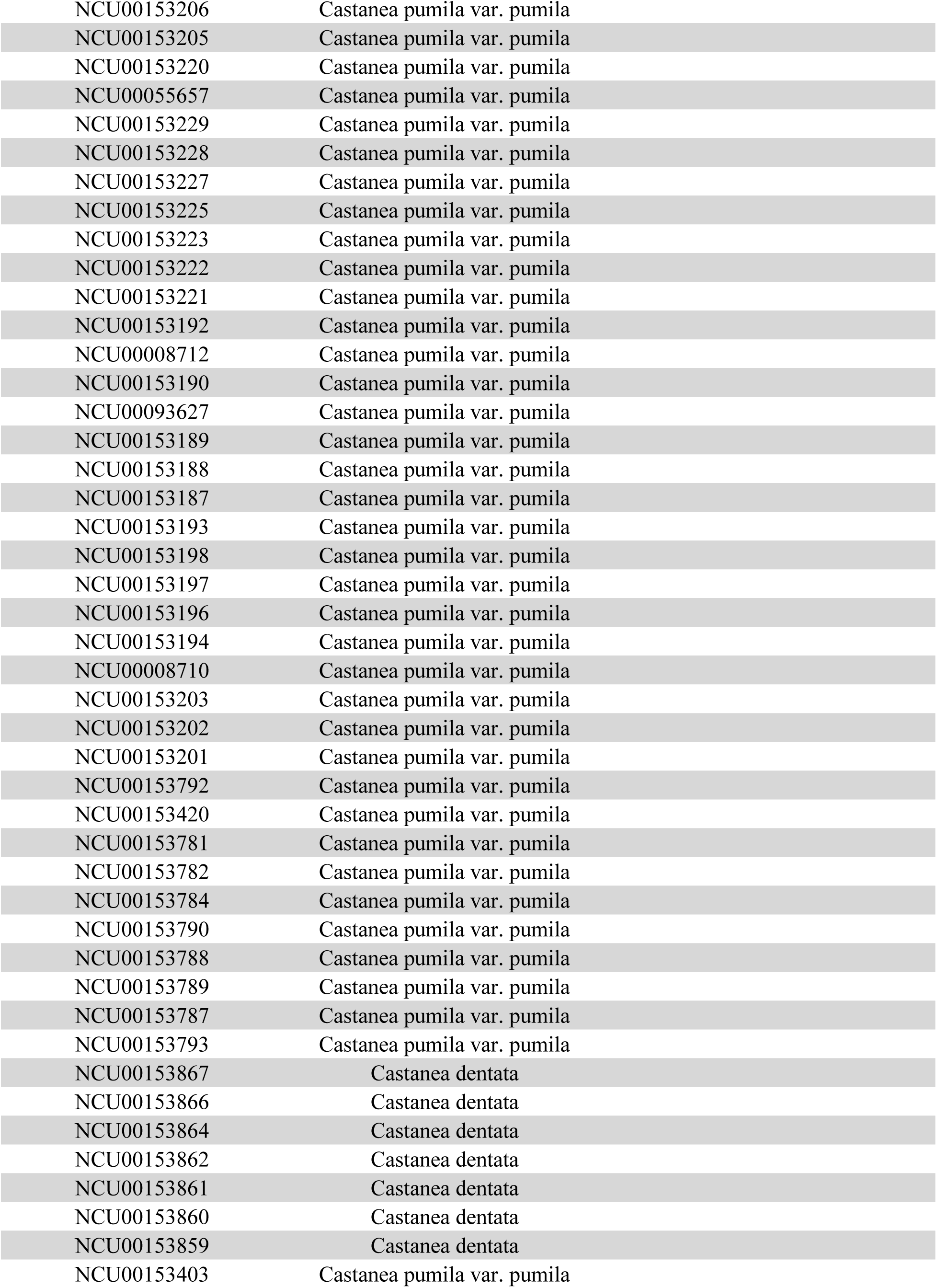
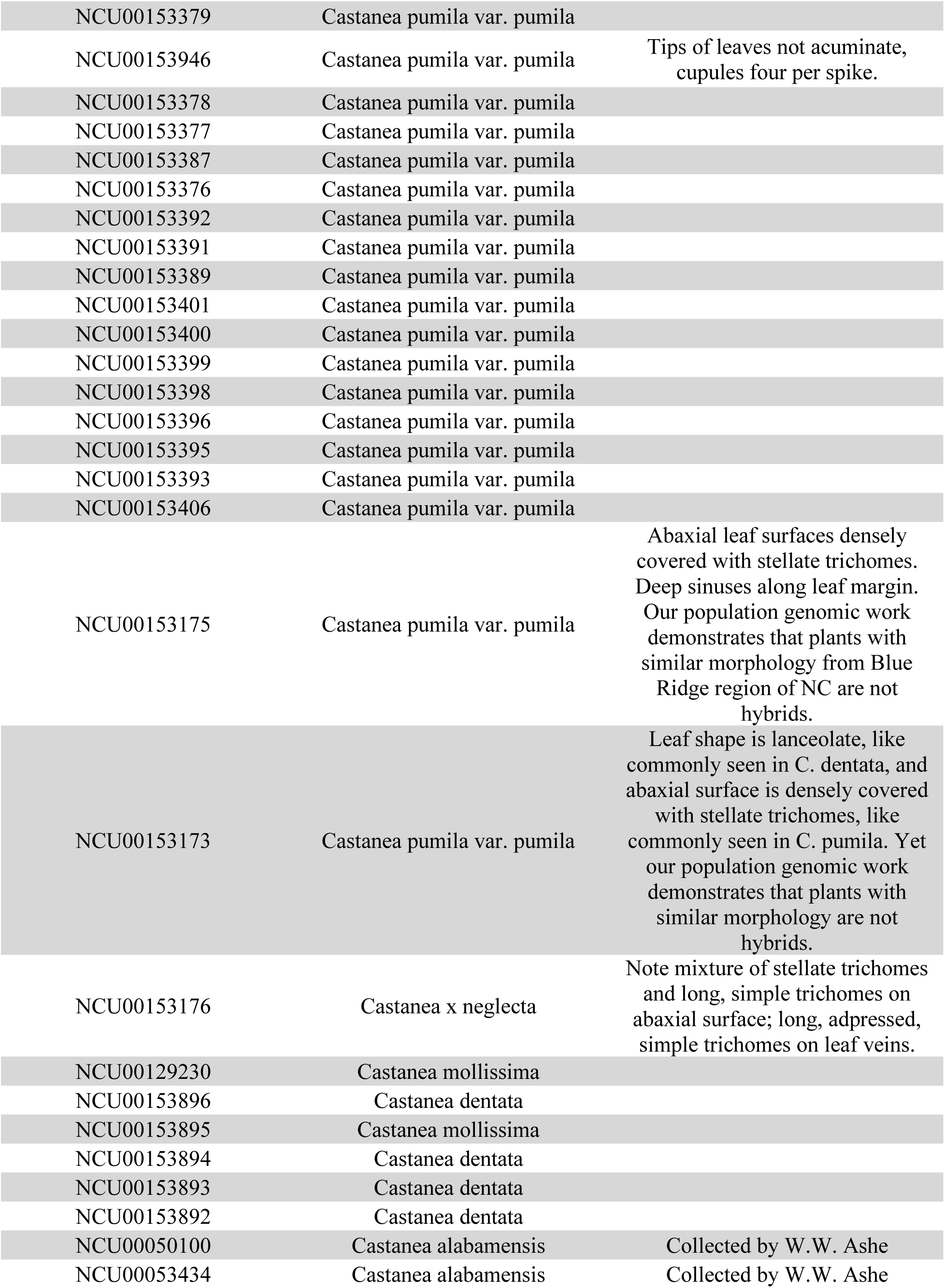
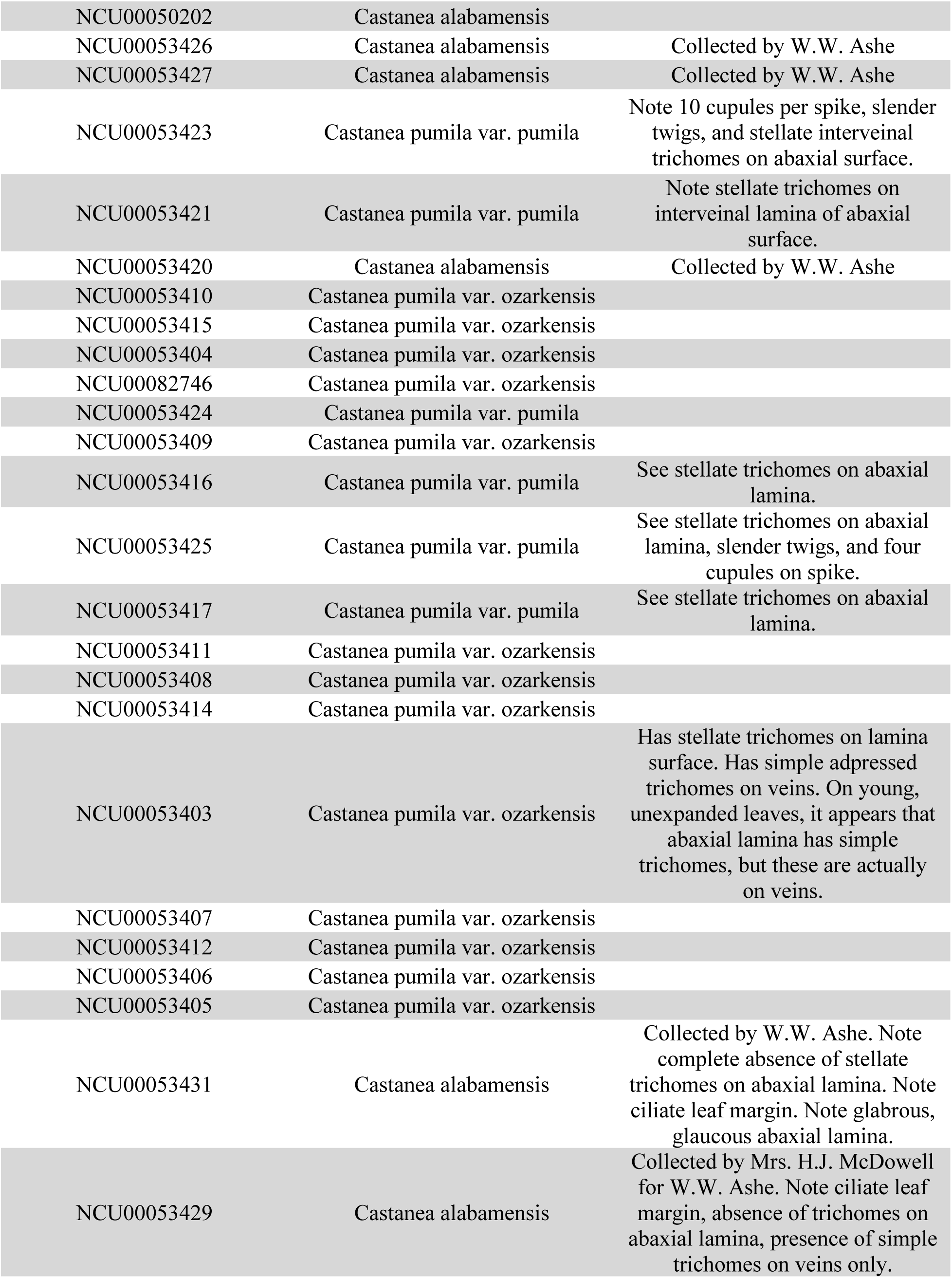
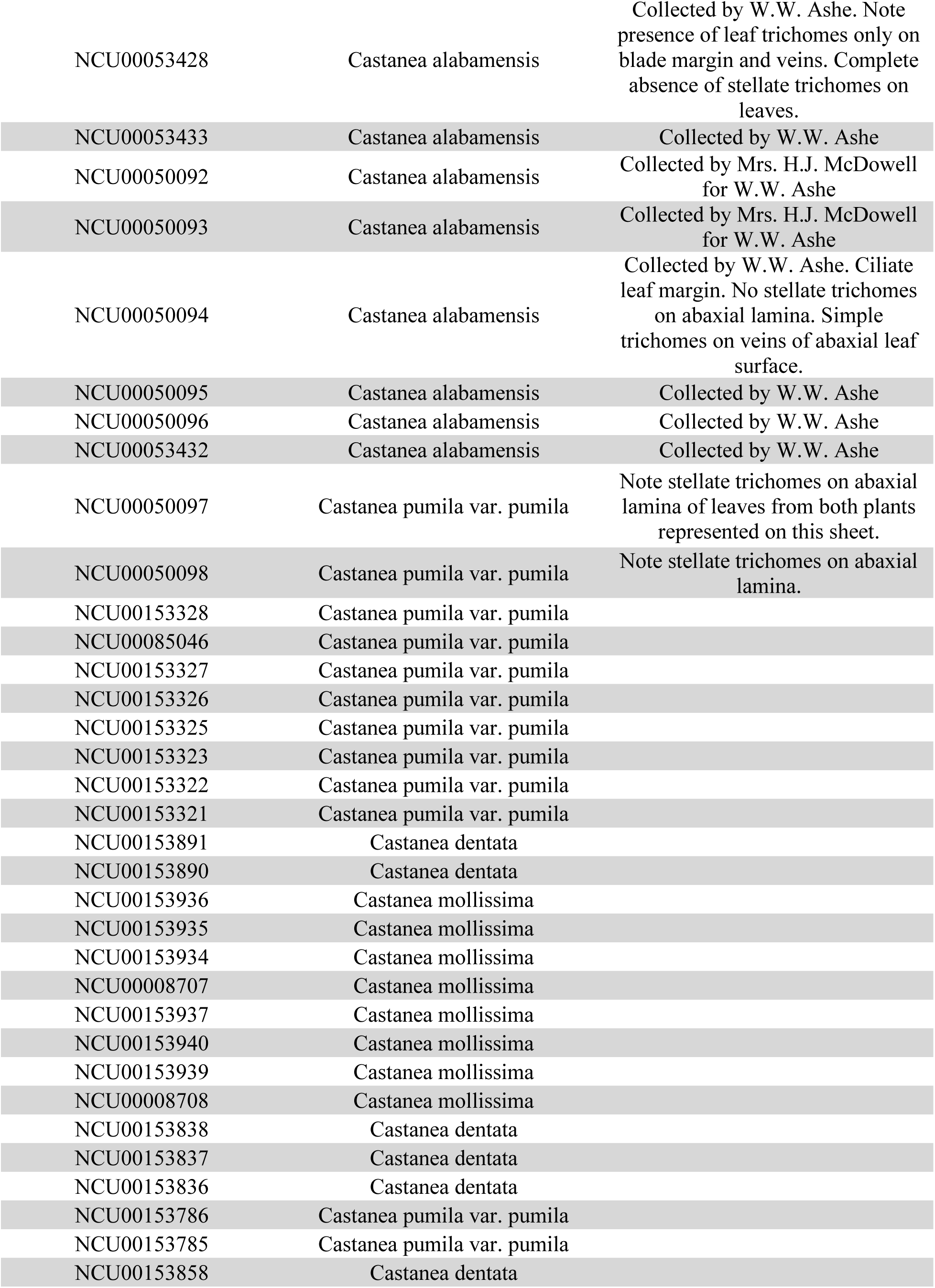
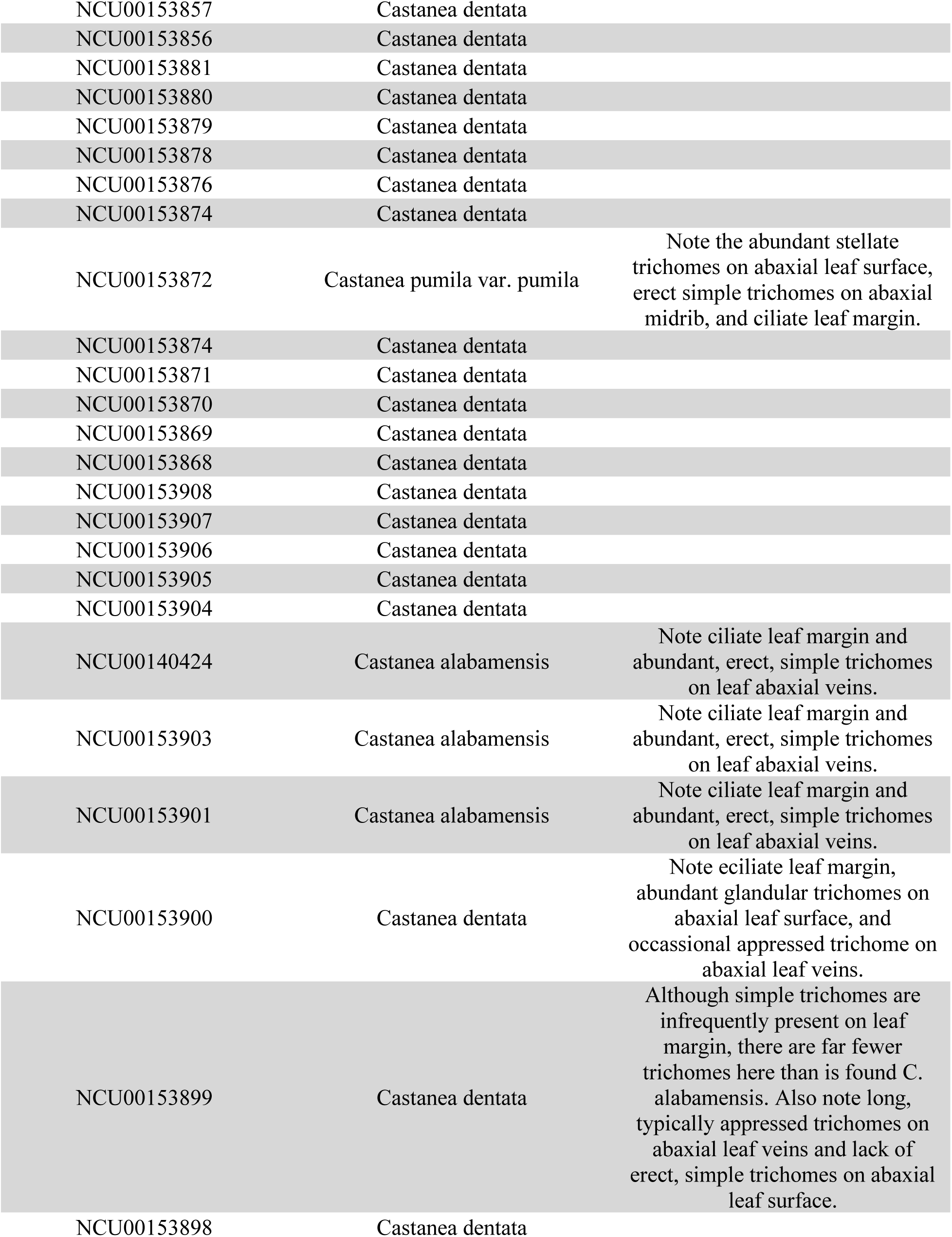
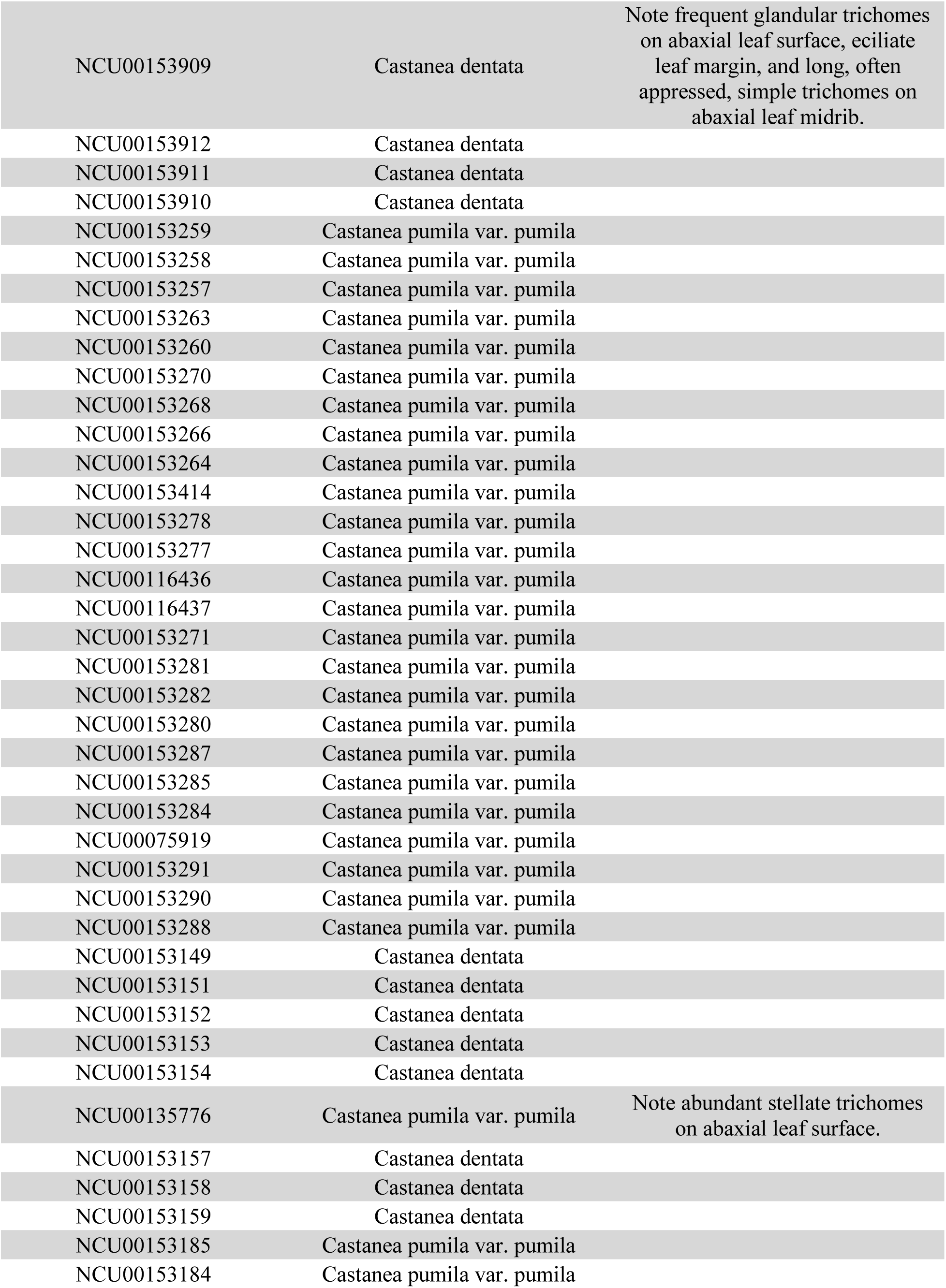
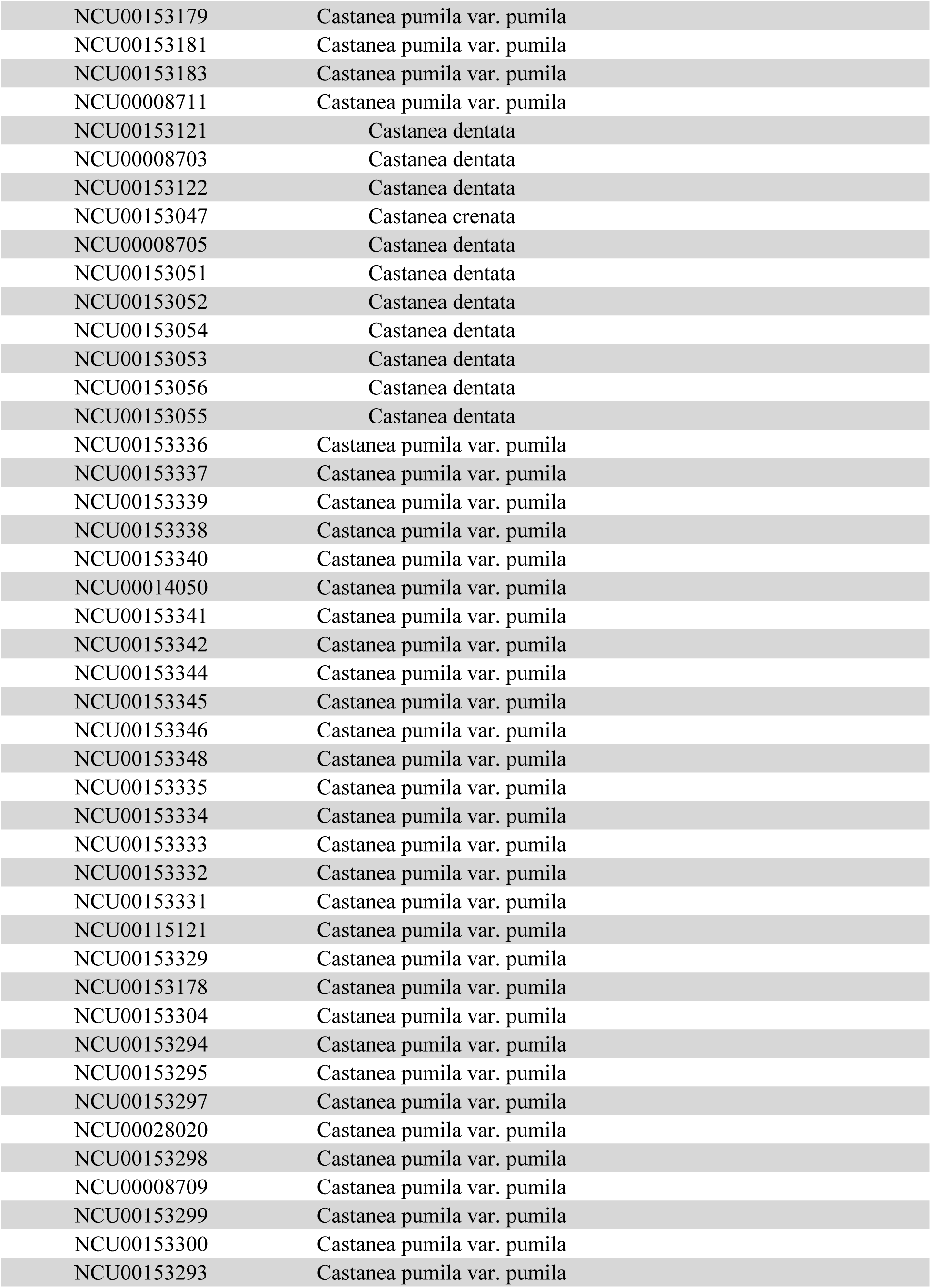
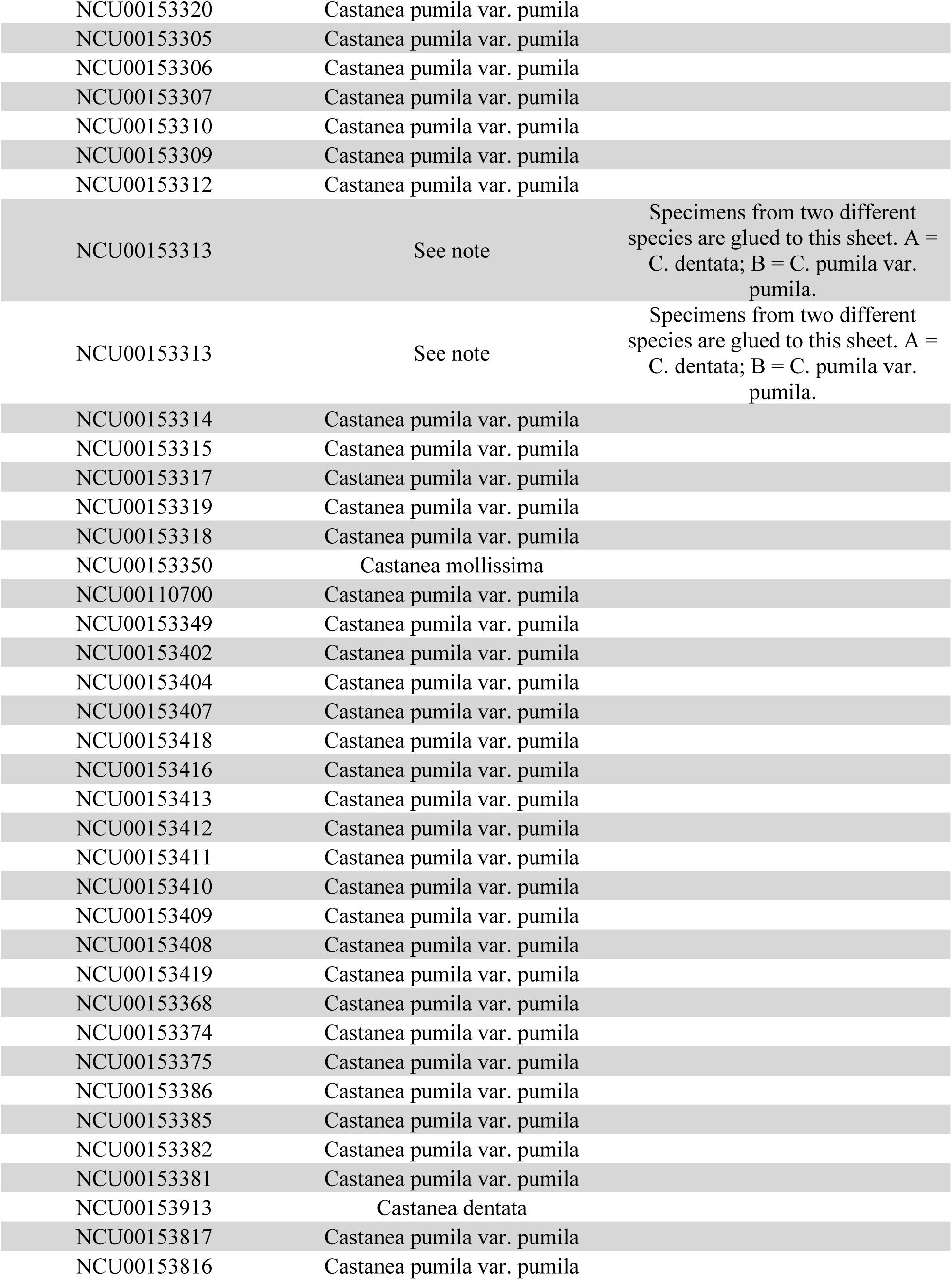
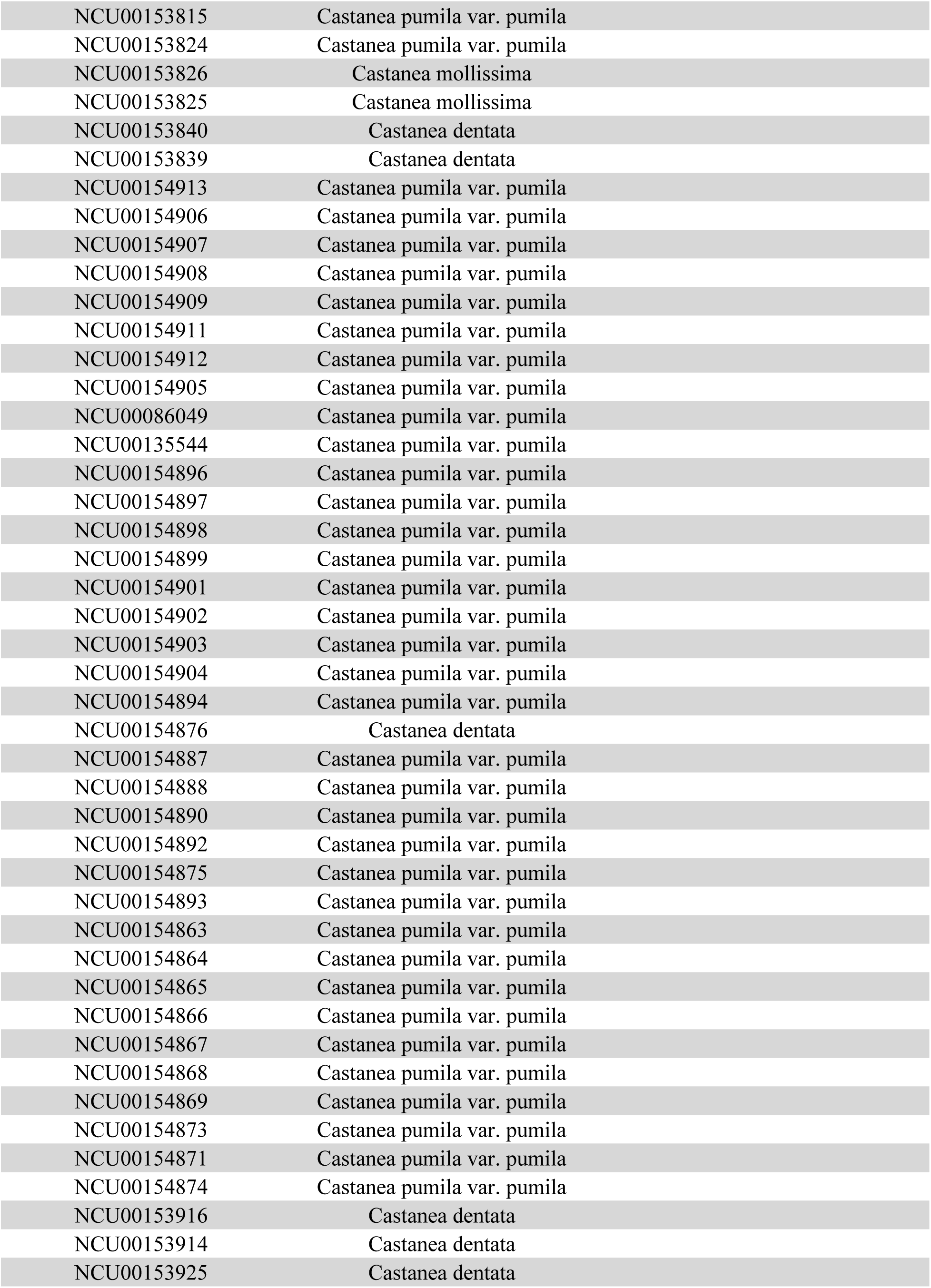
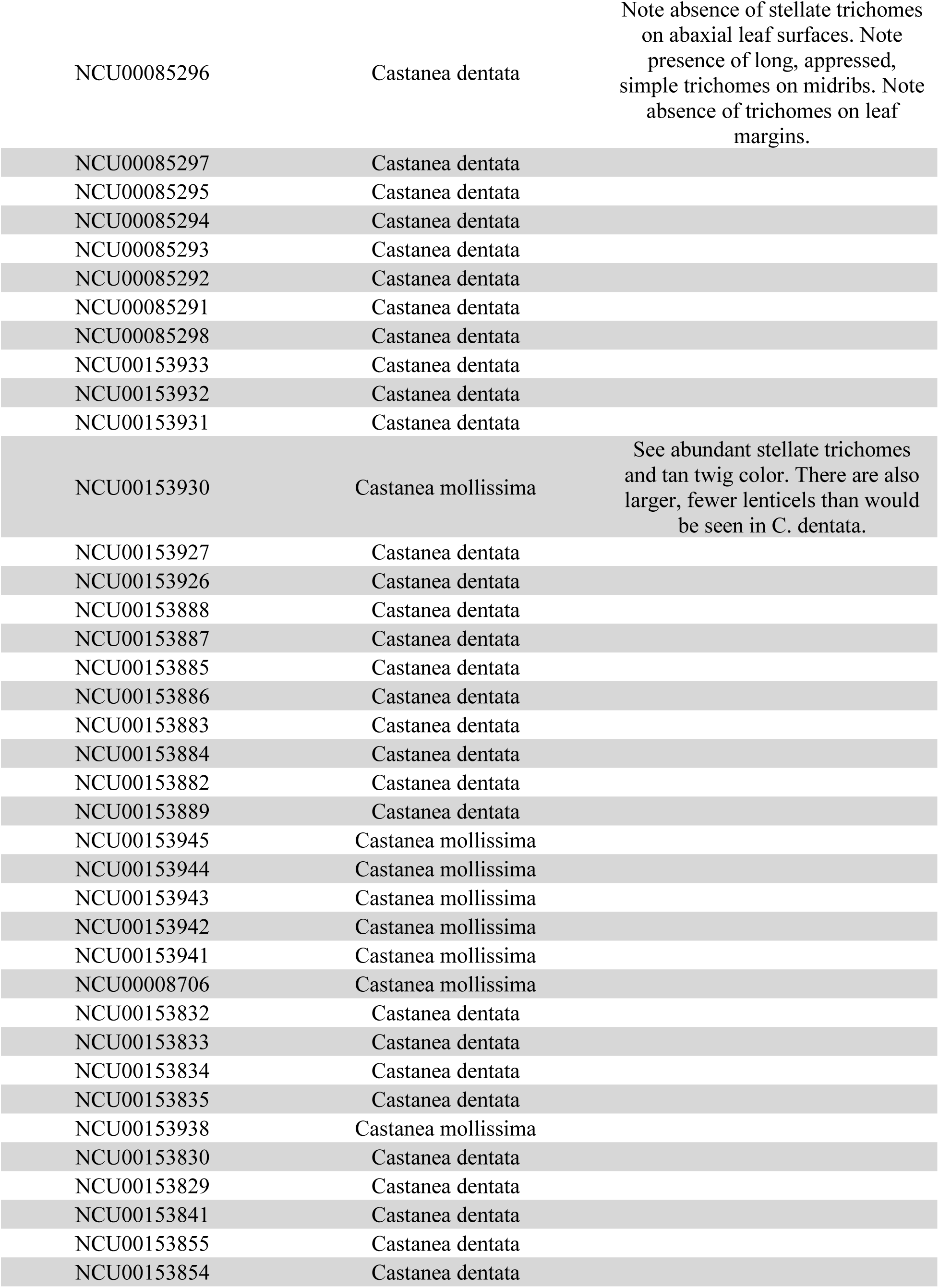
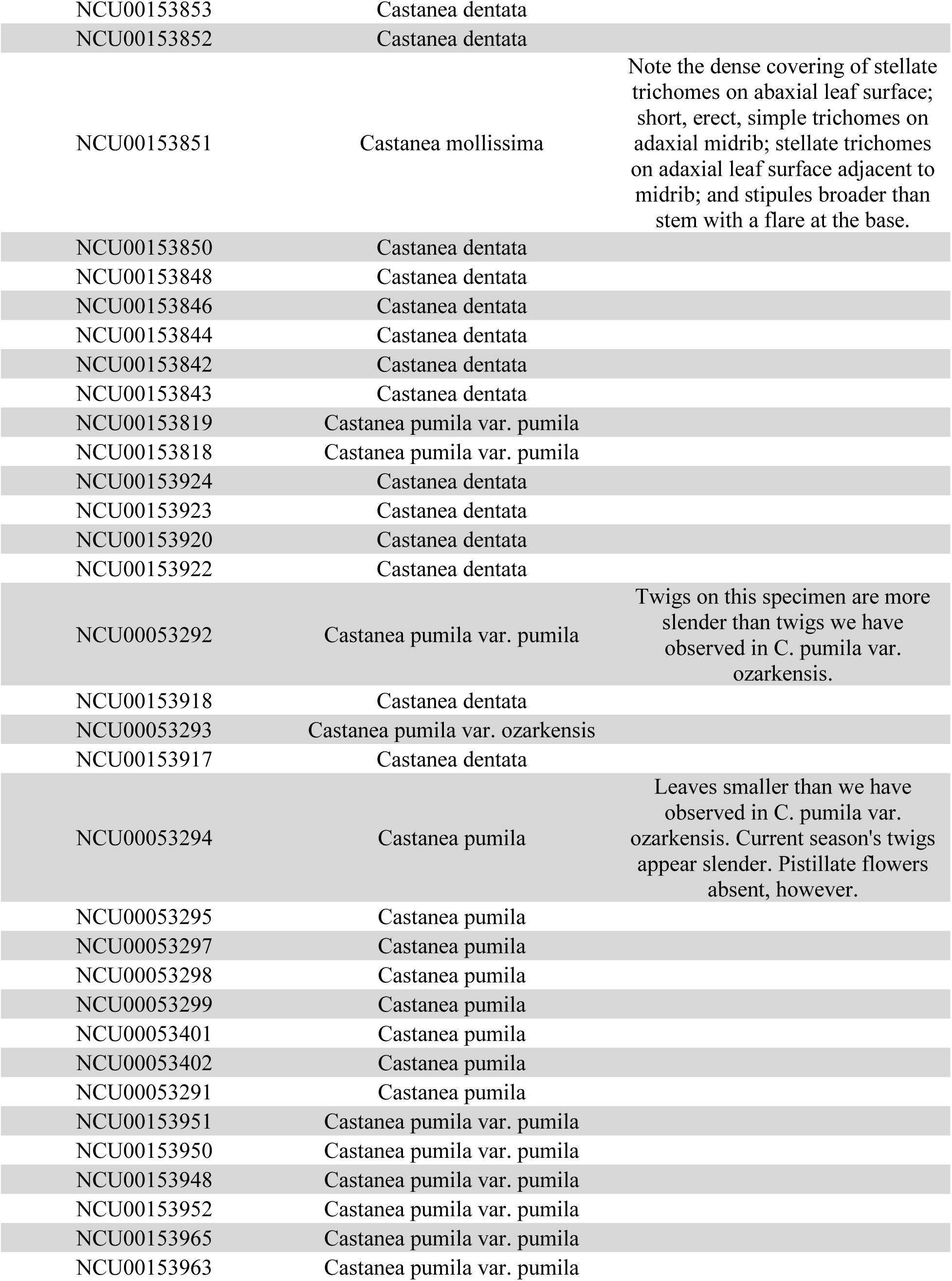
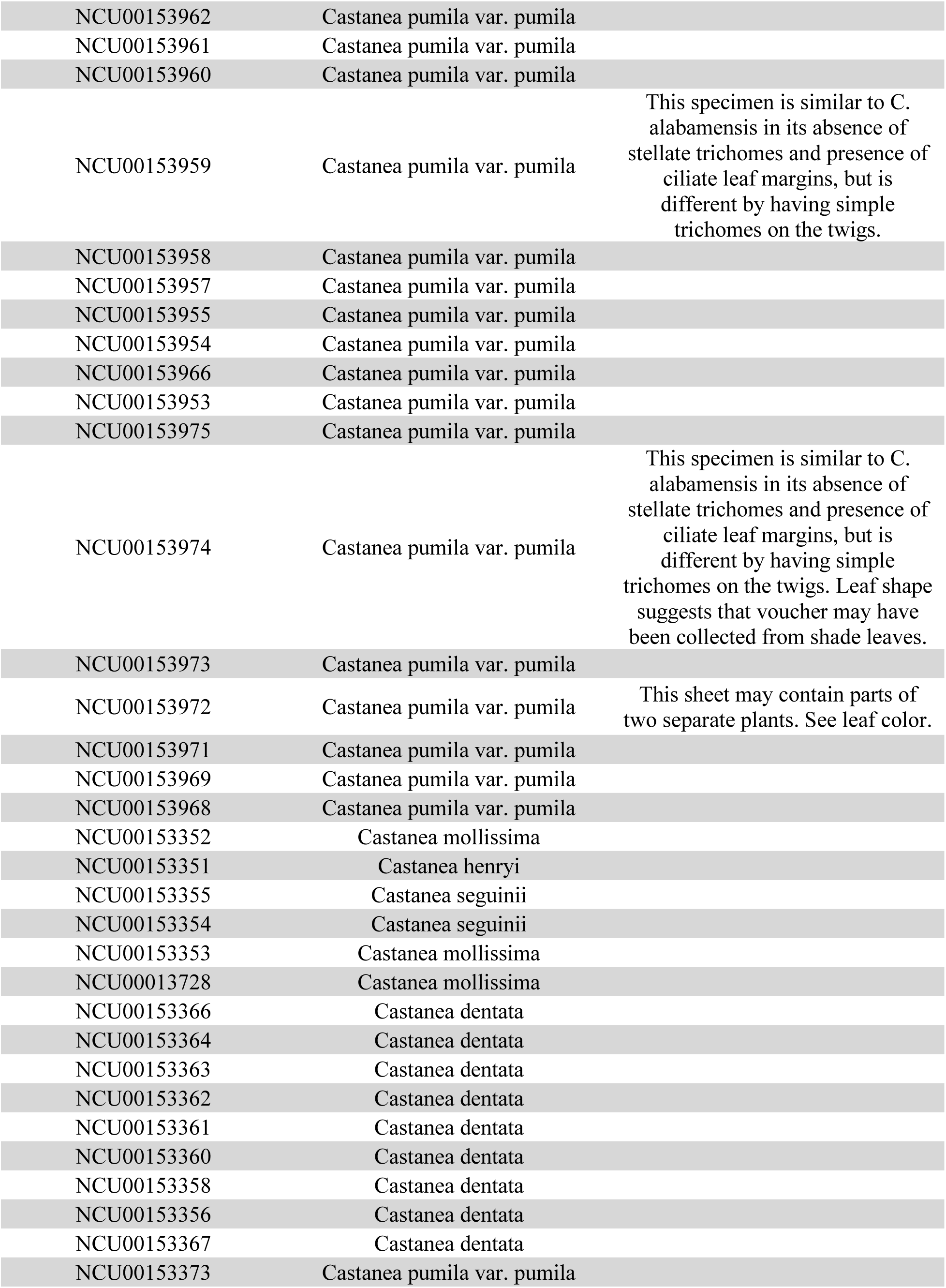
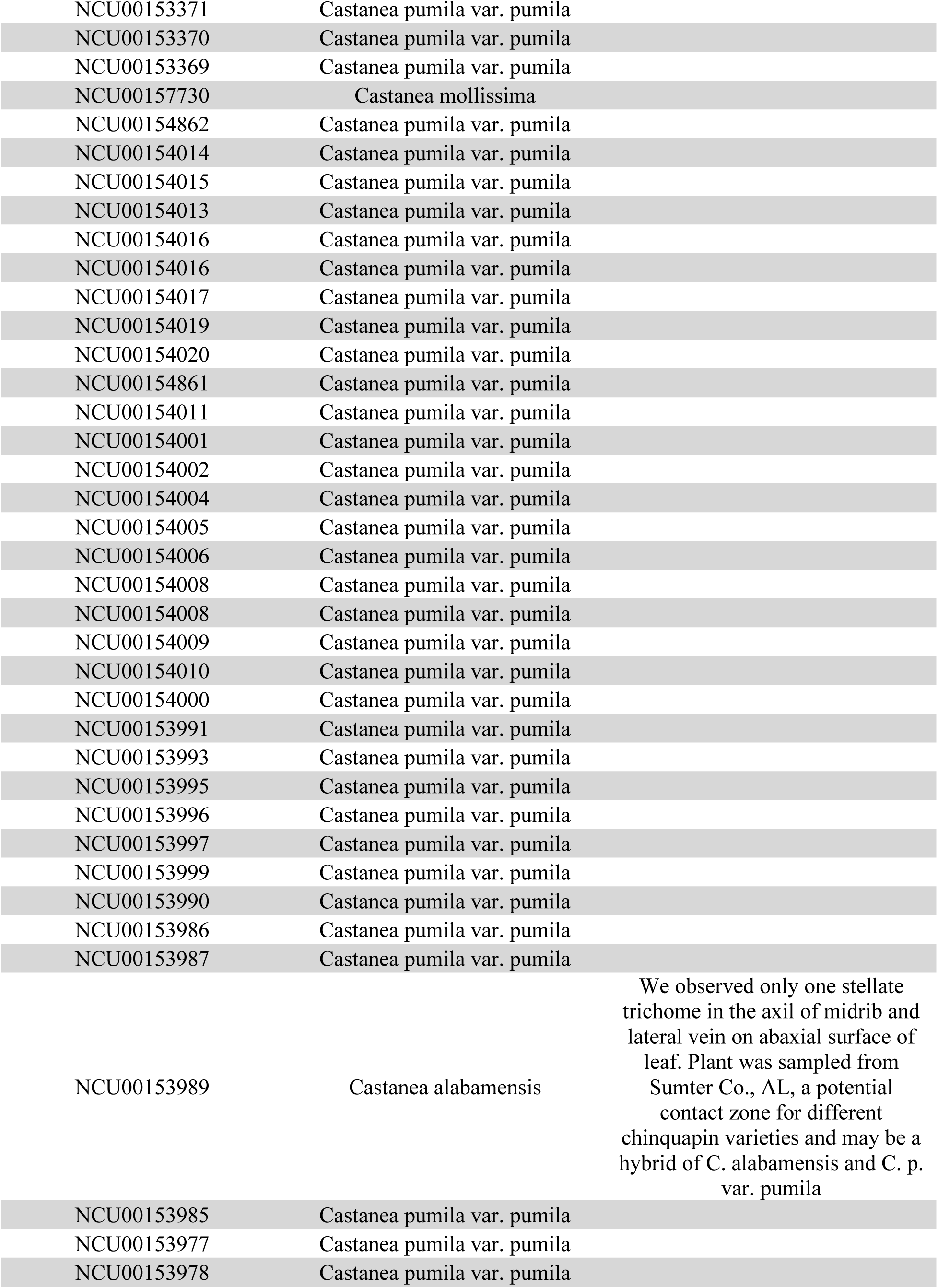
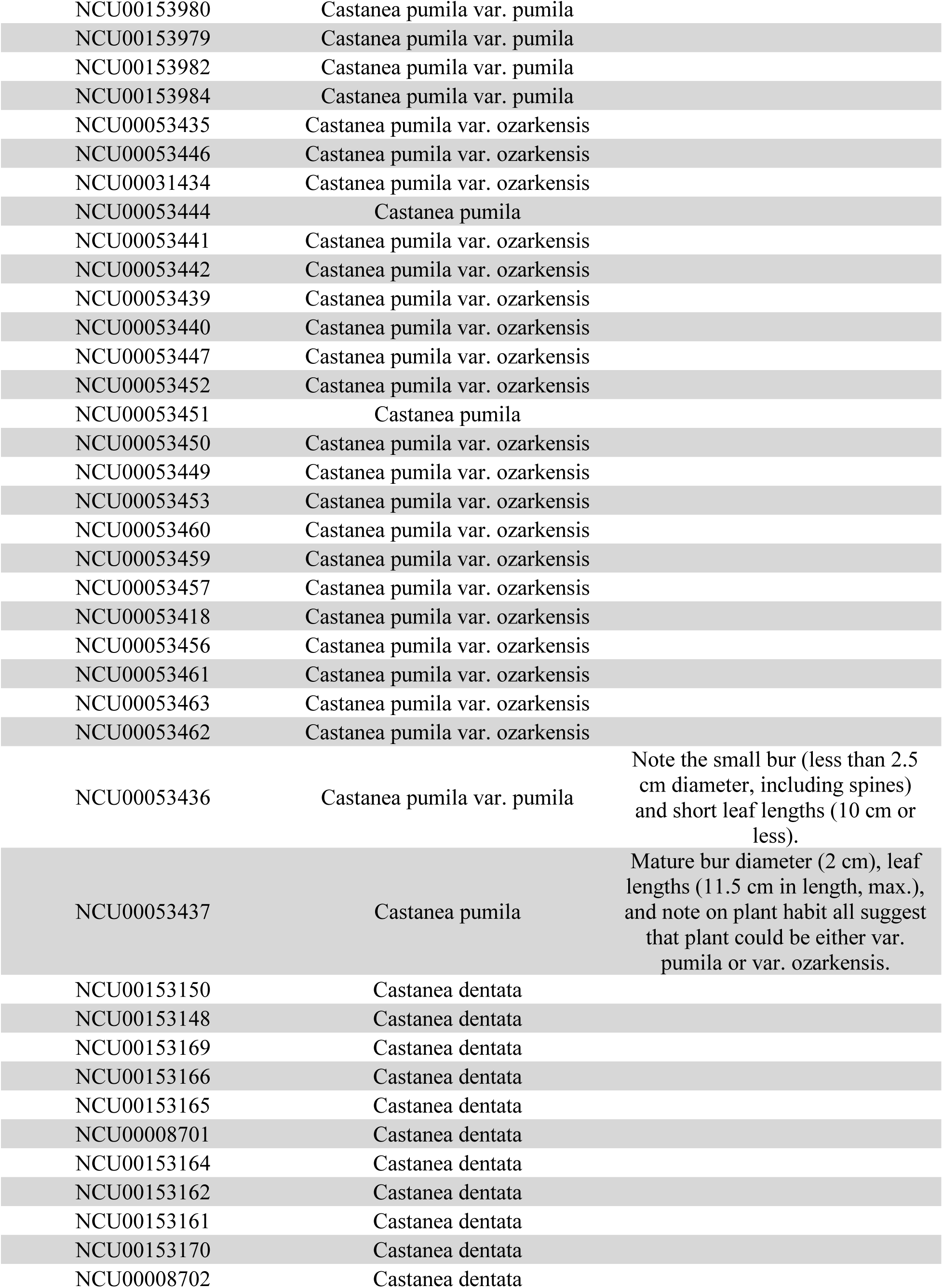
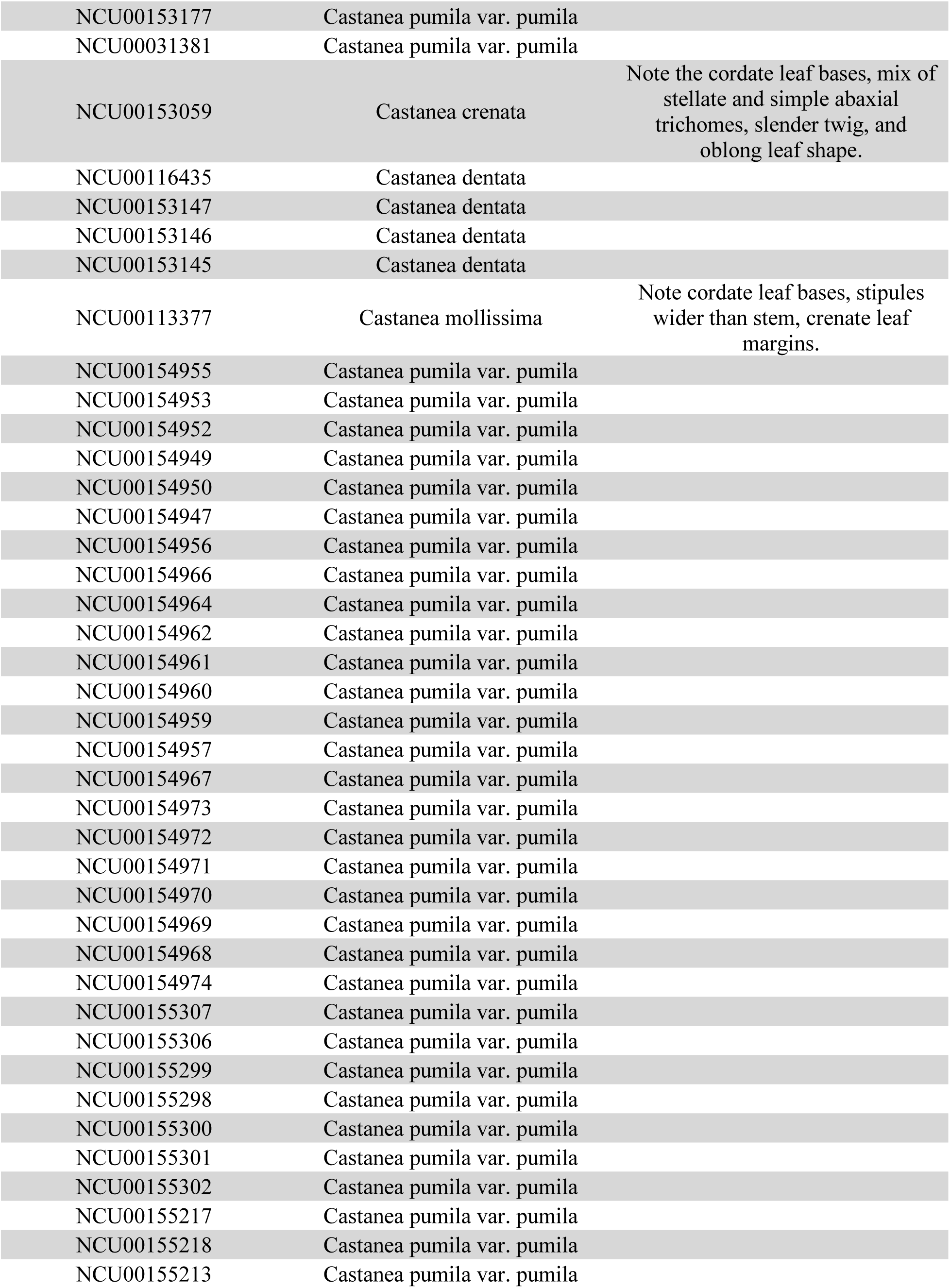
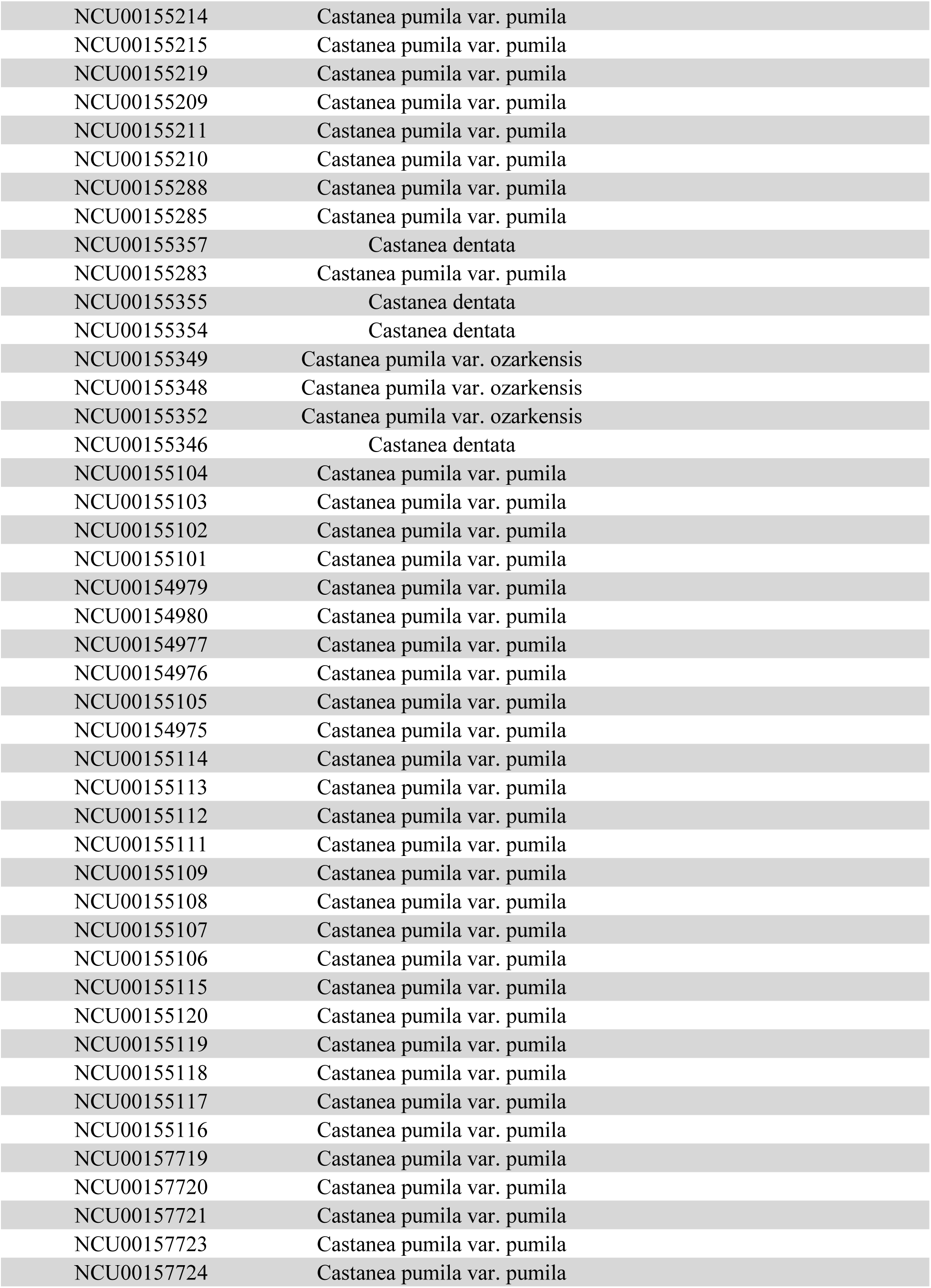
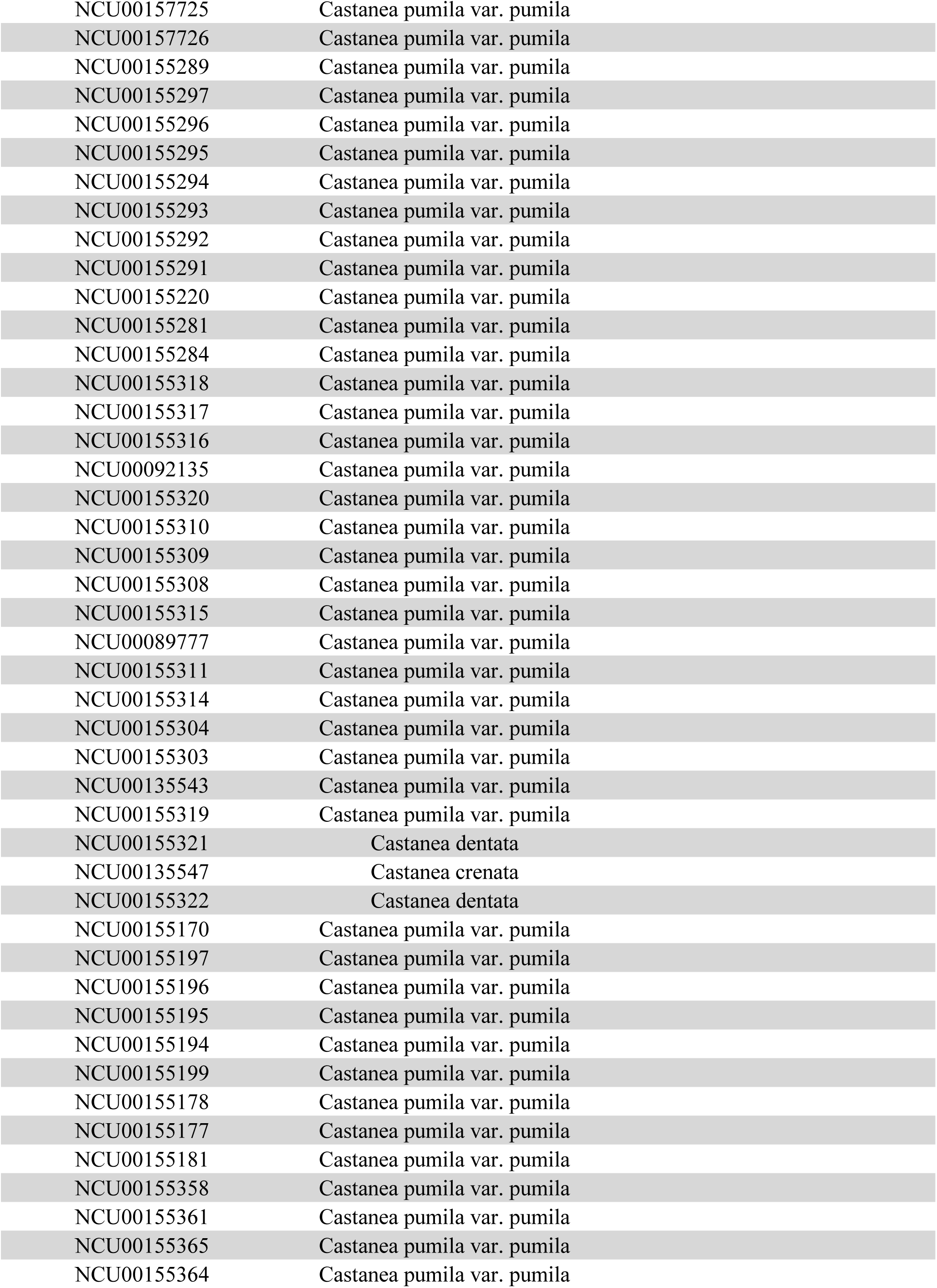
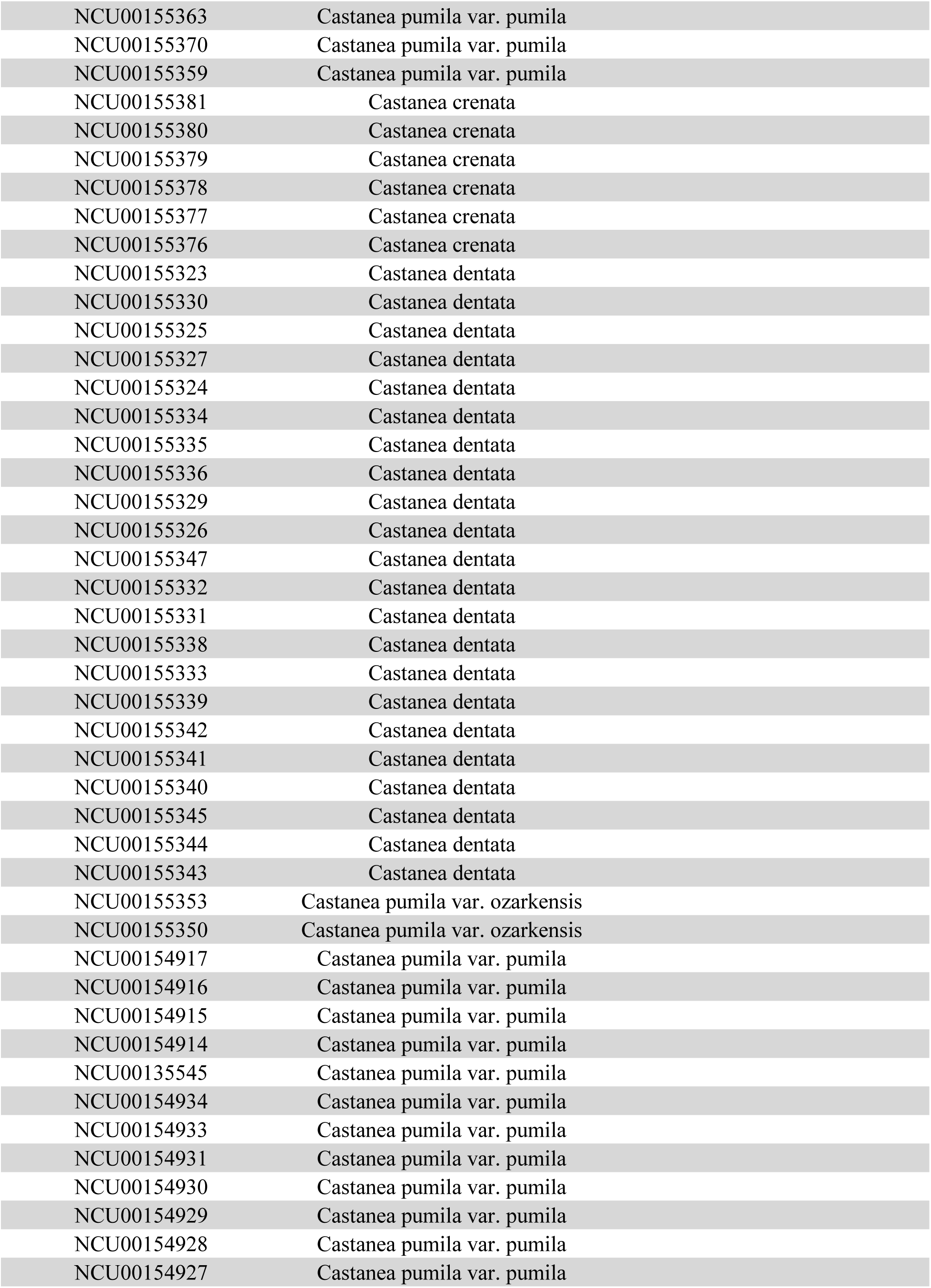
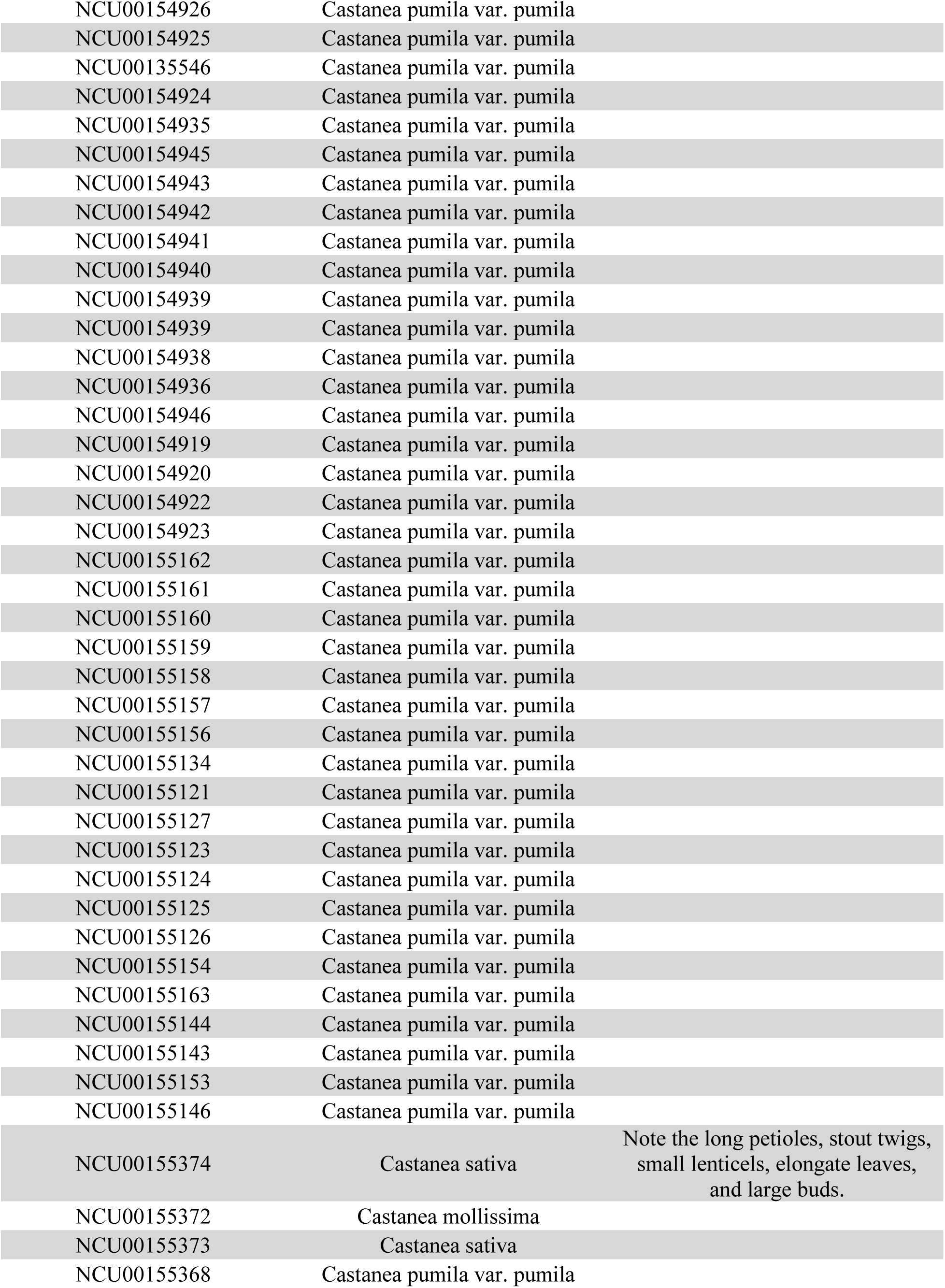
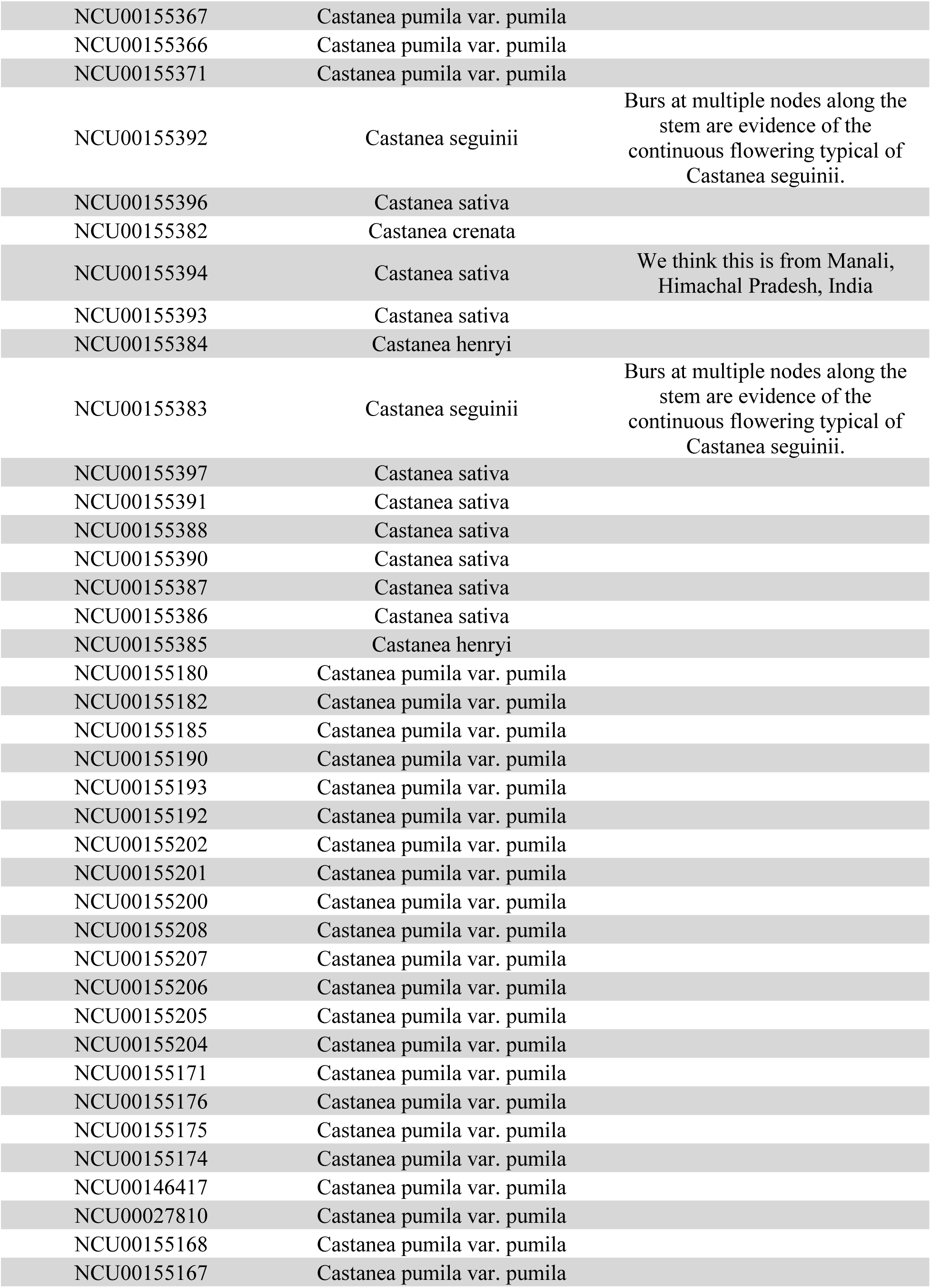
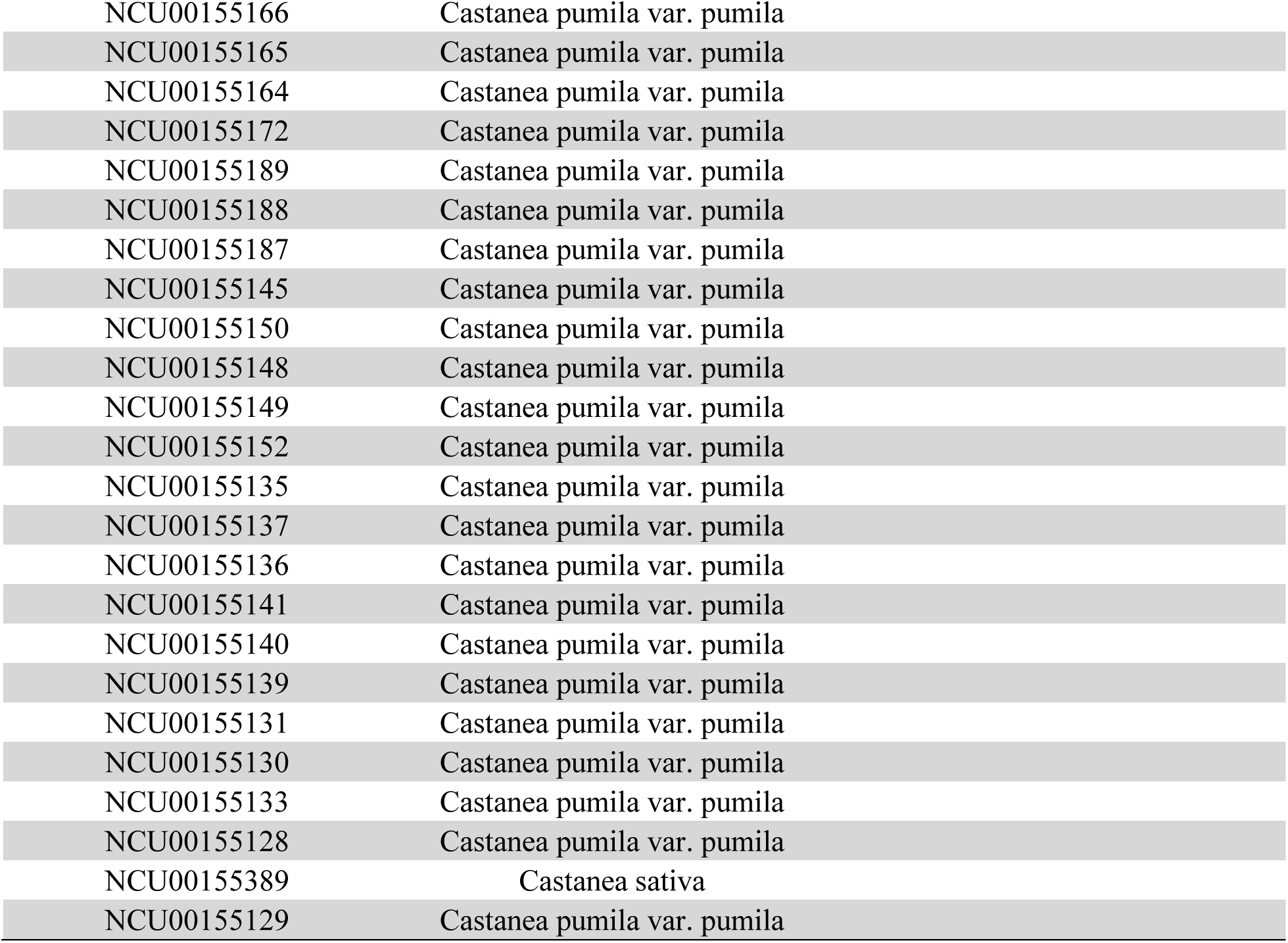
Herbarium accessions housed at the University of North Carolina Chapel Hill Herbarium (NCU) that were assessed for morphological characters and annotated. Vouchers collected by W.W. Ashe and used for his description of *C. alabamensis* are denoted with “collected by W.W. Ashe” in the identification remarks column. SERNEC catalog numbers are unique to each accession in the Southeast Regional Network of Expertise and Collections (SERNEC) project. Images of each accession, along with locality and collector information, can be accessed at the SERNEC website: http://sernecportal.org/portal/index.php.

| SERNEC catalog number | Taxonomic determination by Perkins et al. | Identification remarks |
| --- | --- | --- |
| NCU00153249 | *Castanea pumila* var. *pumila* |  |
| NCU00153250 | *Castanea pumila* var. *pumila* |  |
| NCU00153251 | *Castanea pumila* var. *pumila* |  |
| NCU00153252 | *Castanea pumila* var. *pumila* |  |
| NCU00153253 | *Castanea pumila* var. *pumila* |  |
| NCU00153254 | *Castanea pumila* var. *pumila* |  |
| NCU00153255 | *Castanea pumila* var. *pumila* |  |
| NCU00153248 | *Castanea pumila* var. *pumila* |  |
| NCU00153247 | *Castanea pumila* var. *pumila* |  |
| NCU00153241 | *Castanea pumila* var. *pumila* |  |
| NCU00153242 | *Castanea pumila* var. *pumila* |  |
| NCU00153243 | *Castanea pumila* var. *pumila* |  |
| NCU00153244 | *Castanea pumila* var. *pumila* |  |
| NCU00153246 | *Castanea pumila* var. *pumila* |  |
| NCU00153238 | *Castanea pumila* var. *pumila* |  |
| NCU00153239 | *Castanea pumila* var. *pumila* |  |
| NCU00153240 | *Castanea pumila* var. *pumila* |  |
| NCU00153231 | *Castanea pumila* var. *pumila* |  |
| NCU00153232 | *Castanea pumila* var. *pumila* |  |
| NCU00153233 | *Castanea pumila* var. *pumila* |  |
| NCU00153235 | *Castanea pumila* var. *pumila* |  |
| NCU00153237 | *Castanea pumila* var. *pumila* |  |
| NCU00027970 | *Castanea pumila* var. *pumila* |  |
| NCU00153211 | *Castanea pumila* var. *pumila* |  |
| NCU00153212 | *Castanea pumila* var. *pumila* |  |
| NCU00153213 | *Castanea pumila* var. *pumila* |  |
| NCU00113544 | *Castanea pumila* var. *pumila* |  |
| NCU00153215 | *Castanea pumila* var. *pumila* |  |
| NCU00153216 | *Castanea pumila* var. *pumila* |  |
| NCU00153217 | *Castanea pumila* var. *pumila* |  |
| NCU00153204 | *Castanea pumila* var. *pumila* |  |
| NCU00153199 | *Castanea pumila* var. *pumila* |  |
| NCU00153130 | *Castanea dentata* |  |
| NCU00153128 | *Castanea dentata* |  |
| NCU00153127 | *Castanea dentata* |  |
| NCU00153125 | Castanea dentata |  |
| NCU00153124 | Castanea dentata |  |
| NCU00153123 | Castanea dentata |  |
| NCU00153057 | Castanea dentata |  |
| NCU00153131 | Castanea dentata | Occasional simple trichome at leaf margin |
| NCU00153058 | Castanea dentata |  |
| NCU00153132 | Castanea dentata |  |
| NCU00153048 | Castanea dentata |  |
| NCU00027754 | Castanea dentata |  |
| NCU00153049 | Castanea dentata |  |
| NCU00145692 | Castanea dentata | Rare stellate trichome |
| NCU00153136 | Castanea dentata |  |
| NCU00153134 | Castanea dentata | Rare stellate trichome |
| NCU00153133 | Castanea dentata |  |
| NCU00153060 | Castanea dentata |  |
| NCU00153897 | Castanea dentata |  |
| NCU00153822 | Castanea pumila var. pumila |  |
| NCU00153823 | Castanea pumila var. pumila |  |
| NCU00153814 | Castanea pumila var. pumila |  |
| NCU00153809 | Castanea pumila var. pumila | Specimen has section of stolon |
| NCU00153811 | Castanea pumila var. pumila | But see Harbison's note re: "C. alnifolia past blooming while ashei is yet in full bloom..." |
| NCU00153798 | Castanea pumila var. pumila |  |
| NCU00153947 | Castanea pumila var. pumila |  |
| NCU00153800 | Castanea pumila var. pumila |  |
| NCU00153799 | Castanea pumila var. pumila |  |
| NCU00153794 | Castanea pumila var. pumila |  |
| NCU00153796 | Castanea pumila var. pumila |  |
| NCU00153804 | Castanea pumila var. pumila |  |
| NCU00153806 | Castanea pumila var. pumila |  |
| NCU00153813 | Castanea pumila var. pumila |  |
| NCU00153812 | Castanea pumila var. pumila |  |
| NCU00153802 | Castanea pumila var. pumila |  |
| NCU00153807 | Castanea pumila var. pumila |  |
| NCU00153144 | Castanea dentata |  |
| NCU00153143 | Castanea dentata |  |
| NCU00153140 | Castanea dentata |  |
| NCU00153139 | Castanea dentata |  |
| NCU00153138 | Castanea dentata |  |
| NCU00153160 | Castanea dentata |  |
| NCU00153209 | Castanea pumila var. pumila |  |
| NCU00153208 | Castanea pumila var. pumila |  |
| NCU00153207 | Castanea pumila var. pumila |  |

|  |  |
| --- | --- |
| NCU00153206 | Castanea pumila var. pumila |
| NCU00153205 | Castanea pumila var. pumila |
| NCU00153220 | Castanea pumila var. pumila |
| NCU00055657 | Castanea pumila var. pumila |
| NCU00153229 | Castanea pumila var. pumila |
| NCU00153228 | Castanea pumila var. pumila |
| NCU00153227 | Castanea pumila var. pumila |
| NCU00153225 | Castanea pumila var. pumila |
| NCU00153223 | Castanea pumila var. pumila |
| NCU00153222 | Castanea pumila var. pumila |
| NCU00153221 | Castanea pumila var. pumila |
| NCU00153192 | Castanea pumila var. pumila |
| NCU00008712 | Castanea pumila var. pumila |
| NCU00153190 | Castanea pumila var. pumila |
| NCU00093627 | Castanea pumila var. pumila |
| NCU00153189 | Castanea pumila var. pumila |
| NCU00153188 | Castanea pumila var. pumila |
| NCU00153187 | Castanea pumila var. pumila |
| NCU00153193 | Castanea pumila var. pumila |
| NCU00153198 | Castanea pumila var. pumila |
| NCU00153197 | Castanea pumila var. pumila |
| NCU00153196 | Castanea pumila var. pumila |
| NCU00153194 | Castanea pumila var. pumila |
| NCU00008710 | Castanea pumila var. pumila |
| NCU00153203 | Castanea pumila var. pumila |
| NCU00153202 | Castanea pumila var. pumila |
| NCU00153201 | Castanea pumila var. pumila |
| NCU00153792 | Castanea pumila var. pumila |
| NCU00153420 | Castanea pumila var. pumila |
| NCU00153781 | Castanea pumila var. pumila |
| NCU00153782 | Castanea pumila var. pumila |
| NCU00153784 | Castanea pumila var. pumila |
| NCU00153790 | Castanea pumila var. pumila |
| NCU00153788 | Castanea pumila var. pumila |
| NCU00153789 | Castanea pumila var. pumila |
| NCU00153787 | Castanea pumila var. pumila |
| NCU00153793 | Castanea pumila var. pumila |
| NCU00153867 | Castanea dentata |
| NCU00153866 | Castanea dentata |
| NCU00153864 | Castanea dentata |
| NCU00153862 | Castanea dentata |
| NCU00153861 | Castanea dentata |
| NCU00153860 | Castanea dentata |
| NCU00153859 | Castanea dentata |
| NCU00153403 | Castanea pumila var. pumila |
| NCU00153379 | Castanea pumila var. pumila |  |
| NCU00153946 | Castanea pumila var. pumila | Tips of leaves not acuminate, cupules four per spike. |
| NCU00153378 | Castanea pumila var. pumila |  |
| NCU00153377 | Castanea pumila var. pumila |  |
| NCU00153387 | Castanea pumila var. pumila |  |
| NCU00153376 | Castanea pumila var. pumila |  |
| NCU00153392 | Castanea pumila var. pumila |  |
| NCU00153391 | Castanea pumila var. pumila |  |
| NCU00153389 | Castanea pumila var. pumila |  |
| NCU00153401 | Castanea pumila var. pumila |  |
| NCU00153400 | Castanea pumila var. pumila |  |
| NCU00153399 | Castanea pumila var. pumila |  |
| NCU00153398 | Castanea pumila var. pumila |  |
| NCU00153396 | Castanea pumila var. pumila |  |
| NCU00153395 | Castanea pumila var. pumila |  |
| NCU00153393 | Castanea pumila var. pumila |  |
| NCU00153406 | Castanea pumila var. pumila |  |
| NCU00153175 | Castanea pumila var. pumila | Abaxial leaf surfaces densely covered with stellate trichomes. Deep sinuses along leaf margin. Our population genomic work demonstrates that plants with similar morphology from Blue Ridge region of NC are not hybrids. |
| NCU00153173 | Castanea pumila var. pumila | Leaf shape is lanceolate, like commonly seen in C. dentata, and abaxial surface is densely covered with stellate trichomes, like commonly seen in C. pumila. Yet our population genomic work demonstrates that plants with similar morphology are not hybrids. |
| NCU00153176 | Castanea x neglecta | Note mixture of stellate trichomes and long, simple trichomes on abaxial surface; long, adpressed, simple trichomes on leaf veins. |
| NCU00129230 | Castanea mollissima |  |
| NCU00153896 | Castanea dentata |  |
| NCU00153895 | Castanea mollissima |  |
| NCU00153894 | Castanea dentata |  |
| NCU00153893 | Castanea dentata |  |
| NCU00153892 | Castanea dentata |  |
| NCU00050100 | Castanea alabamensis | Collected by W.W. Ashe |
| NCU00053434 | Castanea alabamensis | Collected by W.W. Ashe |
| NCU00050202 | Castanea alabamensis |  |
| NCU00053426 | Castanea alabamensis | Collected by W.W. Ashe |
| NCU00053427 | Castanea alabamensis | Collected by W.W. Ashe |
| NCU00053423 | Castanea pumila var. pumila | Note 10 cupules per spike, slender twigs, and stellate interveinal trichomes on abaxial surface. |
| NCU00053421 | Castanea pumila var. pumila | Note stellate trichomes on interveinal lamina of abaxial surface. |
| NCU00053420 | Castanea alabamensis | Collected by W.W. Ashe |
| NCU00053410 | Castanea pumila var. ozarkensis |  |
| NCU00053415 | Castanea pumila var. ozarkensis |  |
| NCU00053404 | Castanea pumila var. ozarkensis |  |
| NCU00082746 | Castanea pumila var. ozarkensis |  |
| NCU00053424 | Castanea pumila var. pumila |  |
| NCU00053409 | Castanea pumila var. ozarkensis |  |
| NCU00053416 | Castanea pumila var. pumila | See stellate trichomes on abaxial lamina. |
| NCU00053425 | Castanea pumila var. pumila | See stellate trichomes on abaxial lamina, slender twigs, and four cupules on spike. |
| NCU00053417 | Castanea pumila var. pumila | See stellate trichomes on abaxial lamina. |
| NCU00053411 | Castanea pumila var. ozarkensis |  |
| NCU00053408 | Castanea pumila var. ozarkensis |  |
| NCU00053414 | Castanea pumila var. ozarkensis |  |
| NCU00053403 | Castanea pumila var. ozarkensis | Has stellate trichomes on lamina surface. Has simple adpressed trichomes on veins. On young, unexpanded leaves, it appears that abaxial lamina has simple trichomes, but these are actually on veins. |
| NCU00053407 | Castanea pumila var. ozarkensis |  |
| NCU00053412 | Castanea pumila var. ozarkensis |  |
| NCU00053406 | Castanea pumila var. ozarkensis |  |
| NCU00053405 | Castanea pumila var. ozarkensis |  |
| NCU00053431 | Castanea alabamensis | Collected by W.W. Ashe. Note complete absence of stellate trichomes on abaxial lamina. Note ciliate leaf margin. Note glabrous, glaucous abaxial lamina. |
| NCU00053429 | Castanea alabamensis | Collected by Mrs. H.J. McDowell for W.W. Ashe. Note ciliate leaf margin, absence of trichomes on abaxial lamina, presence of simple trichomes on veins only. |
| NCU00053428 | Castanea alabamensis | Collected by W.W. Ashe. Note presence of leaf trichomes only on blade margin and veins. Complete absence of stellate trichomes on leaves. |
| NCU00053433 | Castanea alabamensis | Collected by W.W. Ashe |
| NCU00050092 | Castanea alabamensis | Collected by Mrs. H.J. McDowell for W.W. Ashe |
| NCU00050093 | Castanea alabamensis | Collected by Mrs. H.J. McDowell for W.W. Ashe |
| NCU00050094 | Castanea alabamensis | Collected by W.W. Ashe. Ciliate leaf margin. No stellate trichomes on abaxial lamina. Simple trichomes on veins of abaxial leaf surface. |
| NCU00050095 | Castanea alabamensis | Collected by W.W. Ashe |
| NCU00050096 | Castanea alabamensis | Collected by W.W. Ashe |
| NCU00053432 | Castanea alabamensis | Collected by W.W. Ashe |
| NCU00050097 | Castanea pumila var. pumila | Note stellate trichomes on abaxial lamina of leaves from both plants represented on this sheet. |
| NCU00050098 | Castanea pumila var. pumila | Note stellate trichomes on abaxial lamina. |
| NCU00153328 | Castanea pumila var. pumila |  |
| NCU00085046 | Castanea pumila var. pumila |  |
| NCU00153327 | Castanea pumila var. pumila |  |
| NCU00153326 | Castanea pumila var. pumila |  |
| NCU00153325 | Castanea pumila var. pumila |  |
| NCU00153323 | Castanea pumila var. pumila |  |
| NCU00153322 | Castanea pumila var. pumila |  |
| NCU00153321 | Castanea pumila var. pumila |  |
| NCU00153891 | Castanea dentata |  |
| NCU00153890 | Castanea dentata |  |
| NCU00153936 | Castanea mollissima |  |
| NCU00153935 | Castanea mollissima |  |
| NCU00153934 | Castanea mollissima |  |
| NCU00008707 | Castanea mollissima |  |
| NCU00153937 | Castanea mollissima |  |
| NCU00153940 | Castanea mollissima |  |
| NCU00153939 | Castanea mollissima |  |
| NCU00008708 | Castanea mollissima |  |
| NCU00153838 | Castanea dentata |  |
| NCU00153837 | Castanea dentata |  |
| NCU00153836 | Castanea dentata |  |
| NCU00153786 | Castanea pumila var. pumila |  |
| NCU00153785 | Castanea pumila var. pumila |  |
| NCU00153858 | Castanea dentata |  |
| NCU00153857 | Castanea dentata |  |
| NCU00153856 | Castanea dentata |  |
| NCU00153881 | Castanea dentata |  |
| NCU00153880 | Castanea dentata |  |
| NCU00153879 | Castanea dentata |  |
| NCU00153878 | Castanea dentata |  |
| NCU00153876 | Castanea dentata |  |
| NCU00153874 | Castanea dentata |  |
| NCU00153872 | Castanea pumila var. pumila | Note the abundant stellate trichomes on abaxial leaf surface, erect simple trichomes on abaxial midrib, and ciliate leaf margin. |
| NCU00153874 | Castanea dentata |  |
| NCU00153871 | Castanea dentata |  |
| NCU00153870 | Castanea dentata |  |
| NCU00153869 | Castanea dentata |  |
| NCU00153868 | Castanea dentata |  |
| NCU00153908 | Castanea dentata |  |
| NCU00153907 | Castanea dentata |  |
| NCU00153906 | Castanea dentata |  |
| NCU00153905 | Castanea dentata |  |
| NCU00153904 | Castanea dentata |  |
| NCU00140424 | Castanea alabamensis | Note ciliate leaf margin and abundant, erect, simple trichomes on leaf abaxial veins. |
| NCU00153903 | Castanea alabamensis | Note ciliate leaf margin and abundant, erect, simple trichomes on leaf abaxial veins. |
| NCU00153901 | Castanea alabamensis | Note ciliate leaf margin and abundant, erect, simple trichomes on leaf abaxial veins. |
| NCU00153900 | Castanea dentata | Note eciliate leaf margin, abundant glandular trichomes on abaxial leaf surface, and occasional appressed trichome on abaxial leaf veins. |
| NCU00153899 | Castanea dentata | Although simple trichomes are infrequently present on leaf margin, there are far fewer trichomes here than is found C. alabamensis. Also note long, typically appressed trichomes on abaxial leaf veins and lack of erect, simple trichomes on abaxial leaf surface. |
| NCU00153898 | Castanea dentata |  |
| NCU00153909 | Castanea dentata | Note frequent glandular trichomes on abaxial leaf surface, eciliate leaf margin, and long, often appressed, simple trichomes on abaxial leaf midrib. |
| NCU00153912 | Castanea dentata |  |
| NCU00153911 | Castanea dentata |  |
| NCU00153910 | Castanea dentata |  |
| NCU00153259 | Castanea pumila var. pumila |  |
| NCU00153258 | Castanea pumila var. pumila |  |
| NCU00153257 | Castanea pumila var. pumila |  |
| NCU00153263 | Castanea pumila var. pumila |  |
| NCU00153260 | Castanea pumila var. pumila |  |
| NCU00153270 | Castanea pumila var. pumila |  |
| NCU00153268 | Castanea pumila var. pumila |  |
| NCU00153266 | Castanea pumila var. pumila |  |
| NCU00153264 | Castanea pumila var. pumila |  |
| NCU00153414 | Castanea pumila var. pumila |  |
| NCU00153278 | Castanea pumila var. pumila |  |
| NCU00153277 | Castanea pumila var. pumila |  |
| NCU00116436 | Castanea pumila var. pumila |  |
| NCU00116437 | Castanea pumila var. pumila |  |
| NCU00153271 | Castanea pumila var. pumila |  |
| NCU00153281 | Castanea pumila var. pumila |  |
| NCU00153282 | Castanea pumila var. pumila |  |
| NCU00153280 | Castanea pumila var. pumila |  |
| NCU00153287 | Castanea pumila var. pumila |  |
| NCU00153285 | Castanea pumila var. pumila |  |
| NCU00153284 | Castanea pumila var. pumila |  |
| NCU00075919 | Castanea pumila var. pumila |  |
| NCU00153291 | Castanea pumila var. pumila |  |
| NCU00153290 | Castanea pumila var. pumila |  |
| NCU00153288 | Castanea pumila var. pumila |  |
| NCU00153149 | Castanea dentata |  |
| NCU00153151 | Castanea dentata |  |
| NCU00153152 | Castanea dentata |  |
| NCU00153153 | Castanea dentata |  |
| NCU00153154 | Castanea dentata |  |
| NCU00135776 | Castanea pumila var. pumila | Note abundant stellate trichomes on abaxial leaf surface. |
| NCU00153157 | Castanea dentata |  |
| NCU00153158 | Castanea dentata |  |
| NCU00153159 | Castanea dentata |  |
| NCU00153185 | Castanea pumila var. pumila |  |
| NCU00153184 | Castanea pumila var. pumila |  |

|  |  |
| --- | --- |
| NCU00153179 | Castanea pumila var. pumila |
| NCU00153181 | Castanea pumila var. pumila |
| NCU00153183 | Castanea pumila var. pumila |
| NCU00008711 | Castanea pumila var. pumila |
| NCU00153121 | Castanea dentata |
| NCU00008703 | Castanea dentata |
| NCU00153122 | Castanea dentata |
| NCU00153047 | Castanea crenata |
| NCU00008705 | Castanea dentata |
| NCU00153051 | Castanea dentata |
| NCU00153052 | Castanea dentata |
| NCU00153054 | Castanea dentata |
| NCU00153053 | Castanea dentata |
| NCU00153056 | Castanea dentata |
| NCU00153055 | Castanea dentata |
| NCU00153336 | Castanea pumila var. pumila |
| NCU00153337 | Castanea pumila var. pumila |
| NCU00153339 | Castanea pumila var. pumila |
| NCU00153338 | Castanea pumila var. pumila |
| NCU00153340 | Castanea pumila var. pumila |
| NCU00014050 | Castanea pumila var. pumila |
| NCU00153341 | Castanea pumila var. pumila |
| NCU00153342 | Castanea pumila var. pumila |
| NCU00153344 | Castanea pumila var. pumila |
| NCU00153345 | Castanea pumila var. pumila |
| NCU00153346 | Castanea pumila var. pumila |
| NCU00153348 | Castanea pumila var. pumila |
| NCU00153335 | Castanea pumila var. pumila |
| NCU00153334 | Castanea pumila var. pumila |
| NCU00153333 | Castanea pumila var. pumila |
| NCU00153332 | Castanea pumila var. pumila |
| NCU00153331 | Castanea pumila var. pumila |
| NCU00115121 | Castanea pumila var. pumila |
| NCU00153329 | Castanea pumila var. pumila |
| NCU00153178 | Castanea pumila var. pumila |
| NCU00153304 | Castanea pumila var. pumila |
| NCU00153294 | Castanea pumila var. pumila |
| NCU00153295 | Castanea pumila var. pumila |
| NCU00153297 | Castanea pumila var. pumila |
| NCU00028020 | Castanea pumila var. pumila |
| NCU00153298 | Castanea pumila var. pumila |
| NCU00008709 | Castanea pumila var. pumila |
| NCU00153299 | Castanea pumila var. pumila |
| NCU00153300 | Castanea pumila var. pumila |
| NCU00153293 | Castanea pumila var. pumila |
| NCU00153320 | Castanea pumila var. pumila |  |
| NCU00153305 | Castanea pumila var. pumila |  |
| NCU00153306 | Castanea pumila var. pumila |  |
| NCU00153307 | Castanea pumila var. pumila |  |
| NCU00153310 | Castanea pumila var. pumila |  |
| NCU00153309 | Castanea pumila var. pumila |  |
| NCU00153312 | Castanea pumila var. pumila |  |
| NCU00153313 | See note | Specimens from two different species are glued to this sheet. A = C. dentata; B = C. pumila var. pumila. |
| NCU00153313 | See note | Specimens from two different species are glued to this sheet. A = C. dentata; B = C. pumila var. pumila. |
| NCU00153314 | Castanea pumila var. pumila |  |
| NCU00153315 | Castanea pumila var. pumila |  |
| NCU00153317 | Castanea pumila var. pumila |  |
| NCU00153319 | Castanea pumila var. pumila |  |
| NCU00153318 | Castanea pumila var. pumila |  |
| NCU00153350 | Castanea mollissima |  |
| NCU00110700 | Castanea pumila var. pumila |  |
| NCU00153349 | Castanea pumila var. pumila |  |
| NCU00153402 | Castanea pumila var. pumila |  |
| NCU00153404 | Castanea pumila var. pumila |  |
| NCU00153407 | Castanea pumila var. pumila |  |
| NCU00153418 | Castanea pumila var. pumila |  |
| NCU00153416 | Castanea pumila var. pumila |  |
| NCU00153413 | Castanea pumila var. pumila |  |
| NCU00153412 | Castanea pumila var. pumila |  |
| NCU00153411 | Castanea pumila var. pumila |  |
| NCU00153410 | Castanea pumila var. pumila |  |
| NCU00153409 | Castanea pumila var. pumila |  |
| NCU00153408 | Castanea pumila var. pumila |  |
| NCU00153419 | Castanea pumila var. pumila |  |
| NCU00153368 | Castanea pumila var. pumila |  |
| NCU00153374 | Castanea pumila var. pumila |  |
| NCU00153375 | Castanea pumila var. pumila |  |
| NCU00153386 | Castanea pumila var. pumila |  |
| NCU00153385 | Castanea pumila var. pumila |  |
| NCU00153382 | Castanea pumila var. pumila |  |
| NCU00153381 | Castanea pumila var. pumila |  |
| NCU00153913 | Castanea dentata |  |
| NCU00153817 | Castanea pumila var. pumila |  |
| NCU00153816 | Castanea pumila var. pumila |  |

|  |  |
| --- | --- |
| NCU00153815 | Castanea pumila var. pumila |
| NCU00153824 | Castanea pumila var. pumila |
| NCU00153826 | Castanea mollissima |
| NCU00153825 | Castanea mollissima |
| NCU00153840 | Castanea dentata |
| NCU00153839 | Castanea dentata |
| NCU00154913 | Castanea pumila var. pumila |
| NCU00154906 | Castanea pumila var. pumila |
| NCU00154907 | Castanea pumila var. pumila |
| NCU00154908 | Castanea pumila var. pumila |
| NCU00154909 | Castanea pumila var. pumila |
| NCU00154911 | Castanea pumila var. pumila |
| NCU00154912 | Castanea pumila var. pumila |
| NCU00154905 | Castanea pumila var. pumila |
| NCU00086049 | Castanea pumila var. pumila |
| NCU00135544 | Castanea pumila var. pumila |
| NCU00154896 | Castanea pumila var. pumila |
| NCU00154897 | Castanea pumila var. pumila |
| NCU00154898 | Castanea pumila var. pumila |
| NCU00154899 | Castanea pumila var. pumila |
| NCU00154901 | Castanea pumila var. pumila |
| NCU00154902 | Castanea pumila var. pumila |
| NCU00154903 | Castanea pumila var. pumila |
| NCU00154904 | Castanea pumila var. pumila |
| NCU00154894 | Castanea pumila var. pumila |
| NCU00154876 | Castanea dentata |
| NCU00154887 | Castanea pumila var. pumila |
| NCU00154888 | Castanea pumila var. pumila |
| NCU00154890 | Castanea pumila var. pumila |
| NCU00154892 | Castanea pumila var. pumila |
| NCU00154875 | Castanea pumila var. pumila |
| NCU00154893 | Castanea pumila var. pumila |
| NCU00154863 | Castanea pumila var. pumila |
| NCU00154864 | Castanea pumila var. pumila |
| NCU00154865 | Castanea pumila var. pumila |
| NCU00154866 | Castanea pumila var. pumila |
| NCU00154867 | Castanea pumila var. pumila |
| NCU00154868 | Castanea pumila var. pumila |
| NCU00154869 | Castanea pumila var. pumila |
| NCU00154873 | Castanea pumila var. pumila |
| NCU00154871 | Castanea pumila var. pumila |
| NCU00154874 | Castanea pumila var. pumila |
| NCU00153916 | Castanea dentata |
| NCU00153914 | Castanea dentata |
| NCU00153925 | Castanea dentata |
| NCU00085296 | Castanea dentata | Note absence of stellate trichomes on abaxial leaf surfaces. Note presence of long, appressed, simple trichomes on midribs. Note absence of trichomes on leaf margins. |
| NCU00085297 | Castanea dentata |  |
| NCU00085295 | Castanea dentata |  |
| NCU00085294 | Castanea dentata |  |
| NCU00085293 | Castanea dentata |  |
| NCU00085292 | Castanea dentata |  |
| NCU00085291 | Castanea dentata |  |
| NCU00085298 | Castanea dentata |  |
| NCU00153933 | Castanea dentata |  |
| NCU00153932 | Castanea dentata |  |
| NCU00153931 | Castanea dentata |  |
| NCU00153930 | Castanea mollissima | See abundant stellate trichomes and tan twig color. There are also larger, fewer lenticels than would be seen in C. dentata. |
| NCU00153927 | Castanea dentata |  |
| NCU00153926 | Castanea dentata |  |
| NCU00153888 | Castanea dentata |  |
| NCU00153887 | Castanea dentata |  |
| NCU00153885 | Castanea dentata |  |
| NCU00153886 | Castanea dentata |  |
| NCU00153883 | Castanea dentata |  |
| NCU00153884 | Castanea dentata |  |
| NCU00153882 | Castanea dentata |  |
| NCU00153889 | Castanea dentata |  |
| NCU00153945 | Castanea mollissima |  |
| NCU00153944 | Castanea mollissima |  |
| NCU00153943 | Castanea mollissima |  |
| NCU00153942 | Castanea mollissima |  |
| NCU00153941 | Castanea mollissima |  |
| NCU00008706 | Castanea mollissima |  |
| NCU00153832 | Castanea dentata |  |
| NCU00153833 | Castanea dentata |  |
| NCU00153834 | Castanea dentata |  |
| NCU00153835 | Castanea dentata |  |
| NCU00153938 | Castanea mollissima |  |
| NCU00153830 | Castanea dentata |  |
| NCU00153829 | Castanea dentata |  |
| NCU00153841 | Castanea dentata |  |
| NCU00153855 | Castanea dentata |  |
| NCU00153854 | Castanea dentata |  |
| NCU00153853 | Castanea dentata |  |
| NCU00153852 | Castanea dentata |  |
| NCU00153851 | Castanea mollissima | Note the dense covering of stellate trichomes on abaxial leaf surface; short, erect, simple trichomes on adaxial midrib; stellate trichomes on adaxial leaf surface adjacent to midrib; and stipules broader than stem with a flare at the base. |
| NCU00153850 | Castanea dentata |  |
| NCU00153848 | Castanea dentata |  |
| NCU00153846 | Castanea dentata |  |
| NCU00153844 | Castanea dentata |  |
| NCU00153842 | Castanea dentata |  |
| NCU00153843 | Castanea dentata |  |
| NCU00153819 | Castanea pumila var. pumila |  |
| NCU00153818 | Castanea pumila var. pumila |  |
| NCU00153924 | Castanea dentata |  |
| NCU00153923 | Castanea dentata |  |
| NCU00153920 | Castanea dentata |  |
| NCU00153922 | Castanea dentata |  |
| NCU00053292 | Castanea pumila var. pumila | Twigs on this specimen are more slender than twigs we have observed in C. pumila var. ozarkensis. |
| NCU00153918 | Castanea dentata |  |
| NCU00053293 | Castanea pumila var. ozarkensis |  |
| NCU00153917 | Castanea dentata |  |
| NCU00053294 | Castanea pumila | Leaves smaller than we have observed in C. pumila var. ozarkensis. Current season's twigs appear slender. Pistillate flowers absent, however. |
| NCU00053295 | Castanea pumila |  |
| NCU00053297 | Castanea pumila |  |
| NCU00053298 | Castanea pumila |  |
| NCU00053299 | Castanea pumila |  |
| NCU00053401 | Castanea pumila |  |
| NCU00053402 | Castanea pumila |  |
| NCU00053291 | Castanea pumila |  |
| NCU00153951 | Castanea pumila var. pumila |  |
| NCU00153950 | Castanea pumila var. pumila |  |
| NCU00153948 | Castanea pumila var. pumila |  |
| NCU00153952 | Castanea pumila var. pumila |  |
| NCU00153965 | Castanea pumila var. pumila |  |
| NCU00153963 | Castanea pumila var. pumila |  |
| NCU00153962 | Castanea pumila var. pumila |  |
| NCU00153961 | Castanea pumila var. pumila |  |
| NCU00153960 | Castanea pumila var. pumila |  |
| NCU00153959 | Castanea pumila var. pumila | This specimen is similar to C. alabamensis in its absence of stellate trichomes and presence of ciliate leaf margins, but is different by having simple trichomes on the twigs. |
| NCU00153958 | Castanea pumila var. pumila |  |
| NCU00153957 | Castanea pumila var. pumila |  |
| NCU00153955 | Castanea pumila var. pumila |  |
| NCU00153954 | Castanea pumila var. pumila |  |
| NCU00153966 | Castanea pumila var. pumila |  |
| NCU00153953 | Castanea pumila var. pumila |  |
| NCU00153975 | Castanea pumila var. pumila |  |
| NCU00153974 | Castanea pumila var. pumila | This specimen is similar to C. alabamensis in its absence of stellate trichomes and presence of ciliate leaf margins, but is different by having simple trichomes on the twigs. Leaf shape suggests that voucher may have been collected from shade leaves. |
| NCU00153973 | Castanea pumila var. pumila |  |
| NCU00153972 | Castanea pumila var. pumila | This sheet may contain parts of two separate plants. See leaf color. |
| NCU00153971 | Castanea pumila var. pumila |  |
| NCU00153969 | Castanea pumila var. pumila |  |
| NCU00153968 | Castanea pumila var. pumila |  |
| NCU00153352 | Castanea mollissima |  |
| NCU00153351 | Castanea henryi |  |
| NCU00153355 | Castanea seguinii |  |
| NCU00153354 | Castanea seguinii |  |
| NCU00153353 | Castanea mollissima |  |
| NCU00013728 | Castanea mollissima |  |
| NCU00153366 | Castanea dentata |  |
| NCU00153364 | Castanea dentata |  |
| NCU00153363 | Castanea dentata |  |
| NCU00153362 | Castanea dentata |  |
| NCU00153361 | Castanea dentata |  |
| NCU00153360 | Castanea dentata |  |
| NCU00153358 | Castanea dentata |  |
| NCU00153356 | Castanea dentata |  |
| NCU00153367 | Castanea dentata |  |
| NCU00153373 | Castanea pumila var. pumila |  |
| NCU00153371 | Castanea pumila var. pumila |  |
| NCU00153370 | Castanea pumila var. pumila |  |
| NCU00153369 | Castanea pumila var. pumila |  |
| NCU00157730 | Castanea mollissima |  |
| NCU00154862 | Castanea pumila var. pumila |  |
| NCU00154014 | Castanea pumila var. pumila |  |
| NCU00154015 | Castanea pumila var. pumila |  |
| NCU00154013 | Castanea pumila var. pumila |  |
| NCU00154016 | Castanea pumila var. pumila |  |
| NCU00154016 | Castanea pumila var. pumila |  |
| NCU00154017 | Castanea pumila var. pumila |  |
| NCU00154019 | Castanea pumila var. pumila |  |
| NCU00154020 | Castanea pumila var. pumila |  |
| NCU00154861 | Castanea pumila var. pumila |  |
| NCU00154011 | Castanea pumila var. pumila |  |
| NCU00154001 | Castanea pumila var. pumila |  |
| NCU00154002 | Castanea pumila var. pumila |  |
| NCU00154004 | Castanea pumila var. pumila |  |
| NCU00154005 | Castanea pumila var. pumila |  |
| NCU00154006 | Castanea pumila var. pumila |  |
| NCU00154008 | Castanea pumila var. pumila |  |
| NCU00154008 | Castanea pumila var. pumila |  |
| NCU00154009 | Castanea pumila var. pumila |  |
| NCU00154010 | Castanea pumila var. pumila |  |
| NCU00154000 | Castanea pumila var. pumila |  |
| NCU00153991 | Castanea pumila var. pumila |  |
| NCU00153993 | Castanea pumila var. pumila |  |
| NCU00153995 | Castanea pumila var. pumila |  |
| NCU00153996 | Castanea pumila var. pumila |  |
| NCU00153997 | Castanea pumila var. pumila |  |
| NCU00153999 | Castanea pumila var. pumila |  |
| NCU00153990 | Castanea pumila var. pumila |  |
| NCU00153986 | Castanea pumila var. pumila |  |
| NCU00153987 | Castanea pumila var. pumila |  |
| NCU00153989 | Castanea alabamensis | We observed only one stellate trichome in the axil of midrib and lateral vein on abaxial surface of leaf. Plant was sampled from Sumter Co., AL, a potential contact zone for different chinquapin varieties and may be a hybrid of C. alabamensis and C. p. var. pumila |
| NCU00153985 | Castanea pumila var. pumila |  |
| NCU00153977 | Castanea pumila var. pumila |  |
| NCU00153978 | Castanea pumila var. pumila |  |
| NCU00153980 | Castanea pumila var. pumila |  |
| NCU00153979 | Castanea pumila var. pumila |  |
| NCU00153982 | Castanea pumila var. pumila |  |
| NCU00153984 | Castanea pumila var. pumila |  |
| NCU00053435 | Castanea pumila var. ozarkensis |  |
| NCU00053446 | Castanea pumila var. ozarkensis |  |
| NCU00031434 | Castanea pumila var. ozarkensis |  |
| NCU00053444 | Castanea pumila |  |
| NCU00053441 | Castanea pumila var. ozarkensis |  |
| NCU00053442 | Castanea pumila var. ozarkensis |  |
| NCU00053439 | Castanea pumila var. ozarkensis |  |
| NCU00053440 | Castanea pumila var. ozarkensis |  |
| NCU00053447 | Castanea pumila var. ozarkensis |  |
| NCU00053452 | Castanea pumila var. ozarkensis |  |
| NCU00053451 | Castanea pumila |  |
| NCU00053450 | Castanea pumila var. ozarkensis |  |
| NCU00053449 | Castanea pumila var. ozarkensis |  |
| NCU00053453 | Castanea pumila var. ozarkensis |  |
| NCU00053460 | Castanea pumila var. ozarkensis |  |
| NCU00053459 | Castanea pumila var. ozarkensis |  |
| NCU00053457 | Castanea pumila var. ozarkensis |  |
| NCU00053418 | Castanea pumila var. ozarkensis |  |
| NCU00053456 | Castanea pumila var. ozarkensis |  |
| NCU00053461 | Castanea pumila var. ozarkensis |  |
| NCU00053463 | Castanea pumila var. ozarkensis |  |
| NCU00053462 | Castanea pumila var. ozarkensis |  |
| NCU00053436 | Castanea pumila var. pumila | Note the small bur (less than 2.5 cm diameter, including spines) and short leaf lengths (10 cm or less). |
| NCU00053437 | Castanea pumila | Mature bur diameter (2 cm), leaf lengths (11.5 cm in length, max.), and note on plant habit all suggest that plant could be either var. pumila or var. ozarkensis. |
| NCU00153150 | Castanea dentata |  |
| NCU00153148 | Castanea dentata |  |
| NCU00153169 | Castanea dentata |  |
| NCU00153166 | Castanea dentata |  |
| NCU00153165 | Castanea dentata |  |
| NCU00008701 | Castanea dentata |  |
| NCU00153164 | Castanea dentata |  |
| NCU00153162 | Castanea dentata |  |
| NCU00153161 | Castanea dentata |  |
| NCU00153170 | Castanea dentata |  |
| NCU00008702 | Castanea dentata |  |
| NCU00153177 | Castanea pumila var. pumila |  |
| NCU00031381 | Castanea pumila var. pumila |  |
| NCU00153059 | Castanea crenata | Note the cordate leaf bases, mix of stellate and simple abaxial trichomes, slender twig, and oblong leaf shape. |
| NCU00116435 | Castanea dentata |  |
| NCU00153147 | Castanea dentata |  |
| NCU00153146 | Castanea dentata |  |
| NCU00153145 | Castanea dentata |  |
| NCU00113377 | Castanea mollissima | Note cordate leaf bases, stipules wider than stem, crenate leaf margins. |
| NCU00154955 | Castanea pumila var. pumila |  |
| NCU00154953 | Castanea pumila var. pumila |  |
| NCU00154952 | Castanea pumila var. pumila |  |
| NCU00154949 | Castanea pumila var. pumila |  |
| NCU00154950 | Castanea pumila var. pumila |  |
| NCU00154947 | Castanea pumila var. pumila |  |
| NCU00154956 | Castanea pumila var. pumila |  |
| NCU00154966 | Castanea pumila var. pumila |  |
| NCU00154964 | Castanea pumila var. pumila |  |
| NCU00154962 | Castanea pumila var. pumila |  |
| NCU00154961 | Castanea pumila var. pumila |  |
| NCU00154960 | Castanea pumila var. pumila |  |
| NCU00154959 | Castanea pumila var. pumila |  |
| NCU00154957 | Castanea pumila var. pumila |  |
| NCU00154967 | Castanea pumila var. pumila |  |
| NCU00154973 | Castanea pumila var. pumila |  |
| NCU00154972 | Castanea pumila var. pumila |  |
| NCU00154971 | Castanea pumila var. pumila |  |
| NCU00154970 | Castanea pumila var. pumila |  |
| NCU00154969 | Castanea pumila var. pumila |  |
| NCU00154968 | Castanea pumila var. pumila |  |
| NCU00154974 | Castanea pumila var. pumila |  |
| NCU00155307 | Castanea pumila var. pumila |  |
| NCU00155306 | Castanea pumila var. pumila |  |
| NCU00155299 | Castanea pumila var. pumila |  |
| NCU00155298 | Castanea pumila var. pumila |  |
| NCU00155300 | Castanea pumila var. pumila |  |
| NCU00155301 | Castanea pumila var. pumila |  |
| NCU00155302 | Castanea pumila var. pumila |  |
| NCU00155217 | Castanea pumila var. pumila |  |
| NCU00155218 | Castanea pumila var. pumila |  |
| NCU00155213 | Castanea pumila var. pumila |  |

|  |  |
| --- | --- |
| NCU00155214 | Castanea pumila var. pumila |
| NCU00155215 | Castanea pumila var. pumila |
| NCU00155219 | Castanea pumila var. pumila |
| NCU00155209 | Castanea pumila var. pumila |
| NCU00155211 | Castanea pumila var. pumila |
| NCU00155210 | Castanea pumila var. pumila |
| NCU00155288 | Castanea pumila var. pumila |
| NCU00155285 | Castanea pumila var. pumila |
| NCU00155357 | Castanea dentata |
| NCU00155283 | Castanea pumila var. pumila |
| NCU00155355 | Castanea dentata |
| NCU00155354 | Castanea dentata |
| NCU00155349 | Castanea pumila var. ozarkensis |
| NCU00155348 | Castanea pumila var. ozarkensis |
| NCU00155352 | Castanea pumila var. ozarkensis |
| NCU00155346 | Castanea dentata |
| NCU00155104 | Castanea pumila var. pumila |
| NCU00155103 | Castanea pumila var. pumila |
| NCU00155102 | Castanea pumila var. pumila |
| NCU00155101 | Castanea pumila var. pumila |
| NCU00154979 | Castanea pumila var. pumila |
| NCU00154980 | Castanea pumila var. pumila |
| NCU00154977 | Castanea pumila var. pumila |
| NCU00154976 | Castanea pumila var. pumila |
| NCU00155105 | Castanea pumila var. pumila |
| NCU00154975 | Castanea pumila var. pumila |
| NCU00155114 | Castanea pumila var. pumila |
| NCU00155113 | Castanea pumila var. pumila |
| NCU00155112 | Castanea pumila var. pumila |
| NCU00155111 | Castanea pumila var. pumila |
| NCU00155109 | Castanea pumila var. pumila |
| NCU00155108 | Castanea pumila var. pumila |
| NCU00155107 | Castanea pumila var. pumila |
| NCU00155106 | Castanea pumila var. pumila |
| NCU00155115 | Castanea pumila var. pumila |
| NCU00155120 | Castanea pumila var. pumila |
| NCU00155119 | Castanea pumila var. pumila |
| NCU00155118 | Castanea pumila var. pumila |
| NCU00155117 | Castanea pumila var. pumila |
| NCU00155116 | Castanea pumila var. pumila |
| NCU00157719 | Castanea pumila var. pumila |
| NCU00157720 | Castanea pumila var. pumila |
| NCU00157721 | Castanea pumila var. pumila |
| NCU00157723 | Castanea pumila var. pumila |
| NCU00157724 | Castanea pumila var. pumila |
| NCU00157725 | Castanea pumila var. pumila |
| NCU00157726 | Castanea pumila var. pumila |
| NCU00155289 | Castanea pumila var. pumila |
| NCU00155297 | Castanea pumila var. pumila |
| NCU00155296 | Castanea pumila var. pumila |
| NCU00155295 | Castanea pumila var. pumila |
| NCU00155294 | Castanea pumila var. pumila |
| NCU00155293 | Castanea pumila var. pumila |
| NCU00155292 | Castanea pumila var. pumila |
| NCU00155291 | Castanea pumila var. pumila |
| NCU00155220 | Castanea pumila var. pumila |
| NCU00155281 | Castanea pumila var. pumila |
| NCU00155284 | Castanea pumila var. pumila |
| NCU00155318 | Castanea pumila var. pumila |
| NCU00155317 | Castanea pumila var. pumila |
| NCU00155316 | Castanea pumila var. pumila |
| NCU00092135 | Castanea pumila var. pumila |
| NCU00155320 | Castanea pumila var. pumila |
| NCU00155310 | Castanea pumila var. pumila |
| NCU00155309 | Castanea pumila var. pumila |
| NCU00155308 | Castanea pumila var. pumila |
| NCU00155315 | Castanea pumila var. pumila |
| NCU00089777 | Castanea pumila var. pumila |
| NCU00155311 | Castanea pumila var. pumila |
| NCU00155314 | Castanea pumila var. pumila |
| NCU00155304 | Castanea pumila var. pumila |
| NCU00155303 | Castanea pumila var. pumila |
| NCU00135543 | Castanea pumila var. pumila |
| NCU00155319 | Castanea pumila var. pumila |
| NCU00155321 | Castanea dentata |
| NCU00135547 | Castanea crenata |
| NCU00155322 | Castanea dentata |
| NCU00155170 | Castanea pumila var. pumila |
| NCU00155197 | Castanea pumila var. pumila |
| NCU00155196 | Castanea pumila var. pumila |
| NCU00155195 | Castanea pumila var. pumila |
| NCU00155194 | Castanea pumila var. pumila |
| NCU00155199 | Castanea pumila var. pumila |
| NCU00155178 | Castanea pumila var. pumila |
| NCU00155177 | Castanea pumila var. pumila |
| NCU00155181 | Castanea pumila var. pumila |
| NCU00155358 | Castanea pumila var. pumila |
| NCU00155361 | Castanea pumila var. pumila |
| NCU00155365 | Castanea pumila var. pumila |
| NCU00155364 | Castanea pumila var. pumila |
| NCU00155363 | Castanea pumila var. pumila |
| NCU00155370 | Castanea pumila var. pumila |
| NCU00155359 | Castanea pumila var. pumila |
| NCU00155381 | Castanea crenata |
| NCU00155380 | Castanea crenata |
| NCU00155379 | Castanea crenata |
| NCU00155378 | Castanea crenata |
| NCU00155377 | Castanea crenata |
| NCU00155376 | Castanea crenata |
| NCU00155323 | Castanea dentata |
| NCU00155330 | Castanea dentata |
| NCU00155325 | Castanea dentata |
| NCU00155327 | Castanea dentata |
| NCU00155324 | Castanea dentata |
| NCU00155334 | Castanea dentata |
| NCU00155335 | Castanea dentata |
| NCU00155336 | Castanea dentata |
| NCU00155329 | Castanea dentata |
| NCU00155326 | Castanea dentata |
| NCU00155347 | Castanea dentata |
| NCU00155332 | Castanea dentata |
| NCU00155331 | Castanea dentata |
| NCU00155338 | Castanea dentata |
| NCU00155333 | Castanea dentata |
| NCU00155339 | Castanea dentata |
| NCU00155342 | Castanea dentata |
| NCU00155341 | Castanea dentata |
| NCU00155340 | Castanea dentata |
| NCU00155345 | Castanea dentata |
| NCU00155344 | Castanea dentata |
| NCU00155343 | Castanea dentata |
| NCU00155353 | Castanea pumila var. ozarkensis |
| NCU00155350 | Castanea pumila var. ozarkensis |
| NCU00154917 | Castanea pumila var. pumila |
| NCU00154916 | Castanea pumila var. pumila |
| NCU00154915 | Castanea pumila var. pumila |
| NCU00154914 | Castanea pumila var. pumila |
| NCU00135545 | Castanea pumila var. pumila |
| NCU00154934 | Castanea pumila var. pumila |
| NCU00154933 | Castanea pumila var. pumila |
| NCU00154931 | Castanea pumila var. pumila |
| NCU00154930 | Castanea pumila var. pumila |
| NCU00154929 | Castanea pumila var. pumila |
| NCU00154928 | Castanea pumila var. pumila |
| NCU00154927 | Castanea pumila var. pumila |
| NCU00154926 | Castanea pumila var. pumila |  |
| NCU00154925 | Castanea pumila var. pumila |  |
| NCU00135546 | Castanea pumila var. pumila |  |
| NCU00154924 | Castanea pumila var. pumila |  |
| NCU00154935 | Castanea pumila var. pumila |  |
| NCU00154945 | Castanea pumila var. pumila |  |
| NCU00154943 | Castanea pumila var. pumila |  |
| NCU00154942 | Castanea pumila var. pumila |  |
| NCU00154941 | Castanea pumila var. pumila |  |
| NCU00154940 | Castanea pumila var. pumila |  |
| NCU00154939 | Castanea pumila var. pumila |  |
| NCU00154939 | Castanea pumila var. pumila |  |
| NCU00154938 | Castanea pumila var. pumila |  |
| NCU00154936 | Castanea pumila var. pumila |  |
| NCU00154946 | Castanea pumila var. pumila |  |
| NCU00154919 | Castanea pumila var. pumila |  |
| NCU00154920 | Castanea pumila var. pumila |  |
| NCU00154922 | Castanea pumila var. pumila |  |
| NCU00154923 | Castanea pumila var. pumila |  |
| NCU00155162 | Castanea pumila var. pumila |  |
| NCU00155161 | Castanea pumila var. pumila |  |
| NCU00155160 | Castanea pumila var. pumila |  |
| NCU00155159 | Castanea pumila var. pumila |  |
| NCU00155158 | Castanea pumila var. pumila |  |
| NCU00155157 | Castanea pumila var. pumila |  |
| NCU00155156 | Castanea pumila var. pumila |  |
| NCU00155134 | Castanea pumila var. pumila |  |
| NCU00155121 | Castanea pumila var. pumila |  |
| NCU00155127 | Castanea pumila var. pumila |  |
| NCU00155123 | Castanea pumila var. pumila |  |
| NCU00155124 | Castanea pumila var. pumila |  |
| NCU00155125 | Castanea pumila var. pumila |  |
| NCU00155126 | Castanea pumila var. pumila |  |
| NCU00155154 | Castanea pumila var. pumila |  |
| NCU00155163 | Castanea pumila var. pumila |  |
| NCU00155144 | Castanea pumila var. pumila |  |
| NCU00155143 | Castanea pumila var. pumila |  |
| NCU00155153 | Castanea pumila var. pumila |  |
| NCU00155146 | Castanea pumila var. pumila |  |
| NCU00155374 | Castanea sativa | Note the long petioles, stout twigs,<br>small lenticels, elongate leaves,<br>and large buds. |
| NCU00155372 | Castanea mollissima |  |
| NCU00155373 | Castanea sativa |  |
| NCU00155368 | Castanea pumila var. pumila |  |
| NCU00155367 | Castanea pumila var. pumila |  |
| NCU00155366 | Castanea pumila var. pumila |  |
| NCU00155371 | Castanea pumila var. pumila |  |
| NCU00155392 | Castanea seguinii | Burs at multiple nodes along the stem are evidence of the continuous flowering typical of Castanea seguinii. |
| NCU00155396 | Castanea sativa |  |
| NCU00155382 | Castanea crenata |  |
| NCU00155394 | Castanea sativa | We think this is from Manali, Himachal Pradesh, India |
| NCU00155393 | Castanea sativa |  |
| NCU00155384 | Castanea henryi |  |
| NCU00155383 | Castanea seguinii | Burs at multiple nodes along the stem are evidence of the continuous flowering typical of Castanea seguinii. |
| NCU00155397 | Castanea sativa |  |
| NCU00155391 | Castanea sativa |  |
| NCU00155388 | Castanea sativa |  |
| NCU00155390 | Castanea sativa |  |
| NCU00155387 | Castanea sativa |  |
| NCU00155386 | Castanea sativa |  |
| NCU00155385 | Castanea henryi |  |
| NCU00155180 | Castanea pumila var. pumila |  |
| NCU00155182 | Castanea pumila var. pumila |  |
| NCU00155185 | Castanea pumila var. pumila |  |
| NCU00155190 | Castanea pumila var. pumila |  |
| NCU00155193 | Castanea pumila var. pumila |  |
| NCU00155192 | Castanea pumila var. pumila |  |
| NCU00155202 | Castanea pumila var. pumila |  |
| NCU00155201 | Castanea pumila var. pumila |  |
| NCU00155200 | Castanea pumila var. pumila |  |
| NCU00155208 | Castanea pumila var. pumila |  |
| NCU00155207 | Castanea pumila var. pumila |  |
| NCU00155206 | Castanea pumila var. pumila |  |
| NCU00155205 | Castanea pumila var. pumila |  |
| NCU00155204 | Castanea pumila var. pumila |  |
| NCU00155171 | Castanea pumila var. pumila |  |
| NCU00155176 | Castanea pumila var. pumila |  |
| NCU00155175 | Castanea pumila var. pumila |  |
| NCU00155174 | Castanea pumila var. pumila |  |
| NCU00146417 | Castanea pumila var. pumila |  |
| NCU00027810 | Castanea pumila var. pumila |  |
| NCU00155168 | Castanea pumila var. pumila |  |
| NCU00155167 | Castanea pumila var. pumila |  |

|  |  |
| --- | --- |
| NCU00155166 | Castanea pumila var. pumila |
| NCU00155165 | Castanea pumila var. pumila |
| NCU00155164 | Castanea pumila var. pumila |
| NCU00155172 | Castanea pumila var. pumila |
| NCU00155189 | Castanea pumila var. pumila |
| NCU00155188 | Castanea pumila var. pumila |
| NCU00155187 | Castanea pumila var. pumila |
| NCU00155145 | Castanea pumila var. pumila |
| NCU00155150 | Castanea pumila var. pumila |
| NCU00155148 | Castanea pumila var. pumila |
| NCU00155149 | Castanea pumila var. pumila |
| NCU00155152 | Castanea pumila var. pumila |
| NCU00155135 | Castanea pumila var. pumila |
| NCU00155137 | Castanea pumila var. pumila |
| NCU00155136 | Castanea pumila var. pumila |
| NCU00155141 | Castanea pumila var. pumila |
| NCU00155140 | Castanea pumila var. pumila |
| NCU00155139 | Castanea pumila var. pumila |
| NCU00155131 | Castanea pumila var. pumila |
| NCU00155130 | Castanea pumila var. pumila |
| NCU00155133 | Castanea pumila var. pumila |
| NCU00155128 | Castanea pumila var. pumila |
| NCU00155389 | Castanea sativa |
| NCU00155129 | Castanea pumila var. pumila |

**Appendix S4.**
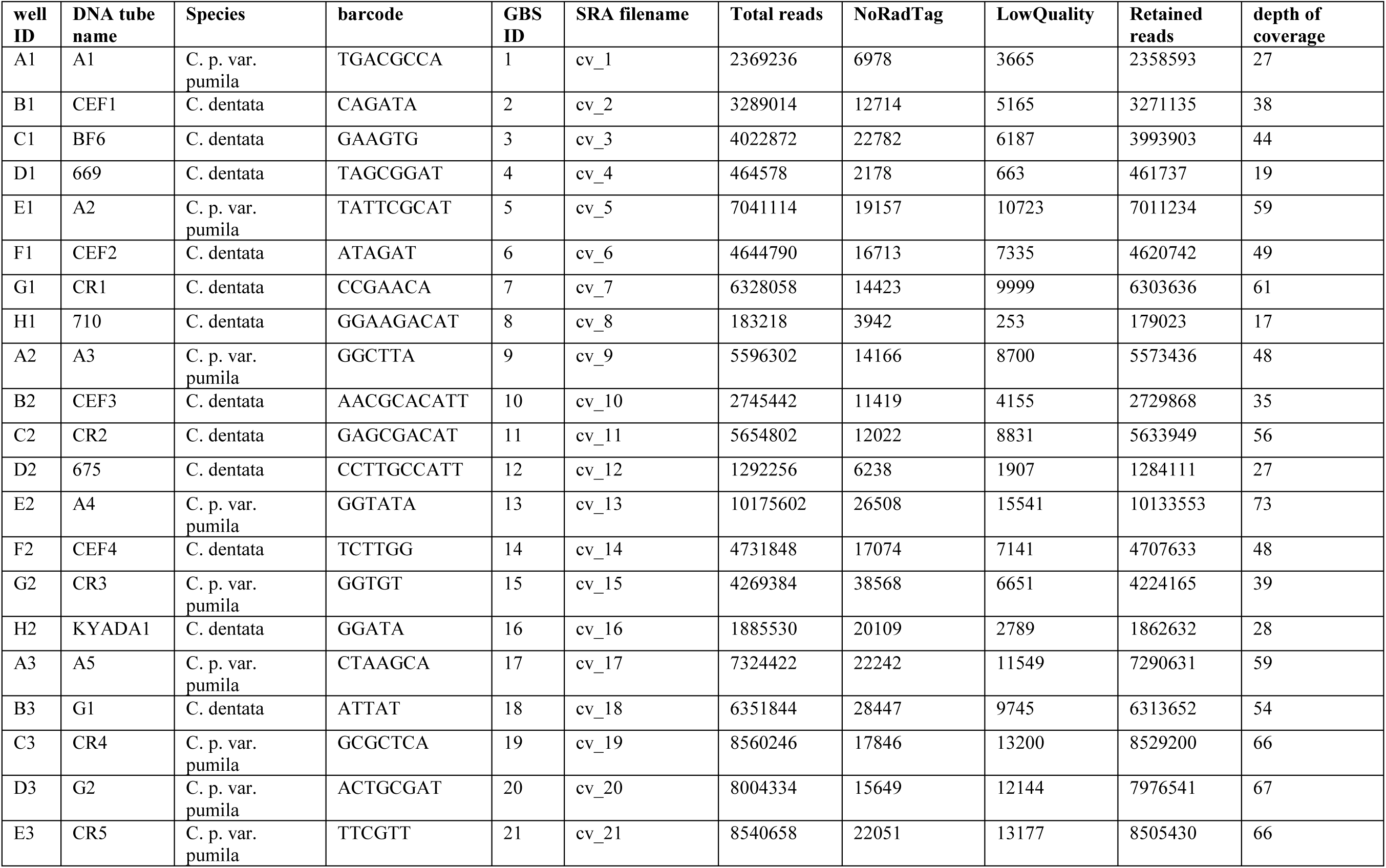
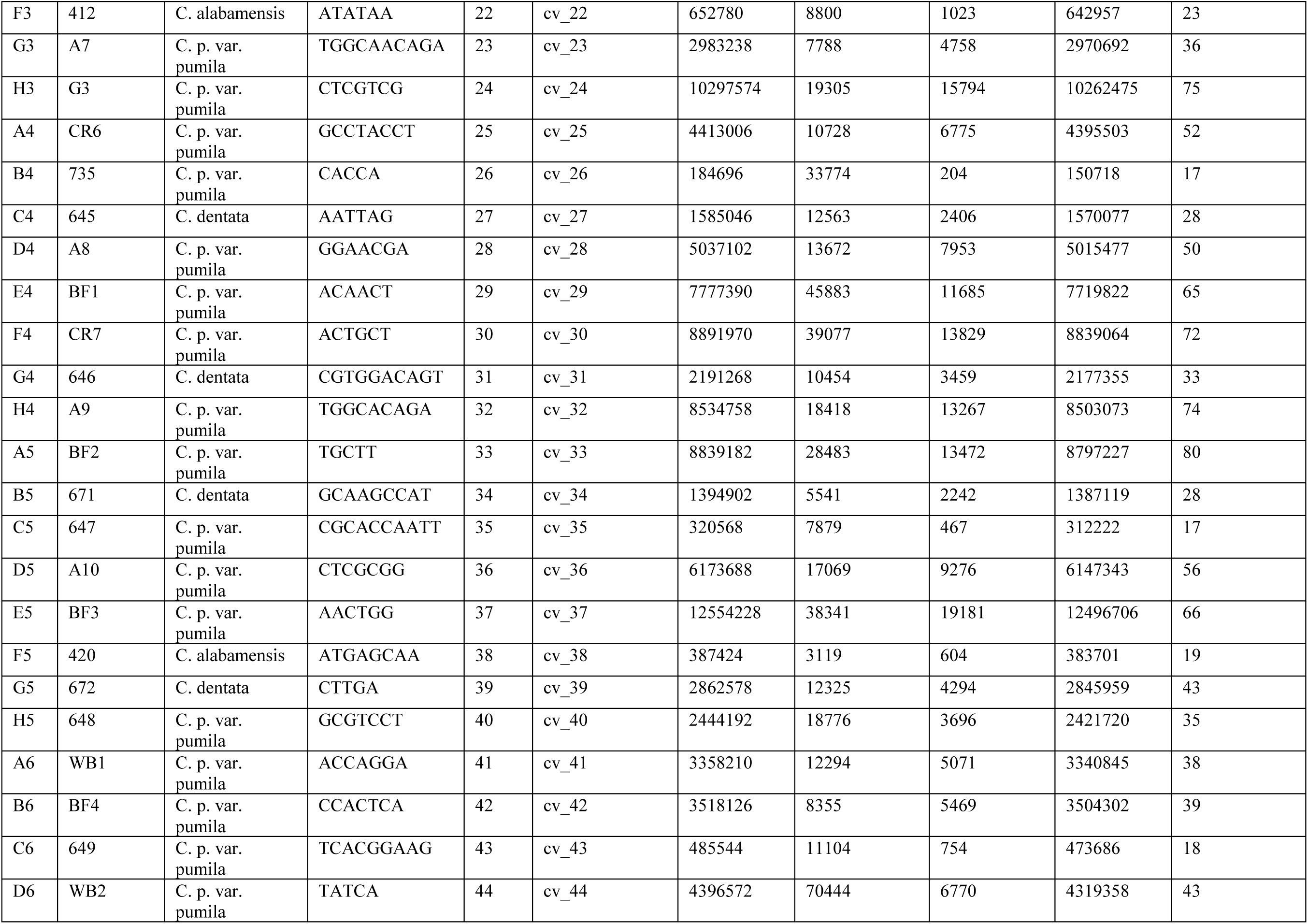
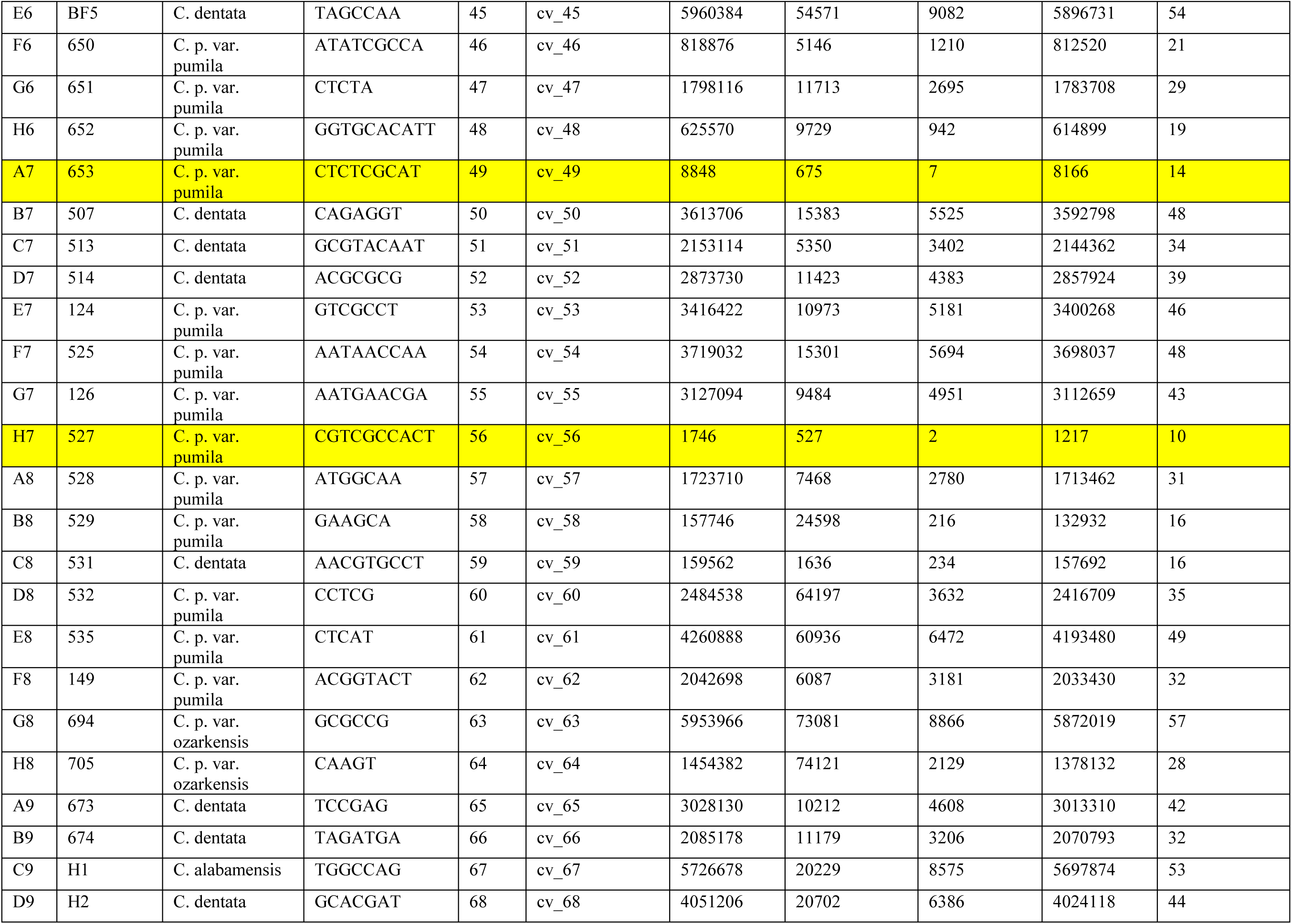
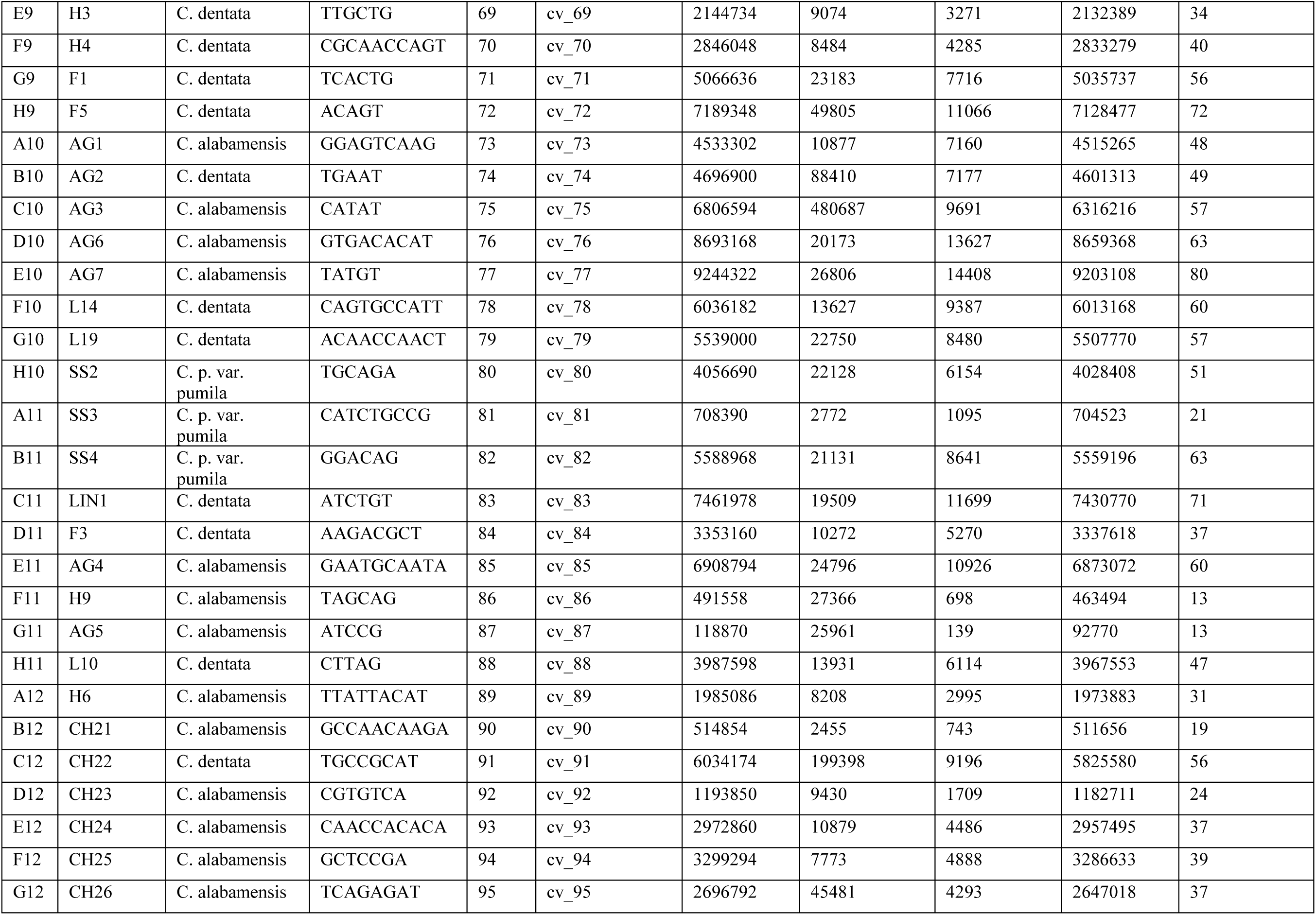
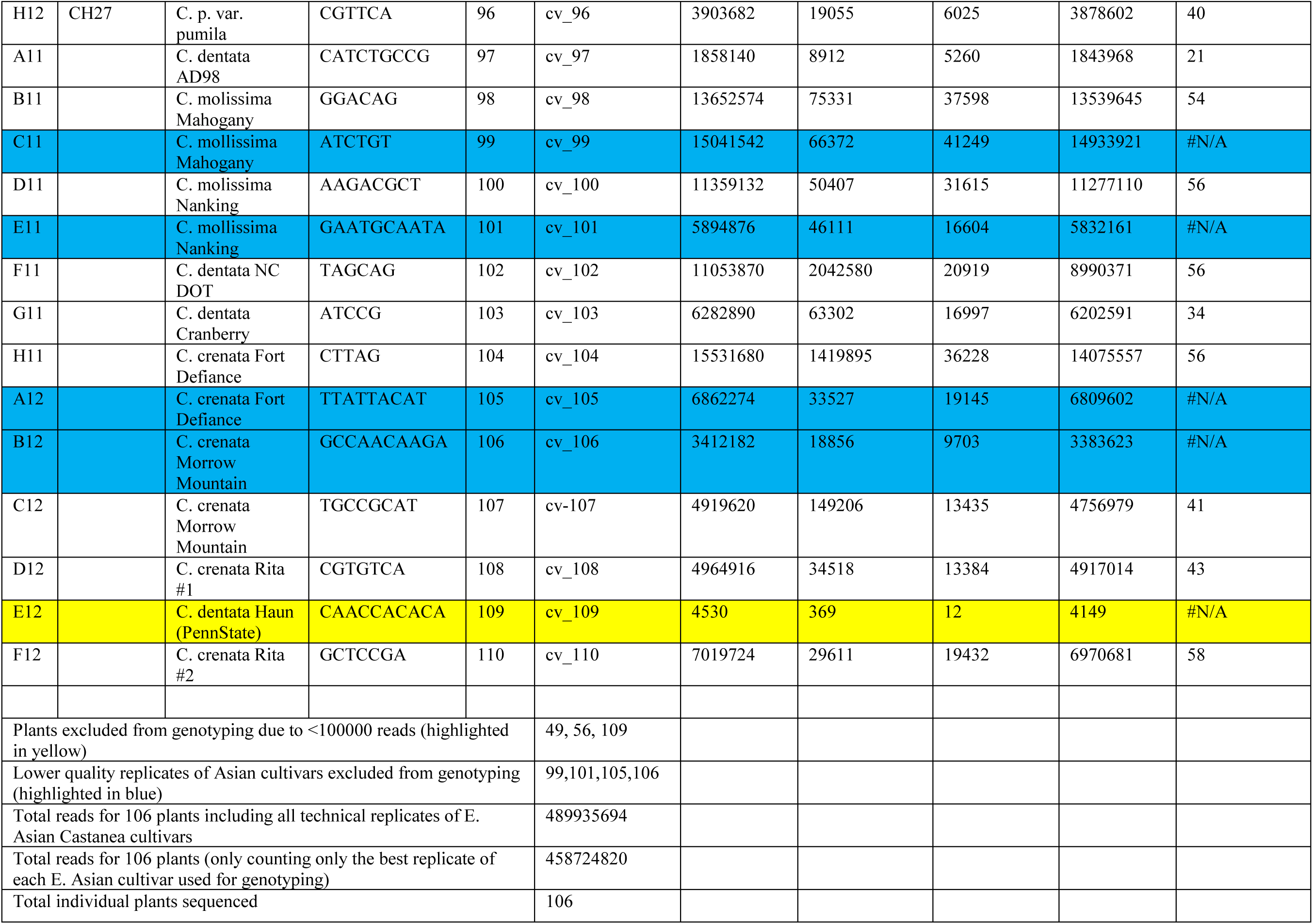
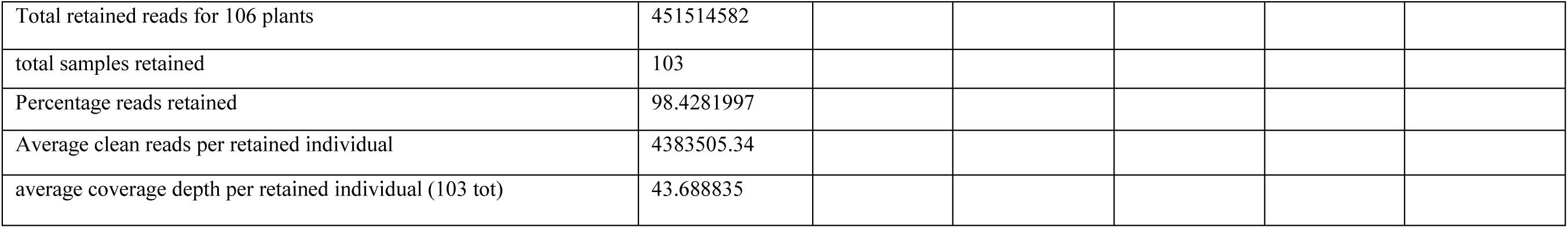
Illumina sequencing summary. Samples highlighted in yellow were not used for genotyping due to a low number (<100,000) of total reads. Samples highlighted in blue are technical replicates of *C. mollissima* and *C. crenata* cultivars that were not genotyped because they produced the fewest retained reads of two technical replicates per cultivar.

**Appendix S5.** Summary of datasets exported by Stacks 1.45.

| Dataset name | File name on GitHub | Taxa included | Total individuals | Total SNPs |
| --- | --- | --- | --- | --- |
| NAC+EAC | batch_1.vcf | *C. alabamensis*, *C. dentata*, *C. pumila* var. *pumila*, *C. pumila* var. *ozarkensis*, *C. crenata*, *C. mollissima* | 103 | 103,616 |
| NAC | out98copy.vcf | *C. alabamensis*, *C. dentata*, *C. pumila* var. *pumila*, *C. pumila* var. *ozarkensis* | 96 | 190,656 |
| NAC+EAC583 | allsamples_popcodes.txt | *C. alabamensis*, *C. dentata*, *C. pumila* var. *pumila*, *C. pumila* var. *ozarkensis*, *C. crenata*, *C. mollissima* | 103 | 583 |

**Appendix S6.**
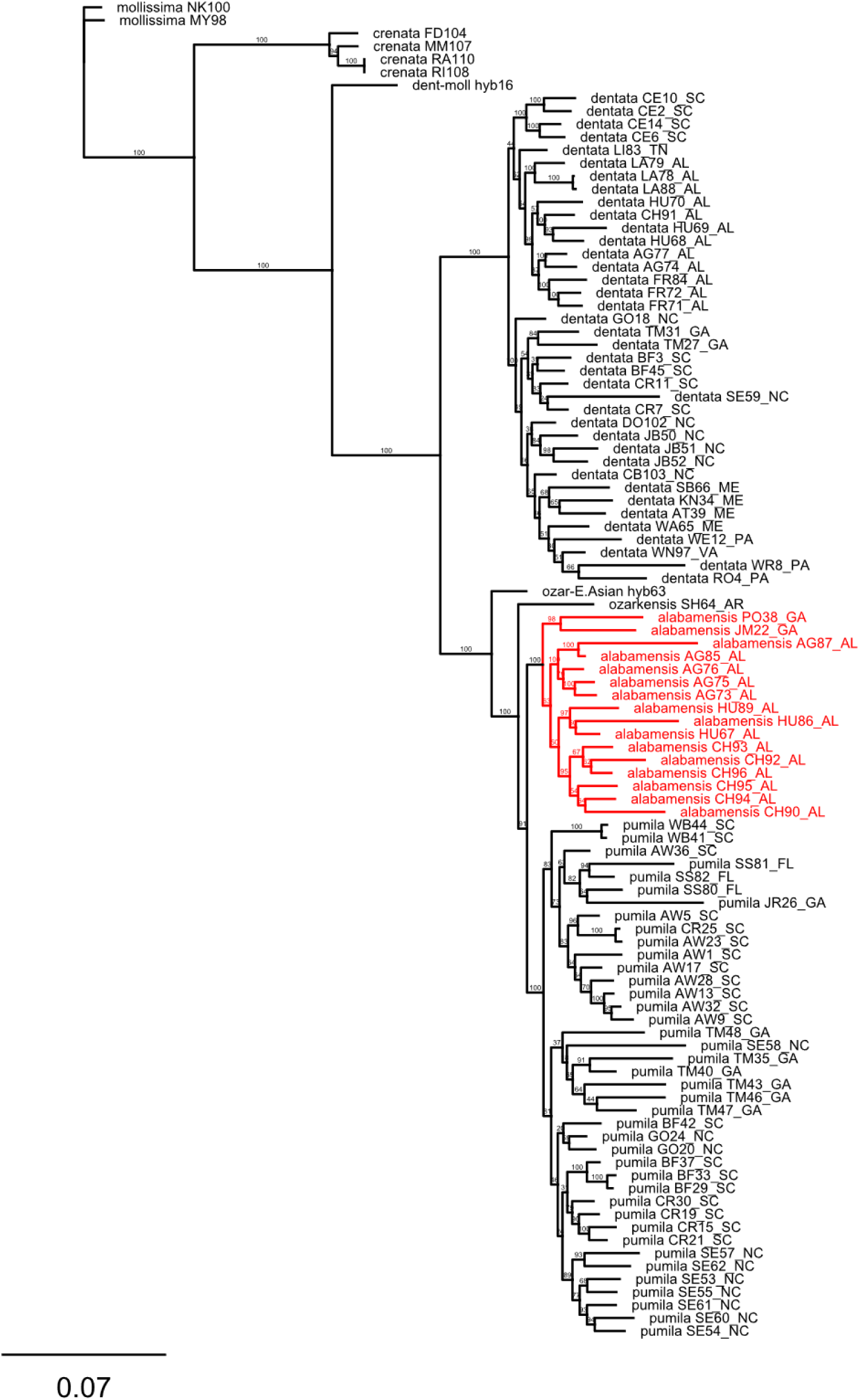
Maximum likelihood phylogenetic tree showing identities of individual samples at branch tips. Samples representing *C. alabamensis* are highlighted in red. Numbers at nodes indicate bootstrap support. Scale bar is proportional to 0.07 substitutions/site.

## Appendix S7.

Weir and Cockerham’s *Fst* between taxa pairs, summary of STRUCTURE analyses, and list of principal components.

**Table S7a.** Weir and Cockerham’s Fst output from SNPrelate. Output: “Fst” = weighted Fst estimate

|  | mollissima | crenata | dentata | pumila | ozarkensis | alabamensis |
| --- | --- | --- | --- | --- | --- | --- |
| mollissima |  | 0.5271483 | 0.6433073 | 0.6378966 | 0.7069509 | 0.6633516 |
| crenata | 0.5271483 |  | 0.6409639 | 0.6345136 | 0.6765049 | 0.6569612 |
| dentata | 0.6433073 | 0.6409639 |  | 0.3846462 | 0.4182757 | 0.3985978 |
| pumila | 0.6378966 | 0.6345136 | 0.3846462 |  | 0.1668146 | 0.1078042 |
| ozarkensis | 0.7069509 | 0.6765049 | 0.4182757 | 0.1668146 |  | 0.204663 |
| alabamensis | 0.6633516 | 0.6569612 | 0.3985978 | 0.1078042 | 0.204663 |  |

**Table S7b.**
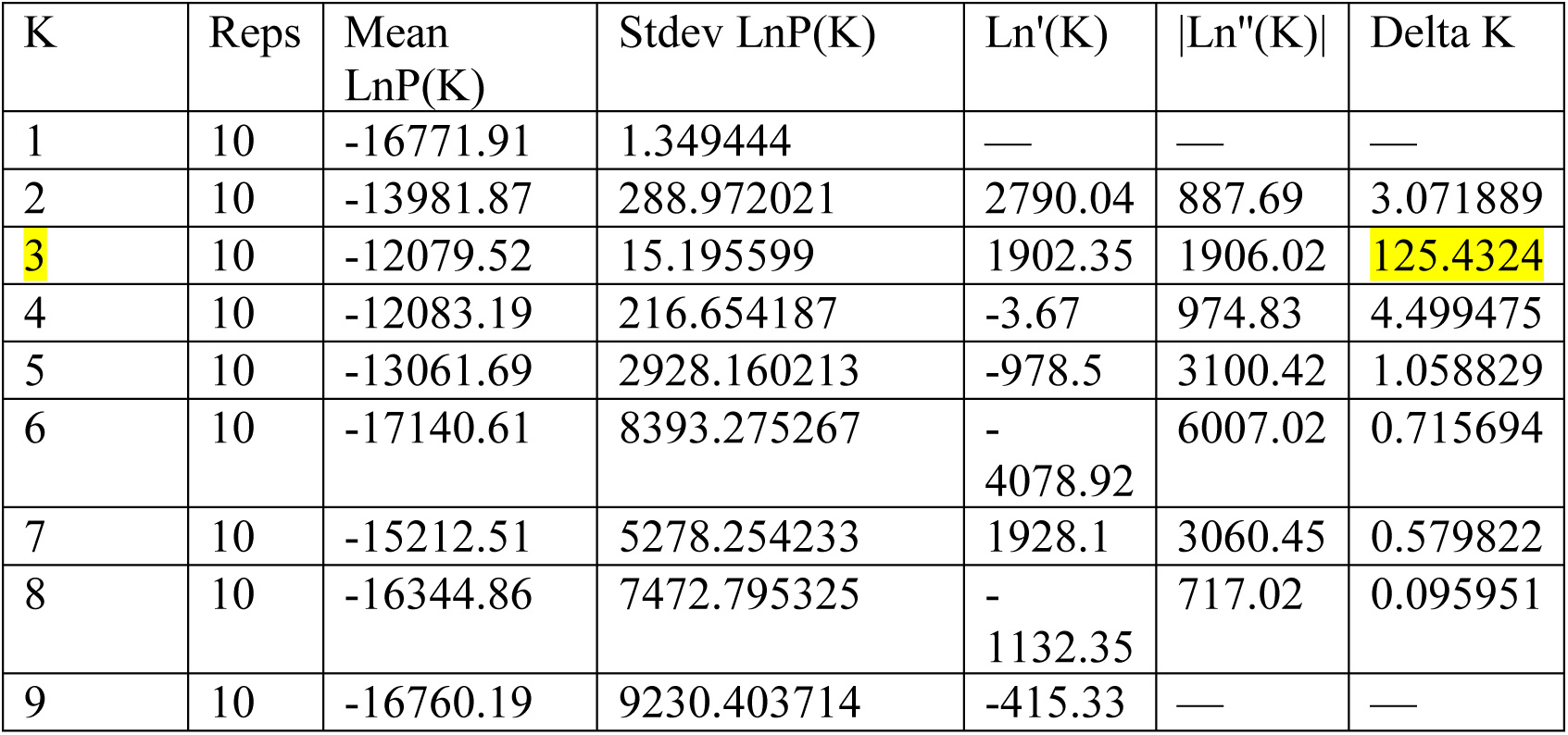
Output of summary statistics from *Structure* analysis of *C. dentata*, *C. pumila sensu lato*, *C. mollissima*, and *C. crenata*. Statistics were calculated using the program STRUCTURE HARVESTER (Earl and vonHoldt 2012). The number of clusters present in the dataset (*K* = 3) was determined using the method of Evanno et al. (2005). *K* = 3 and associated Δ*K* value are highlighted in yellow.

**Table S7c.** Output of summary statistics from Structure analysis of chinquapin samples. Statistics were calculated using the program STRUCTURE HARVESTER (Earl and vonHoldt 2012). The number of clusters present in the dataset (*K* = 2) was determined using the method of Evanno et al. (2005). *K* = 2 and associated Δ*K* value are highlighted in yellow.

| K | Reps | Mean<br>LnP(K) | Stdev<br>LnP(K) | Ln'(K) | \|Ln''(K)\| | Delta K |
| --- | --- | --- | --- | --- | --- | --- |
| 1 | 10 | -6426.61 | 0.479467 | — | — | — |
| 2 | 10 | -6041.17 | 2.833941 | 385.44 | 296.32 | 104.5611 |
| 3 | 10 | -5952.05 | 137.78048 | 89.12 | 31.65 | 0.229713 |
| 4 | 10 | -5831.28 | 39.063219 | 120.77 | 449.57 | 11.50878 |
| 5 | 10 | -6160.08 | 1208.25738 | -328.8 | 644.53 | 0.533438 |
| 6 | 10 | -5844.35 | 182.341421 | 315.73 | 265.61 | 1.456663 |
| 7 | 10 | -5794.23 | 223.654088 | 50.12 | 222.98 | 0.996986 |
| 8 | 10 | -5967.09 | 647.43212 | -172.86 | 301.03 | 0.46496 |
| 9 | 10 | -5838.92 | 294.650689 | 128.17 | 423.32 | 1.436684 |
| 10 | 10 | -6134.07 | 1064.66472 | -295.15 | 525.14 | 0.493244 |
| 11 | 10 | -5904.08 | 372.702293 | 229.99 | 285.78 | 0.766778 |
| 12 | 10 | -5959.87 | 336.013763 | -55.79 | 154.31 | 0.459237 |
| 13 | 10 | -5861.35 | 275.391916 | 98.52 | 128.35 | 0.466063 |
| 14 | 10 | -5891.18 | 306.478028 | -29.83 | 93.25 | 0.304263 |
| 15 | 10 | -6014.26 | 285.24903 | -123.08 | 286.41 | 1.00407 |
| 16 | 10 | -5850.93 | 306.721405 | 163.33 | — | — |

**Table S7d.** Proportions of total variance explained by the 32 principal components identified from principal components analysis of North American *Castanea* samples. The 32 principal components combined explain 56.5% of the total variance.

| Principal components | Proportion variance |
| --- | --- |
| PC1 | 0.10957356 |
| PC2 | 0.02748213 |
| PC3 | 0.02287274 |
| PC4 | 0.0209522 |
| PC5 | 0.02088753 |
| PC6 | 0.01773292 |
| PC7 | 0.01741902 |
| PC8 | 0.01648643 |
| PC9 | 0.01636512 |
| PC10 | 0.01508126 |
| PC11 | 0.01485904 |
| PC12 | 0.01454827 |
| PC13 | 0.01430424 |
| PC14 | 0.01416125 |
| PC15 | 0.01406701 |
| PC16 | 0.01339484 |
| PC17 | 0.01296989 |
| PC18 | 0.01289406 |
| PC19 | 0.0127412 |
| PC20 | 0.01270266 |
| PC21 | 0.01257472 |
| PC22 | 0.01237297 |
| PC23 | 0.01229619 |
| PC24 | 0.01221846 |
| PC25 | 0.01204935 |
| PC26 | 0.01201129 |
| PC27 | 0.01188767 |
| PC28 | 0.01183929 |
| PC29 | 0.01173212 |
| PC30 | 0.0116869 |
| PC31 | 0.01160848 |
| PC32 | 0.01147692 |

## Notes

https://www.ncbi.nlm.nih.gov/sra/?term=PRJNA541592

https://github.com/MTPerkins/Nonhybrid_origin_of_Castanea_alabamensis

